# RCDome: A Multi-Database Human Gene Atlas of Regulated Cell Death

**DOI:** 10.64898/2026.10.29.761548

**Authors:** Emanuell Rodrigues de Souza, Victor dos Santos Lopes, Higor Almeida Cordeiro Nogueira, Enrique Medina-Acosta

## Abstract

**Introduction:** Regulated cell death (RCD) governs tissue homeostasis, development, and disease, and is executed or regulated by specific genes. Yet for the researcher asking which genes execute or regulate a given RCD form, the answer is rarely simple: gene lists for the same form rarely agree across databases, many associations carry no traceable source, and most have never been systematically checked against primary literature. We built RCDome to answer the most basic question a researcher can ask: where did this gene-RCD association come from, and what is the evidence?

**Methods:** RCDome is a multi-database human gene atlas built through a staged architecture integrating multi-source compilation, literature validation and evidence recovery, mechanistic classification, human-governed ontological adjudication, and web deployment. Five complementary evidence channels—PubMed Boolean search, curated databases, AI-assisted screening, manual curation, and Elicit systematic review—were integrated from 20 public databases into a unified matrix. Sixty-two original name strings were canonicalized into 53 RCD-related entities with curator-adjudicated operational definitions and ontological levels. Gene-RCD associations were systematically evaluated through an independent DeepSeek two-call LLM validation, Europe PMC full-text recovery and deep evidence extraction, an independent Consensus literature-search channel, and GPT-4o arbitration of disagreements. Source-grounding controls rejected extracted evidence not found verbatim in the source document, while the governance contract was enforced as executable code.

**Results:** The compendium holds 22,473 association rows spanning 9,265 unique HGNC-validated genes (21,568 unique gene-RCD pairs). Evidence-first adjudication resolved 5,138 pairs (23.8%) as mechanistically involved; across the 4,775 role-resolved pairs, 225 (4.7%) are executors, 4,455 (93.3%) are regulators, and 95 are markers, revealing a small death-executing core embedded in a vast regulatory network. Under a separate seven-class mechanistic criterion applied to all 22,473 rows, only 3,063 (13.6%) were mechanistically confirmed and 286 (1.3%) were database-only with no verifiable paper trail.

**Conclusion:** RCDome makes the RCD evidence landscape measurable and traceable, showing that AI-assisted curation can be verifiable, human-governed, and explicit about uncertainty. It is delivered as a Shiny web application that lets any researcher query the atlas and inspect the evidence behind each association.

**Availability:** RCDome is freely available at https://rcdome.org/ (source code and data at https://github.com/BioCancerInformatics/RCDome).

## INTRODUCTION

Regulated cell death (RCD) governs tissue homeostasis, development, and disease pathogenesis through genetically encoded entities that eliminate superfluous, damaged, or dangerous cells (Galluzzi et al., 2018). The Nomenclature Committee on Cell Death (NCCD) established the foundational consensus framework with molecularly oriented operational definitions for 12 RCD modalities (intrinsic apoptosis, extrinsic apoptosis, MPT-driven necrosis, necroptosis, ferroptosis, pyroptosis, parthanatos, entotic cell death, NETotic cell death, lysosome-dependent cell death, autophagy-dependent cell death, and immunogenic cell death) and three non-lethal processes (cellular senescence, mitotic catastrophe, and terminal differentiation) (Galluzzi et al., 2018). The NCCD’s operational-definition list totals 20 entities: the 12 modalities and three non-lethal processes listed above, plus anoikis, autosis, efferocytosis, mitotic death, and programmed cell death, all retained verbatim in RCDome. Building on this framework, our group compiled gene-level associations for 25 RCD forms and provided tentative operational definitions for forms not covered in the NCCD consensus (Rodrigues de Souza et al., 2025, Rodrigues de Souza et al., 2026). Since that compilation, the RCD landscape has continued to expand, with the recognition of necrosis by sodium overload (NECSO), mitoxyperilysis, triaptosis, spectosis, floatptosis, lysozincrosis, and metalloptosis (forms that had emerged in the literature before this compilation), alongside the proliferation of specialized databases; the field now encompasses dozens of RCD forms. Each RCD form is claimed to be executed or regulated by specific genes, and these gene-RCD relationships are the molecular basis on which each death program is defined and experimentally interrogated. However, the operational reality for researchers has not kept pace: gene-RCD associations are recorded under heterogeneous naming conventions, supported by evidence of widely varying quality, and scattered across databases that were never designed to interoperate.

Specialized databases such as FerrDB v3 (22 RCD forms, 3,481 genes) (Zhou et al., 2026) and RCDdb (Wang et al., 2024), together with our prior compilation (25 RCD forms, 5,913 genes) (Rodrigues de Souza et al., 2025), represent major advances in RCD knowledge organization. FerrDB v3 is primarily a catalog of RCD regulators and markers extracted from >20,000 publications, designed for form-specific browsing rather than cross-form integration. RCDdb provides explicit PMIDs for a valuable subset of its entries but is limited to 15 RCD forms. Our prior compilation covered 25 RCD forms but was derived from a single literature-search strategy. Critically, none of these resources was designed to integrate all known RCD forms across the governed source universe, normalize heterogeneous naming conventions, or independently validate every association against the published literature. A multi-database compilation that integrates these and other resources is therefore necessary for three reasons: (i) no single database covers the full RCD landscape; (ii) gene-RCD associations reported by different databases are not directly comparable due to heterogeneous naming conventions (e.g., FerrDB v3 uses “Entotic-cell-death,” LncRCD uses “Entotic cell death,” while RCDdb and iPCD use “Entosis”; our prior compilation similarly used “Autophagy” as an umbrella term without differentiating autophagy-dependent cell death); and (iii) even those well-curated databases lack per-gene literature traceability, making it impossible to verify the provenance of individual associations. These gaps motivate the development of the platform we describe below.

The underutilization of published biological data is a well-documented structural problem: a large-scale survey of 454,200 public omics datasets found that only 2.7% had undergone documented reanalysis (Perez-Riverol et al., 2019). This pattern extends to curated gene-function databases, where the vast majority of recorded associations remain unvalidated against primary literature. Combined with recent calls for rigorous standards, validation, and human oversight in AI-assisted data reanalysis (Ghaderi et al., 2026), this creates both an opportunity and an obligation: to build platforms that do not merely aggregate existing annotations, but actively verify, contextualize, and trace every association to its evidence source. The RCD field presents a particularly acute case for such a platform, given the proliferation of specialized databases, the heterogeneity of RCD nomenclature, and the accelerating pace of discovery of new cell death forms.

Yet even with these few dedicated RCD resources, and despite the broader recognition that published data remains critically underutilized, finding which genes are linked to which RCD forms remains unnecessarily difficult. The gene lists for a given RCD form rarely agree from one database to the next. Some associations appear without a single PubMed Identifier (PMID) or PubMed Central Identifier (PMCID) attached, making it impossible to trace the claim back to a source article or to determine whether the link is real or an artifact of database propagation. Even when a source is traceable, it is rarely clear what kind of association it describes: is the gene an executioner, a regulator, or merely mentioned alongside the RCD term? We therefore set out to build a resource that could answer the most basic question a researcher should be able to ask in RCD biology: “Where did this gene-RCD association come from, and what is the evidence?”

Here we present RCDome, a multi-database knowledge platform that compiles, normalizes, and checks every gene-level association against the published record, extracting the evidence that supports, or fails to support, each database claim. The platform is named RCDome to evoke the architectural dome: a structure that, by definition, provides comprehensive overhead coverage, unified enclosure, and a single vantage point from which to survey an entire space, in this case, the gene-RCD associations across the RCD landscape.

Every gene-RCD association in RCDome carries full source provenance and has been systematically checked against the published record, with both supporting and non-supporting outcomes retained. The compendium, extraction pipeline, and web platform are publicly available at https://github.com/BioCancerInformatics/RCDome.

## MATERIALS AND METHODS

The platform was built in three stages, compile (Channels A–D), validate (Phases 2A–2G, organized as Layers 0–4), and govern (Phase 3), followed by the web application. Each stage is described in turn.

### Study Design and Strategy Overview

RCDome was conceived as a comprehensive, gene-centric, multi-source knowledge platform encompassing the RCD-related entities represented in the governed source universe (Figure 1). The platform follows a layered, database-centered strategy in which each successive validation layer builds upon the database compilation foundation without overwriting earlier results: Layer 0 (Database Compilation), Layer 1 (National Center for Biotechnology Information (NCBI) + DeepSeek (deepseek.com) Abstract Validation), Layer 2 (Europe PMC + DeepSeek Full-Text Recovery), Layer 3 (Europe PMC XML Deep Retrieval-Augmented Generation (RAG)), and Layer 4 (Elicit Literature Mining; independent of the DeepSeek and Consensus channels, though overlapping with Elicit’s own Phase 3 discovery contribution). This layered architecture ensures that original database claims are preserved intact. The platform’s development followed a four-phase strategy: (i) establishment of a canonicalized RCD-related entity vocabulary through systematic literature and database discovery; (ii) multi-channel gene compilation from 20 association databases and resources organized into four compilation channels (A through D), with a fifth validation/literature-mining channel (E) added at validation; (iii) integration into a unified, traceability-compliant gene-RCD association matrix; and (iv) deployment as a publicly available interactive web platform.

**Figure 1.**
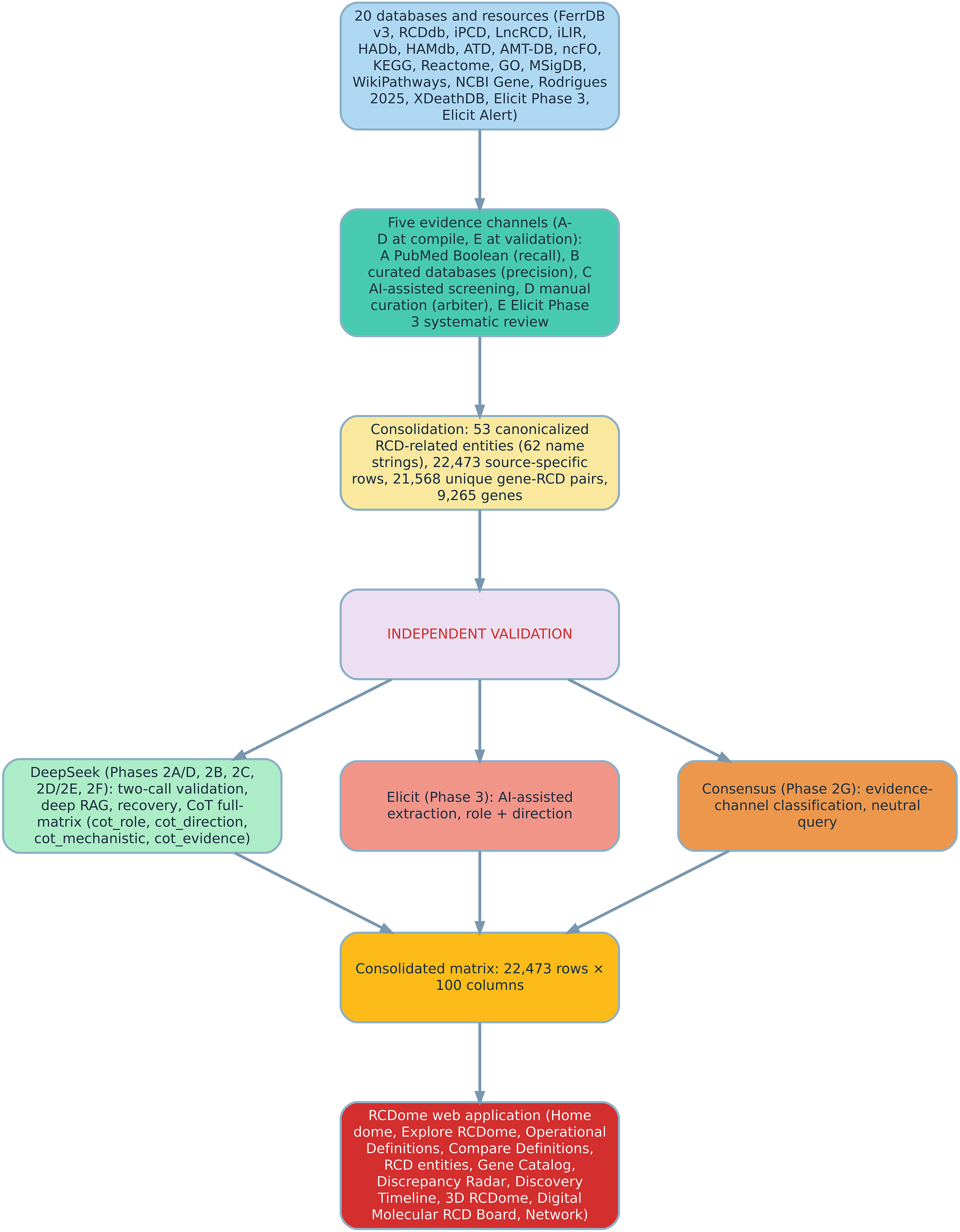
RCDome compilation and validation pipeline. RCDome is a human gene atlas of regulated cell death (RCD) assembled in a single top-to-bottom workflow. Twenty databases and resources constitute the input layer: ten RCD-specific curated resources (FerrDB version 3, RCDdb, iPCD, LncRCD, iLIR, HADb, HAMdb, ATD, AMT-DB and ncFO), five pathway and molecular databases (KEGG, Reactome, Gene Ontology, the Molecular Signatures Database and WikiPathways), the NCBI Gene resource, the multi-optosis pan-cancer gene set, XDeathDB, and two Elicit literature channels (the Phase 3 systematic review and the Alert review). These resources are compiled through five complementary channels: Channel A (programmatic PubMed Boolean searches) maximizes recall; Channel B (topic-specific database extraction) provides precision; Channel C (generative-AI-assisted literature screening) captures recent discoveries; Channel D (manual expert curation) serves as the final arbiter; and Channel E (Elicit Phase 3 systematic review) recovers gene-RCD associations from the full-text literature. Consolidation normalizes the 62 original database name strings into 53 canonicalized RCD-related entities, yielding 22,473 gene-RCD association rows across 9,265 unique human genes. The consolidated matrix then enters an independent validation stage in which three mutually independent analyses are applied in execution order. The first is the DeepSeek chain-of-thought classifier (Phases 2A through 2F), which performs two-call abstract validation, full-text recovery, deep retrieval-augmented generation and a full-matrix chain-of-thought classification that annotates every pair with four columns (cot_role, cot_direction, cot_mechanistic, cot_evidence) encoding chain-of-thought role, direction, mechanistic verdict and evidence. The second is the Elicit systematic-review extraction (Phase 3), which assigns functional role and direction of regulation. The third is the Consensus evidence-channel classification (Phase 2G), which queries each pair with a neutral formulation, the gene symbol followed only by “cell death”, so that evidence is retrieved without form-name priming. The three analyses are preserved in separate columns and are never conflated. The resulting consolidated matrix (22,473 rows and 100 columns) powers the RCDome web application, whose interface comprises the interactive dome, the Explore RCDome guide, Operational Definitions, Compare Definitions, the RCD entities browser, the Gene Catalog, the Discrepancy Radar, the Discovery Timeline, the 3D RCDome, the Digital Molecular RCD Board, and the Network module. Boxes are color-coded for visual distinction, and arrows indicate the flow of data through the pipeline.

### Establishment of the Canonicalized RCD-Related Entity Vocabulary

The initial RCD form vocabulary is based on our prior work establishing operational definitions for 25 RCD forms based on the NCCD 2018 consensus (Galluzzi et al., 2018) and extended through AI-curated definitions (Rodrigues de Souza et al., 2025, Rodrigues de Souza et al., 2026). These 25 forms comprised 20 executioner entities, 2 trigger events, and 3 functional categories (Supplementary Table S1).

Throughout RCDome, “canonicalized” refers exclusively to vocabulary normalization: heterogeneous source terms are mapped to a preferred RCDome identifier while their original terminology and provenance are retained. Canonicalization therefore does not imply that an entity is a canonical, mechanistically established, or human-validated RCD modality. Biological and ontological status is evaluated independently through the RCDome evidence and hard-gate framework.

During database discovery, newly emerging RCD forms were encountered, demanding systematic expansion. We executed a multi-layered literature and database surveillance strategy: (a) forward-citation tracking of NCCD 2018 article (Galluzzi et al., 2018) to identify post-consensus RCD forms; (b) PubMed systematic searches using the “-ptosis” and “-osis” suffix patterns combined with “cell death” terminology; (c) surveillance of bioRxiv and medRxiv preprints (2024–2026); and (d) cross-referencing against the RCD form inventories of all association databases and resources (Supplementary Table S2).

Candidate RCD forms were validated against five modality criteria (MC1–MC5) adapted from the NCCD framework (Galluzzi et al., 2018): (MC1) possession of distinct molecular machinery not fully subsumed by another form; (MC2) an identifiable biochemical or biophysical trigger; (MC3) a characteristic and reproducible morphological phenotype; (MC4) explicit naming in at least one peer-reviewed publication; and (MC5) pharmacological or genetic distinguishability from other forms. Co-activation umbrellas (e.g., PANoptosis (Pandian et al., 2022)) and morphological descriptors without distinct mechanisms were classified as conceptual categories. Following the biological curation, admission to the modality level (L1) was additionally governed by hard biological gates (BG0–BG6) (whole-cell death, causal machinery, functional necessity/distinguishability, supportive phenotype, naming/provenance, replication/maturity, and species scope), which operationalize MC1–MC5 with BG1 (machinery) and BG2 (necessity) obligatory. The resulting five-level ontology is encoded in the unified matrix as two columns (rcdome_level and rcdome_level_name; see the Matrix Descriptor tab of the Supplementary Tables (Excel)), preserving the external multi-optosis classification (rodr_category) untouched (Supplementary Table S3; Supplementary Table S4). Evidence grades in Supplementary Tables S3 and S6 use the A–D scale: A, well-established (>100 publications); B, established (10–100); C, emerging (2–10); D, proposed (1); N, not yet classified. The adjudication framework and its governance are documented in Supplementary Note S1; the complete evaluation architecture (RCD entity classification, gene-RCD association evidence, and pipeline quality gates) is summarized in Figure 2. Discordance between the independent classification layers is adjudicated, on the labels they share, by an independent vote and a human curator verdict that never replaces the original labels; this benchmark, the evidence-first adjudication of the final verdicts, and the repair of internal semantic contradictions are documented in Supplementary Note S2.

**Figure 2.**
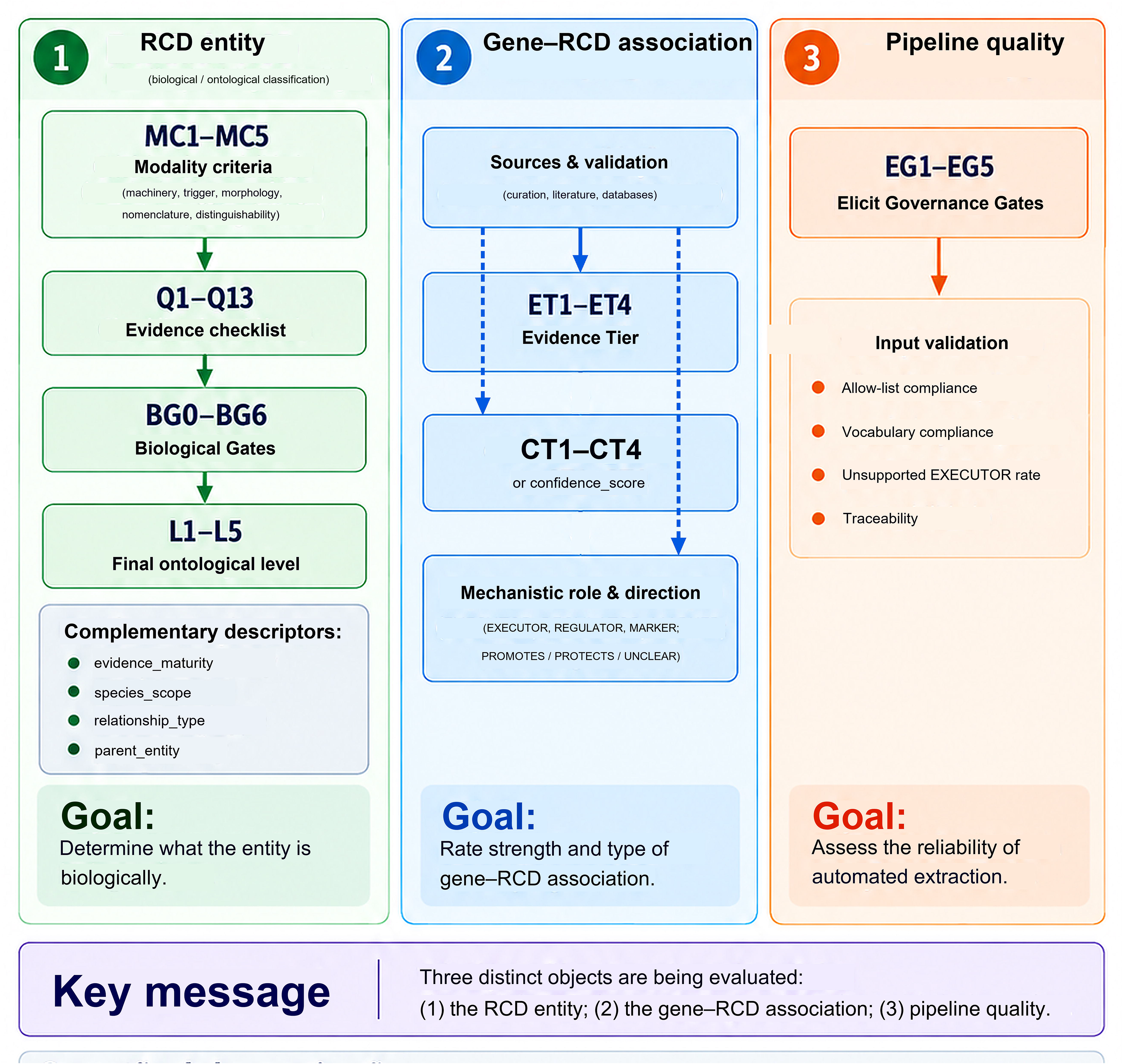
Evaluation architecture of RCDome. RCDome decomposes its curation into three objects that answer distinct scientific questions and are therefore assessed independently and never conflated. (1) For the RCD entity, the question is: “What kind of cell-death entity is this?” The answer is provided by modality criteria (MC1–MC5: machinery, trigger, morphology, nomenclature, distinguishability), an evidence checklist (Q1–Q13), biological gates (BG0–BG6, with BG1 machinery and BG2 necessity obligatory), and a final ontological level (L1–L5); complementary descriptors record evidence maturity, species scope, relationship type, and parent entity. (2) For the gene–RCD association, the question is: “Is gene X mechanistically linked to this entity?” The answer is provided by evidence tier (ET1–ET4) and a distinct confidence tier (CT1–CT4 or confidence_score), with mechanistic role and direction (EXECUTOR, REGULATOR, MARKER; PROMOTES/PROTECTS/UNCLEAR). (3) For pipeline quality, the question is: “Can the automated extraction be trusted?” The answer is provided by Elicit governance gates (EG1–EG5), input validation (allow-list and vocabulary compliance), unsupported EXECUTOR rate, and traceability. This three-way separation ensures that an ontological judgment is never mistaken for gene-level evidence, and that an extraction error is never mistaken for biological signal.

### Operational definitions of RCD entities

The operational definitions were developed through AI-assisted analysis under human supervision, using dedicated code and systematic human validation (assistant-draft/human-approval review and the Knock Test). The RCDome controlled vocabulary comprises 53 canonicalized RCD-related entities, where “canonicalized” denotes normalization to a preferred RCDome vocabulary identifier rather than recognition as a canonical or established RCD modality; it is derived from the unified gene-RCD evidence matrix (Dataset S1; 22,473 rows; 21,568 gene-RCD pairs). The controlled vocabulary is fixed and is not modified by downstream definition work, except manual curation. Each entity’s operational definition was assigned from the highest-priority source available, in the order: (i) the verbatim operational definitions of the NCCD (Galluzzi et al., 2018), treated as invariant (content and meaning never altered); (ii) the expanded operational definitions of our prior study ((Rodrigues de Souza et al., 2025), hereafter the multi-optosis dataset), preserved verbatim; and (iii) for entities lacking a committee definition, a de novo definition drafted from the original discovery article (founding PMID), written in the NCCD MC1–MC5 logic (distinct molecular machinery, biochemical/biophysical trigger, reproducible morphology, explicit naming in a peer-reviewed publication, and pharmacological or genetic modulation) and structured in the ten-field multi-optosis format, which the RCDome record expands to the 29 definition fields described below (plus a review-status annotation in the locked version).

### Human review granularity

The ontological classification of all 53 entities was curated individually by a primary curator (EM-A), who read, inspected, and approved each case separately (recorded per entity as ontology_curator_verified) during a single curation session rather than through unattended batch approval. For the 20 de novo RCDome-drafted operational definitions, the primary curator read and validated each definition against its founding evidence before marking it as approved; the one composite record (Senescence and Autophagy in Cancer) was validated by the provenance of its two parent entities rather than by a founding PMID of its own. The 20 NCCD committee definitions and the 12 multi-optosis (Rodrigues de Souza et al., 2025) definitions remained byte-identical (LOCKED-VERBATIM). Field-level evidence was established through the systematic assistant-draft/human-approval review and the Knock Test. Each ontological classification was subsequently assessed independently by two additional curators (VdSL and HACN), each reviewing all 53 entities on an entity-by-entity basis. Cases requiring reconsideration or involving disagreement among the independent assessments were jointly adjudicated by the curators, with the resulting consensus classification recorded as the final ontological assignment.

Each RCD entity is represented by a 42-field record comprising three evidence layers: (i) definition: the operational definition (prose), key molecules, inflammatory status, description of overlaps, category, and year of discovery; (ii) quantified ranking: the six ranking-score criteria (RS1–RS6) and the overall score S; and (iii) provenance: the founding PMID, reference PMID and citation, primary and secondary evidence PMIDs, the authoring source, the contributing source databases and channels, the provenance note, the definition status, the source-of-definition flag (NCCD-verbatim, multi-optosis-dataset, or RCDome-drafted), and seven orthogonal ontological dimensions (species_scope, evidence_maturity, relationship_type, parent_entity, ontology_curator_verified, independent_replication, primary_discovery_reference).

The *`evidence_maturity`* dimension records the consensus status of each entity as established, supported, emerging, legacy, or provisional. Quantified scoring. The six multi-optosis ranking-score criteria (RS1–RS6) were retained: RS1 extent of research (25%), RS2 direct involvement (25%), RS3 impact on cell/tissue homeostasis (20%), RS4 emerging potential (15%), RS5 biological significance (10%), and RS6 pathophysiological relevance (5%), each scored 1–10. Three criteria were reinterpreted from the cancer-oriented Rodrigues usage to the cell-biology scope of the RCDome: RS2 (“direct involvement in cancer therapy” → “direct involvement”), RS3 (“impact on cancer progression” → “impact on cell/tissue homeostasis”), and RS6 (“clinical relevance” → “pathophysiological relevance”). The overall score follows the deterministic weighted formula S = 0.25·RS1 + 0.25·RS2 + 0.20·RS3 + 0.15·RS4 + 0.10·RS5 + 0.05·RS6, which reproduces all 25 the multi-optosis anchors exactly. The RS1–RS6 values are expert judgments, not computable from gene counts (necrosis, 1,092 genes, is scored RS1 = 4 whereas cuproptosis, 77 genes, is scored RS1 = 7), demonstrating that the criteria are semantic. Consequently, for the entities without a committee definition the RS1–RS6 values were drafted from a rubric informed by matrix objective inputs (gene count, mechanistic classification ratio, evidence-tier distribution, and discovery year) and then subjected to assistant-draft/human-approval review.

Hard gates (L1 admission). Following the biological curation, admission of an entity to the modality level (L1) was governed by hard gates BG0-BG6: BG0 whole-cell death (irreversible), BG1 identifiable causal machinery (obligatory), BG2 functional necessity/distinguishability (obligatory), BG3 supportive phenotype/trigger, BG4 published name (necessary for traceability, not sufficient for mechanism), BG5 independent replication (a maturity input), and BG6 species scope. These gates operationalize the NCCD MC1-MC5 criteria (BG1 ∼ MC1 machinery; BG2 ∼ MC5 modulation; BG3 ∼ MC2/MC3 trigger and morphology; BG4 ∼ MC4 naming). Entities passing BG1 and BG2 were classified *`hard_gate_status`* = *`L1-ELIGIBLE`* (19/53); the remainder were *`NOT-L1`* (subtypes, triggers, umbrellas, morphology, contextual, and taxon-specific programs). The multi-optosis RS1-RS6 scoring remains an orthogonal ranking metric, not an admission gate. Independent replication (BG5) is now resolved for all 53 entities: none = 14 · limited = 1 · multiple = 38 (verdict standard: none = only founding paper · limited = 1–2 independent reproductions · multiple = ≥3 independent groups).

### Relationship to the NCCD “canonical” ontology

The 19 L1 modalities are deliberately a subset of a larger reference set, not a relabeling of it. The NCCD assigns operational definitions to 20 entities that it terms “canonical”; RCDome retains those 20 definitions verbatim (invariant) but applies a stricter, mechanistic admission criterion before any entity is elevated to L1. Of the 19 RCDome L1 entities, 9 carry NCCD-verbatim definitions (ferroptosis, necroptosis, pyroptosis, entotic cell death, NETotic cell death, parthanatos, MPT-driven necrosis, autophagy-dependent cell death, and lysosome-dependent cell death). Ten L1 entities fall outside the NCCD 2018 list: apoptosis (which the NCCD splits into extrinsic and intrinsic subtypes, both retained at L2 in RCDome) and nine modalities not included in the NCCD 2018 list, cuproptosis, disulfidptosis, methuosis, oxeiptosis, paraptosis, alkaliptosis, NECSO, mitoxyperilysis, and triaptosis, admitted on the strength of their distinct machinery and functional necessity. Conversely, 11 NCCD-verbatim entities are not L1 because they are subtypes (anoikis, extrinsic and intrinsic apoptosis, autosis), a synonym (lysosomal cell death, of lysosome-dependent cell death), a trigger (mitotic catastrophe), contextual or outcome programs (cellular senescence, efferocytosis, immunogenic cell death, mitotic death), or an umbrella term (programmed cell death). This asymmetry reflects a deliberate governance rule: an NCCD operational definition is necessary for traceability but not sufficient for modality status; L1 requires identifiable causal machinery (BG1) and functional necessity/distinguishability (BG2). RCDome therefore treats the NCCD list as a nomenclature baseline and re-screens it through the BG0–BG6 hard gates, rather than equating “named by the NCCD” with “mechanistically established as an independent death program.”

Four principles governed definition construction. (1) Invariance: published committee definitions (NCCD and the multi-optosis dataset) are preserved byte-identical to their sources and may only be complemented, never rewritten; verbatim integrity was verified programmatically against the PMC XML and the multi-optosis dataset spreadsheet. (2) Primacy of provenance (P0–P3): no entity is ever deleted; composite or derived labels are retained with an explicit provenance note and the evidence PMIDs of their parent entities; an entity outside the controlled vocabulary is marked retired rather than removed; and every entity carries its full provenance chain (source databases, source channels, reference PMID, provenance note). (3) No make-believe: no field is filled without a source; unsourced fields are left empty or marked “not specified”; PMIDs are verified to resolve in PubMed (Knock Test (verification that every cited PMID resolves to a real PubMed record)). (4) Human-in-the-loop verification: each drafted field was produced by an assistant (draft step) and is approved, edited, or rejected by a human (approval step), with every decision logged to its source for traceability.

Every operational definition is anchored to a reference PMID: the NCCD verbatim definitions to Galluzzi et al (Galluzzi et al., 2018) the multi-optosis definitions to their primary characterization PMIDs, and the de novo definitions to their founding PMID. The 25 multi-optosis entities additionally carry their full primary and secondary evidence PMIDs from the multi-optosis reference table. Each de novo definition was corroborated against independent evidence streams, coinage (a coining verb proximal to the entity name in the founding abstract), mechanism (the drafted key molecules appearing in the founding abstract), and PubMed consensus (a fresh search returning the founding PMID). Where a founding article was open-access, the full text (PMC) was used for claim-level validation; where no PMC full text was available, full-text recovery was additionally attempted via Europe PMC, and where the article remained paywalled the Consensus academic search Application Programming Interface (API) provided a full-text-derived takeaway as the richest available evidence, so that no definition was rejected on an abstract gap alone. Definitions whose founding article does not yet describe the MC2 trigger or MC3 morphology required by the NCCD logic were flagged criterion-pending, and umbrella categories were flagged umbrella-scope rather than as errors.

Consistent with the assistant-draft/human-approval principle of our prior multi-optosis study, in which the operational definitions of the five RCD forms not described by the NCCD were extracted by generative AI (*`GPT-4o`*, via the PDF Ai Drive tool, over a corpus of 6,603 free-access PDFs) and subsequently validated by manual review (Rodrigues de Souza et al., 2025), the RCDome employed a distinct AI instrument set. Definitions lacking a committee definition were drafted by an assistant and substantiated by a three-stream validation: (i) DeepSeek (*`deepseek-chat`* and *`deepseek-reasoner`*) for claim-level validation of each operational definition against the founding article’s full text where open-access and against its abstract plus a Consensus full-text-derived takeaway otherwise; (ii) the Consensus academic search API as an independent cross-reference for the founding reference; and (iii) PubMed E-utilities for deep search and PMID verification. Every AI-drafted field was then subjected to assistant-draft/human-approval review (approve, edit, or reject), with each decision logged to its source; no definition was accepted on the basis of AI output alone. The NCCD-verbatim and multi-optosis-verbatim definitions were preserved byte-identical to their sources and were not re-generated.

The RCDome decoder (the curated 53-entity reference of operational definitions, key molecules, ontological category, and scored evidence) was audited for completeness, score reproducibility (the composite score S (0.25·RS1 + 0.25·RS2 + 0.20·RS3 + 0.15·RS4 + 0.10·RS5 + 0.05·RS6) recomputed deterministically for every record), PMID verifiability (Knock Test), calibration (draft scores versus the locked anchors), and invariance (NCCD definitions byte-identical to the source). Agreement between the assistant-drafted and human-reviewed RS1–RS6 scores was assessed across 168 paired observations (28 entities × 6 criteria). All 168 score-pairs were identical between the preserved draft and final reviewed records (observed agreement = 100%), yielding Cohen’s κ = 1.00, exceeding the prespecified κ ≥ 0.80 target. All definition-building code and the resulting 53-entity decoder are provided in the reproducibility repository. Across the broader field-level review, all 336 drafted fields for the 28 newly-added entities were approved unchanged, with each decision retained in the audit record.

### Multi-Channel Gene Compilation Architecture

Gene-level evidence for each of the RCD forms was compiled through five independent but complementary channels (Channels A–D at compile, Channel E added at validation), designed to maximize both recall (Channel A: PubMed Boolean searches) and precision (Channels B, C, D: curated databases, AI-assisted screening, and manual curation; Channel E: Elicit Phase 3 systematic review) (Figure 1).

Channel A: Programmatic PubMed Boolean Searches. The curated gene database from (Rodrigues de Souza et al., 2025, Rodrigues de Souza et al., 2026) served as the gene universe for Boolean queries. For each gene in the Cancer Immunome and Beyond (CIB) database, we executed PubMed E-utilities queries pairing the gene symbol with each RCD form term and cancer-relevant filters: “[GENE]”[Gene] AND (“[*`RCD_TERM`*]”[Title/Abstract] OR “[*`RCD_ALIAS`*]”[Title/Abstract]) AND (cancer[Title/Abstract] OR tumor[Title/Abstract]). This channel produced the initial compendium of 5,913 RCD-associated genes validated in the multi-optosis dataset, which is exploded from slash-separated RCD form annotations into a one-gene-per-RCD-form-per-row format yielding 9,923 gene-RCD association pairs across all 25 forms (*`channel_a_rodrigues2025.py`*).

Channel B: Topic-Specific Database Extraction. Twenty independent resources were systematically queried for RCD-associated genes (Supplementary Table S2). Tier 1 comprised 10 RCD-specific curated databases: FerrDB v3 (22 RCD modalities, 3,481 unique genes; http://www.zhounan.org/ferrdb; (Zhou et al., 2026)), RCDdb (15 RCD types, 1,846 unique genes; http://chenyclab.com/RCDdb; (Wang et al., 2024)), iPCD (15 RCD forms, filtered to 3,393 Homo sapiens genes; https://ipcd.biocuckoo.cn; (Tang et al., 2022)), LncRCD (12 RCD types, 1,595 protein-coding target genes; https://lncrcddb.bio-database.com; (Zhao et al., 2026)), iLIR (6,087 LC3-interacting region proteins; https://ilir.warwick.ac.uk; (Jacomin et al., 2016)), HADb (371 human autophagy genes; https://autophagy.lu; (Homma et al., 2011)), HAMdb (551 human autophagy modulators; http://hamdb.scbdd.com; (Wang et al., 2018a)), ATD (87 autophagy-disease genes; http://auto2disease.nwsuaflmz.com; (Wang et al., 2018b)), AMT-DB (860 anti-tumor autophagy modulators; https://amtdb.vercel.app; (Fu et al., 2022)), and ncFO (90 experimentally verified ncRNA-ferroptosis associations; http://www.jianglab.cn/ncFO; (Zhou et al., 2023)).

Tier 2 comprised 5 pathway and molecular databases containing RCD-relevant subsets immersed within larger resources: KEGG (5 pathways via KGML parsing; https://rest.kegg.jp; (Kanehisa et al., 2017)), Reactome (Programmed Cell Death hierarchy R-HSA-5357801; https://reactome.org; (Gillespie et al., 2022)), Gene Ontology (7 RCD-specific GO terms via QuickGO REST API; https://www.ebi.ac.uk/QuickGO; (Gene Ontology et al., 2023)), MSigDB/GSEA (12 RCD-relevant gene sets in GMT format; https://www.gsea-msigdb.org; (Liberzon et al., 2011)), and WikiPathways (https://www.wikipathways.org (Martens et al., 2021)). STRING (https://string-db.org; (Szklarczyk et al., 2023)) and BioGRID (https://thebiogrid.org; (Oughtred et al., 2021)) were used for protein-protein interaction annotation rather than gene extraction.

Tier 3 comprised NCBI Gene (32 RCD form queries (run against the forms known at that pipeline stage; the final 53-form compilation retains 29 NCBI_Gene-contributed forms after synonym consolidation) via E-utilities API; https://eutils.ncbi.nlm.nih.gov; (Brown et al., 2015)). Tier 4 omic-annotation resources (TCGA, https://www.cancer.gov/tcga; (Cancer Genome Atlas Research et al., 2013); GEO, https://www.ncbi.nlm.nih.gov/geo; (Clough et al., 2024); CPTAC, https://proteomics.cancer.gov/programs/cptac; (Ellis et al., 2013); CCLE, https://sites.broadinstitute.org/ccle; (Barretina et al., 2012); GDSC, https://www.depmap.org/portal/gdsc; (Garnett et al., 2012); and DepMap, https://depmap.org; (Tsherniak et al., 2017)) were used for omic layer annotation rather than gene extraction.

MSigDB was extracted through a systematic, curated process rather than a fixed gene-set list. A manual, biologically oriented curation of all 53 RCD entities in MSigDB was performed (release 2026.1), searching each entity by its normalized name plus complementary terms (e.g., APOPTOTIC, NECROPTOTIC, PYROPTOTIC) to recover gene sets that a name-only search would miss. Each of the 360 candidate gene sets was individually classified as include (251: 238 new plus 13 already present in the matrix) or exclude (109) with a documented biological rationale (RCDome_MSigDB_Manual_Curation.xlsx): inclusion prioritized gene sets representing cell-death processes, executioner machinery, or form-specific signaling/regulation, while exclusion rejected drug, cancer-type, pathogen, or expression signatures that reflect context rather than RCD machinery. Of the 238 new inclusion sets, 236 matched the MSigDB 2026.1 release and were integrated; one set (REACTOME_EXTRINSIC_PATHWAY_FOR_APOPTOSIS) was archived as an obsolete Reactome build and excluded, while another (GOBP_ACTIVATION_OF_CYSTEINE_TYPE_ENDOPEPTIDASE_ACTIVITY_INVOLVED_IN_APOPT OTIC_PROCESS) had been renamed and was superseded by its successor (GOMF_CYSTEINE_TYPE_ENDOPEPTIDASE_ACTIVATOR_ACTIVITY_INVOLVED_IN_APOPTOTIC_PROCESS). This curation contributed 2,501 genes across 4,550 gene-RCD pairs: 1,832 pairs were created as MSigDB-only rows (spanning 1,701 genes), and 2,718 pairs corroborated already-present pairs (MSigDB appended as an additional source). The extraction code (channel_b_msigdb.py) was amended to read the curated list directly, making the selection reproducible and auditable.

ncFO (Zhou et al., 2023), a curated database of experimentally verified ncRNA-ferroptosis associations, was integrated as a corroborating source for Ferroptosis: 50 HGNC-validated human miRNA and lncRNA genes were appended to existing Ferroptosis pairs (one new pair, MIR92A1); three entries were excluded (two clone names without HGNC identifiers and one miRNA withdrawn by miRBase). ncFO’s human circRNA entries were documented but not integrated, because circRNAs lack HGNC nomenclature and a complete circRNA layer would require class-wide curation across all RCD forms.

We also considered the RCDscore compendium (Zhang et al., 2024), a pan-cancer resource that collects 18 RCD signatures (11,206 gene-RCD pairs, 7,460 genes) from prior studies to predict overall survival and immune checkpoint inhibitor response. RCDscore was not integrated into the RCDome matrix because it catalogues gene pathways rather than curated gene-RCD association pairs: its signatures are broad Gene Set Variation Analysis (GSVA)/ssGSEA gene sets (up to 3,449 genes for a single form) whose membership reflects pathway involvement, not evidence of a specific role in the death mechanism. Accordingly, the 67.8% of gene-RCD form pairs that do not overlap with the matrix were excluded as pathway-level genes lacking curated RCD-specific support; the remaining 32.2% overlap consists of genes already present in the matrix and would add no new associations. RCDscore is cited here as an independent clinical-prognosis resource validating the translational relevance of RCD gene signatures.

For each database, extraction was performed using dedicated Python scripts (*`channel_b_*.py`*), with all gene symbols validated against HGNC nomenclature. Every gene-RCD association was annotated with its source database, evidence type, and extraction timestamp.

Channel C: AI-Assisted Literature Exploration. The Consensus academic search engine (https://consensus.app) was employed to screen the literature for gene-RCD associations not captured by Channels A or B. Every AI-extracted association was flagged as *`curator_verified`* = FALSE until manual expert review; following the completed review, all 53 entities carry *`ontology_curator_verified`* = TRUE (each entity individually reviewed and approved, logged 2026-09-05). The Consensus.ai academic search engine was used for semantic searches across 200M+ papers, returning claim-level extractions with citation context.

Channel D: Manual Expert Curation: Primary discovery papers for each RCD form were read in full to extract reference gene sets. NCCD consensus documents, major mechanistic reviews, and cancer-context reviews were mined for molecular machinery annotations. Conflicts between Channels A, B, and C were adjudicated by expert review of the primary literature.

### Gene-RCD Association Traceability Specification

All gene-RCD associations entering the RCDome knowledge base must satisfy a strict traceability contract: (i) one gene per RCD form per row: multi-form associations are exploded into individual rows; (ii) every source database is explicitly named using a closed vocabulary of 20 controlled database identifiers; (iii) every RCD form is named using the closed vocabulary of 53 RCD-related entities; (iv) forbidden patterns including “and others”, “etc.”, “multiple”, and “various” are automatically rejected at integration; and (v) every association was individually human-curated, as recorded by the association-level curation flag (*`curator_verified`*, set during Channel C extraction), distinct from the entity-level *`ontology_curator_verified`* flag in the decoder, which records that each of the 53 entities, not each association, was individually reviewed and approved (logged 2026-09-05). An automated rejection engine enforces these rules at the integration step.

### Data Integration and Unified gene-RCD Matrix

Outputs from the association-producing channels (A, B, and E) were merged into a unified gene-RCD association matrix using (gene_symbol, rcd_form) as the primary merge key, preserving the original RCD form name exactly as recorded in each source database. Channels C (AI-assisted literature screening) and D (manual expert curation and conflict adjudication) acted as governance steps on these records rather than as additional association sources. A parallel rcd_form_canonical column was reserved for subsequent expert-governed unification of synonymous form names (e.g., “Entosis” to “Entotic cell death”).

Each gene-RCD association was annotated with an *`evidence_quality`* classification reflecting the nature of the source evidence. Associations derived from manually curated databases (FerrDB, RCDdb, iPCD, LncRCD, iLIR, HADb, HAMdb, ATD, AMT-DB, KEGG, Reactome, GO, MSigDB, ncFO) and the multi-optosis dataset were classified as curated. Associations from the NCBI Gene E-utilities API, which uses keyword-based record matching rather than explicit curation, were classified as keyword. Rows supported exclusively by keyword evidence (n = 10,857) were excluded following a spot audit that revealed 0% overlap with known RCD executioner and regulator genes. Fifteen keyword-only rows for NECSO, a newly described entity whose gene associations were available solely through the NCBI Gene keyword search at compilation time, were retained and are flagged as such in the evidence_quality column. Rows with curated-only or mixed curated+keyword evidence were retained (n = 18,991). Subsequent database integrations through the external audit (Reactome pathway-specific gene sets, WikiPathways, and XDeathDB, all classified as curated) and the Elicit Phase 3 systematic review (classified as *`ai_extracted`*) expanded the matrix to the final 22,473 rows.

Additionally, all gene symbols in the merged matrix were validated against the HGNC complete approved symbol set (n = 45,030; https://rest.genenames.org; (Seal et al., 2023)). Symbols failing HGNC validation (n = 638 unique, 696 rows) were excluded. These comprised UniProt accessions misassigned as gene symbols (from iLIR mapping artifacts), protein family abbreviations (AMPK, PI3K, JNK), non-human orthologs (Gm21596, OsFER2), circRNA names, and chemical/protein complex names erroneously extracted as gene identifiers. Their exclusion is registered in the *`gene_symbol_status`* column of the raw matrix and documented in the extraction audit log.

After both exclusion steps (NCBI Gene keyword-only noise and HGNC-invalid symbols), the final unified gene-RCD matrix comprised 22,473 rows representing 9,265 unique HGNC-validated gene symbols across 62 original RCD form names (Dataset S1).

### RCD Form Vocabulary Expansion, External Audit, and Governance

During manuscript preparation, nine additional cell death-related terms were proposed for inclusion (NECSO, Triaptosis, Mitoxyperilysis, Eryptosis, Cobaltosis, Calcioptosis, Corneoptosis, plus Pyronecrosis and Spectosis already catalogued). Each was evaluated against the same five-criteria framework. Four met the threshold for inclusion in the controlled vocabulary (NECSO, Triaptosis, Mitoxyperilysis, Eryptosis). Three further forms encountered during database discovery (Floatptosis, Lysozincrosis, and Metalloptosis) were also admitted, Floatptosis and Lysozincrosis as a cell-type program (L4) and a subtype (L2), respectively, and Metalloptosis as an umbrella term, completing the 53-entity vocabulary. An external audit subsequently integrated three additional databases (Reactome pathway-specific gene sets, WikiPathways, and XDeathDB), which contributed association records to existing entities rather than new entities (Supplementary Table S3).

Three proposals were formally excluded on fundamental grounds:

1. Cobaltosis: This term describes systemic cobalt toxicity (cobaltism) from metal-on-metal hip implants, not a genetically encoded cell death program. The cellular damage is a consequence of heavy metal poisoning, not a regulated entity with distinct molecular machinery. No unique trigger, executioner, or inhibitor distinguishes it from general metal toxicity.
2. Calcioptosis: Calcium is a universal second messenger involved in the initiation of multiple RCD forms, including apoptosis (cytochrome c release), MPT-driven necrosis (mitochondrial permeability transition), excitotoxicity, and eryptosis. Naming a distinct death form after calcium overload conflates a shared trigger with a unique entity. No molecular machinery separates proposed calcioptosis from existing Ca2+-mediated death mechanisms.
3. Corneoptosis: The supporting literature describes cell death in corneal tissue across a broad temporal range (1977-2026), consistent with apoptosis and necrosis occurring in a specific anatomical context rather than a distinct RCD form. Insufficient evidence exists for unique molecular machinery, trigger, or pharmacological distinguishability. Two further terms were surfaced by post-freeze literature surveillance, which included a news feature on the expanding cell death landscape (Dance 2026). Both terms were evaluated against the same framework. Independently of their biological status, neither term is present in any of the 20 association databases and resources used in the compilation (ruptosis entered specialized databases only from October 2026, after the matrix data freeze), so under the no-invented-names rule (every RCDome entity originates from at least one source database’s published terminology), neither could be admitted as a compendium entity. They are nevertheless ruled out explicitly, on both database-absence and biological-gate grounds, rather than left unaddressed:
4. Ruptosis: This term describes the explosive disintegration of specialized effector cells (“ruptoblasts”) in planarians, which release antimicrobial material onto invading microbes as part of a hormone-surveillance immune-defense program. It is an invertebrate-specific effector program documented in a single discovery article (Chai et al., 2026) at the time of writing; no human orthologous machinery has been identified, no human gene-level evidence exists in the compendium, and it has not been independently replicated or shown to be pharmacologically or genetically distinguishable from lytic effector degranulation in human cells. It therefore fails the species-scope (BG6) and independent-replication (BG5) gates and, as with corneoptosis, names a context-specific process rather than a distinct human RCD modality.
5. Anastasis: This term denotes the reversal of an initiated death program and the subsequent survival of the cell, documented for apoptosis (Tang et al., 2012, Tang et al., 2018). and subsequently proposed for ferroptosis and necroptosis. It is not a cell death form but the failure of one to complete: a rescue and survival process that fails the most fundamental admission gate (BG0, whole-cell death, irreversible) by definition. It is excluded on the same categorical basis as the non-lethal processes that the NCCD itself distinguishes from cell death modalities.

The exclusion of cobaltosis, calcioptosis, corneoptosis, ruptosis, and anastasis is documented here to establish a precedent: not every “-ptosis”, “-osis”, or “-stasis” term proposed in the literature qualifies as a distinct regulated cell death form; the five-criteria framework and the BG0–BG6 hard gates serve as the gatekeeper, and the no-invented-names rule independently bars terms absent from every source database. Forms named after the matrix data freeze, absent from the source databases, or lacking human gene-level evidence are acknowledged and ruled out rather than silently omitted.

The unified matrix contained 62 distinct original RCD form name strings due to heterogeneous naming conventions across source databases. To enable cross-database comparison, a normalization mapping was applied to consolidate synonymous names into 53 RCD-related entities, all with gene-level evidence in the unified matrix. The original name (*`rcd_form`*) is preserved in every row; the normalized name (*`rcd_form_canonical`*) is added as a parallel column. All normalization decisions are registered in the extraction audit log and are fully reversible. Two original name strings are mapped source-dependently to preserve source-specific classification: ‘Lysosomal cell death’ is resolved to ‘Lysosomal cell death’ (subtype) for XDeathDB rows and to ‘Lysosome-dependent cell death’ (modality) for iPCD rows, while ‘Mitotic death’ is resolved to ‘Mitotic catastrophe’ (trigger) for FerrDB rows and to ‘Mitotic death’ (subtype) for XDeathDB and Elicit rows. The two entities are not synonyms: mitotic catastrophe is the oncosuppressive process that is not an RCD form per se, and mitotic death is the RCD outcome driven by it. The 11 synonym renames are listed in Supplementary Table S5.

### Normalization mapping (11 renames to produce 53 RCD-related entities, all with gene-level evidence)

The final RCD form vocabulary consolidates 62 original name strings from 20 association databases and resources into 53 RCD-related entities through 9 synonym-merge groups (11 name eliminations, 2 split names). All 53 RCD-related entities carry gene-level evidence in the unified matrix (Supplementary Table S3). The full expansion and normalization methodology is documented in the Materials and Methods (Normalization mapping) and Supplementary Table S3. The newly admitted RCD-related entities are listed in Supplementary Table S6.

All 53 normalized names were retained as-is (kept verbatim, without renaming or merging, because at least one source database had already contributed them in their normalized form). No names were invented; every normalized term originates from at least one database’s published terminology.

### Phase 2: Automated gene-RCD Association Validation

#### Phase 2A: Two-Call Independent Validation

To systematically evaluate every gene-RCD association in the compilation matrix, we implemented a two-call independent LLM-based verification pipeline. The validation operates on all 21,568 unique (gene, RCD-related entity) pairs in the unified matrix, adding evidence tiers, literature support, and verification status to each row. Throughout this work, matrix rows refer to the 22,473 source-specific association records, whereas unique canonical gene-RCD pairs refer to the 21,568 distinct gene and RCD entity combinations after deduplication.

Two distinct validations are performed and never conflated: (i) gene-RCD association validation, whether a gene and an RCD entity are genuinely linked in the published record, which is automated and scalable across the three independent channels; and (ii) RCD modality status validation: whether an entity constitutes a bona fide human cell-death modality, which is an ontological judgment performed by human curation, not by the automated channels.

### Two-Call Independent Validation Architecture

The pipeline executes two completely independent LLM calls per gene-RCD pair, with no cross-contamination between them:

#### Call A (Internal Knowledge)

The LLM (DeepSeek-chat) is asked whether gene X is mechanistically involved in RCD form Y, relying solely on its training data (knowledge cutoff: late 2024). No external data is provided. The LLM responds CONFIRMED or UNCLEAR.

#### Call B (PubMed Abstract Evidence)

PubMed E-utilities API retrieves ALL available PMIDs via paginated E-utilities queries, with no arbitrary cap on paper count. Evidence force is derived from the total paper count returned by PubMed E-search (instant, no abstract fetch required). For content verification, the first 100 most relevant abstracts (5 batches of 20) are presented to the LLM. The full paper count serves as the quantitative evidence force metric, while the LLM verification of the top papers confirms the qualitative nature of the association. This design balances comprehensive evidence quantification with computational feasibility, and presents them to the LLM with the same question. The LLM responds CONFIRMED or UNCLEAR based solely on the abstract evidence.

#### DeepSeek Validation

PMIDs recovered by Europe PMC are used to fetch PubMed abstracts via E-utilities EFetch, which are then validated through the same two-call independent DeepSeek architecture used in Phase 2A. Call A uses internal LLM knowledge; Call B evaluates the abstract evidence from the newly discovered PMIDs. Pairs confirmed by both calls are upgraded to MECHANISTIC.

The two calls are merged independently according to a predefined matrix: both CONFIRMED → CONFIRMED; Call A CONFIRMED + Call B UNCLEAR → *NEEDS_LITERATURE* (LLM knowledge exists but literature is lacking); Call A UNCLEAR + Call B CONFIRMED → *NEEDS_CONTEXT* (literature support exists but the LLM is uncertain); both UNCLEAR → UNCLEAR.

### Evidence Tier Assignment

Each matrix row (gene-RCD association record) received an evidence tier (ET1-ET4) based on its own source combination and LLM validation status; a pair contributed by multiple sources could therefore carry different tiers across its rows:

- ET1 (Gold Standard): Four or more independent database sources, or two or more sources with RCDdb-provided PMIDs.
- ET2 (Strong Evidence): Three or more sources, or confirmed by both LLM calls.
- ET3 (Moderate Evidence): Two sources with LLM confirmation, or one source with LLM confirmation.
- ET4 (Weak Evidence): Single source without LLM confirmation, or LLM unable to verify.

Separately, a confidence score was computed as a weighted sum of the compilation channels (Channel D = 4, Channel B = 3, Channel A = 1, Channel C = 1) and recorded as an independent confidence tier (CT1 = 7–9, CT2 = 4–6, CT3 = 2–3, CT4 = 0–1). Channels C and D are validation/annotation layers and are not retained in the final source_channels field (which records A, B, and E); Elicit (Channel E) rows carry a fixed score of 3. This confidence tier is a metric distinct from the evidence tier (ET1–ET4) defined above and must not be conflated with it. This confidence score applied only to rows carrying compilation-channel evidence (Channels A through D). The 1,874 rows contributed solely by a single database or by the Elicit validation layer lay outside this scoring universe and therefore carried no confidence tier. No row reached a score of 7 to 9, so the CT1 tier remained unpopulated.

### Anti-Block and Rate-Limiting Strategy

To ensure compliance with API rate limits without interruption, global thread-safe rate limiters were implemented: LLM calls are capped at 30 requests per minute (2-second inter-call gap, well below the provider’s limit), and PubMed E-utilities calls at 8 requests per second (below the 10/second limit with API key). Validation is parallelized across 2 concurrent workers, with progress saved every 2,000 pairs to enable resumption if interrupted. The unified matrix is modified only upon successful completion of all validation; intermediate progress is stored in a separate JSON checkpoint file (*llm_verification_progress.json*), ensuring that the matrix remains uncorrupted if the pipeline is interrupted. The validation pipeline is implemented as *`channel_c_evidence_tier.py`* in the extraction code repository. All LLM API keys are stored in environment variables (.env file), never hardcoded.

### Independent Accuracy Assessment (Phase 2A/D, comprising the Phase 2A two-call validation and the MSigDB extension; distinct from Phase 2D)

The compilation amassed 319,560 unique PMIDs across its four evidence channels (curated primary, LLM-confirmed, Europe PMC, and Elicit). In a dedicated quality-assurance step, the 10,141 curated PMIDs that constitute the association evidence were resolved to their PubMed article titles through the E-utilities API.

An independent accuracy test was embedded in the pipeline. A ground truth set of 17 well-established gene-RCD associations (spanning 10 RCD forms, including 4 negative controls where the gene is known NOT to be involved) is processed through the same two-call validation. Accuracy is reported as the percentage of correct classifications. This assessment is documented in the extraction audit log and is reproducible via the --validate flag. The Phase 2A/D validation output columns are described in Supplementary Table S7.

#### Phase 2A/D MSigDB extension

The 1,832 MSigDB-only gene-set pairs were validated with the same two-call architecture extended to capture functional role and direction: the internal-knowledge call and the abstract-evidence call each return a structured verdict, mechanistic (yes/no), role (executor/regulator/marker/uncertain), and direction (promotes/protects), and the merged verdict assigns the mechanistic classification together with the role and direction. Because ‘Programmed cell death’ is a parent form whose child terms (apoptosis, necroptosis, pyroptosis, ferroptosis) dominate the literature, the PubMed query for those 1,512 pairs was broadened to the child terms rather than the literal parent name.

### Phase 2B: Europe PMC Full-Text XML Deep RAG Evidence Extraction

Europe PMC Full-Text Search. Europe PMC indexes both abstracts and full-text articles; Phase 2B queried the Europe PMC full-text index and processed 167,868 open-access full-text articles available for XML-based analysis, enabling retrieval of gene-RCD co-occurrences hidden from PubMed’s (gene[Gene]) AND (rcd[Title/Abstract]) strategy. For each non-MECHANISTIC pair, the query ({gene}) AND ({rcd}) is executed against the full-text index, returning PMIDs, PMC IDs, and total hit counts.

To extract deep, verbatim evidence from the primary literature, we developed a three-pass Retrieval-Augmented Generation (RAG) pipeline that processes open-access full-text XML articles (via Europe PMC API) as the source of truth. This complements Phase 2A (abstract-level validation) by capturing evidence buried in paper bodies that never appears in titles or abstracts.

### Combined-Pass Per-Article Architecture

Each article was processed through a single combined-pass LLM call per text chunk, which performs three tasks in one request:

Task 1: Gene Symbol Extraction. The full text is split into overlapping chunks of 2,000 characters. The LLM is prompted to extract all human gene symbols mentioned in the passage. Returned gene symbols are cross-referenced against the HGNC-approved 45,030-symbol set. Only validated human genes proceed to Task 2.

Task 2: RCD Form Detection. For chunks containing matched gene symbols, the LLM is asked which forms of regulated cell death are discussed and which genes are linked to each form. The prompt includes the complete list of 53 RCD-related entities to guide detection. Only chunks with confirmed gene-RCD associations proceed to Task 3.

Task 3: Verbatim Evidence Extraction. For confirmed gene-RCD pairs, the LLM extracts exact verbatim sentences from the text, classifies the evidence type (DIRECT_EXPERIMENTAL, GENETIC_INTERVENTION, PHARMACOLOGICAL, EXPRESSION_CORRELATION, or CITATION_ONLY), determines the gene’s mechanistic role (EXECUTOR, REGULATOR, MARKER, SUBSTRATE, or INTERACTOR), and assigns a direction (PROMOTES, PROTECTS, or *`CONTEXT_DEPENDENT`*). Role and direction are evidence-based labels assigned from the cited literature and the model; they are not determinations made by our own experiments. A gene labeled REGULATOR is recorded as such in the sources, not proven to regulate the pathway by a specific assay.

### Hallucination Prevention and Governance

A strict anti-hallucination framework operates at the code level. Every extracted sentence is verified against the complete raw XML text using exact substring matching (verify_sentence()). Sentences not found in the source article are rejected with a logged *`HALLUCINATION_DETECTED`* flag and excluded from accumulation. Per-PDF hallucination rates are monitored; rates exceeding 30% trigger a CRITICAL alert. A global hallucination counter and per-incident log provide full auditability.

### Multi-Paper Evidence Aggregation

Evidence for each gene-RCD pair is aggregated across all processed articles. Governance rules determine the final classification: a gene’s role requires consensus from at least two papers or one paper with DIRECT_EXPERIMENTAL evidence. Direction conflicts (PROMOTES vs PROTECTS) are resolved to *`CONTEXT_DEPENDENT`* and flagged for manual review. Confidence is calculated as HIGH (5 or more papers with consensus role), MEDIUM (2 or more papers or at least one DIRECT_EXPERIMENTAL), or LOW (single paper without direct experimental support). Evidence score (fraction of papers with DIRECT_EXPERIMENTAL evidence) provides a continuous quality metric.

### Ground Truth Audit (Phase 2B)

Before processing the full corpus, the pipeline executes a ground truth audit (--validate mode) on four landmark papers with definitively known gene-RCD associations: GPX4/Ferroptosis (Yang et al., 2014), MLKL/Necroptosis (Cai et al., 2014), GSDMD/Pyroptosis (Sborgi et al., 2016), and (Tsvetkov et al., 2022) (FDX1/Cuproptosis). The audit measures gene symbol recall, RCD form detection accuracy, role classification accuracy, sentence verifiability, and hallucination rate. It yielded 100% gene recall, 100% RCD form detection, and 100% role classification across the four papers, with 53 of the 54 extracted evidence sentences verified verbatim and the single non-verbatim sentence rejected by the anti-hallucination filter.

### Output and Matrix Integration

Phase 2B full-text XML deep evidence extraction (*`channel_c_deep_xml_rag.py`*) processes all 21,283 gene-RCD pairs carrying at least one PMID. PMID-to-PMCID conversion yields open-access PMC full-text articles. An 8-thread parallel architecture fetches XML and applies a combined-pass DeepSeek LLM retrieval-augmented generation pipeline that extracts gene symbols, detects RCD entities against the 53-entity controlled vocabulary, and captures verbatim evidence sentences in a single API call per text chunk. An anti-hallucination framework performs exact-string verification of every extracted sentence against the source XML; sentences not found verbatim are rejected and logged. Evidence is deduplicated by PMCID and aggregated across papers with multi-paper consensus scoring. Ten new columns are added to the matrix (*`deep_pmids_total`*, *`deep_pmids_free`*, *`deep_pmids_skipped`*, *`deep_paper_count`*, *`deep_best_sentence`*, *`deep_best_evidence`*, *`deep_best_role`*, *`deep_best_direction`*, *`deep_confidence`*, *`deep_evidence_score`*). Three existing columns are upgraded where deep evidence confirms Phase 2A/D classifications: *`evidence_tier`* (ET2 to ET1 when *`deep_confidence`*=HIGH and role=EXECUTOR), *`rcd_gene_type`* (populated from *`deep_best_role`* for empty rows), and *`mechanistic_class`* (LITERATURE_GAP to MECHANISTIC when *`deep_confidence`* is HIGH or MEDIUM). Processing is resumable via checkpoint files (*phase2b_checkpoint.json*, *phase2b_pair_results.json*) and generates a governance report documenting every PMCID processed, every hallucination rejected, and the full evidence distribution.

### Phase 2C: Full-Text Europe PMC Recovery

#### Output and Traceability

Phase 2C adds four new columns to the unified matrix without overwriting Phase 2A values. Phase 2C is distinct from Phase 2B (the deep RAG step): 2C recovers PMIDs, whereas 2B extracts deep evidence sentences. These columns capture MECHANISTIC classifications from full-text Europe PMC recovery, in addition to the 3,063 MECHANISTIC rows from Phase 2A/D abstract validation: *`epmc_hits`* (full-text hit count), *`epmc_status`* (Phase 2C LLM verdict), *`epmc_pmids`* (PMIDs recovered by Europe PMC), and phase2c_mechanistic_class (combined classification). Phase 2A columns (*`mechanistic_class`*, *`llm_status`*, *`llm_pmids`*) remain untouched, preserving full traceability of the validation provenance. The pipeline is implemented as *`channel_c_recovery_europe_pmc.py`*, runs autonomously with auto-restart on failure, and saves progress every 10 pairs.

### Phase 2D and 2E: Extended Literature Validation After External Audit

An external audit identified coverage gaps in the initial extraction: (a) Reactome contained pathway-specific gene sets (REACTOME_AUTOPHAGY, REACTOME_APOPTOSIS, REACTOME_REGULATED_NECROSIS) not captured by the parent ‘Programmed Cell Death’ extraction; (b) WikiPathways gene sets were cited but not separately extracted; and (c) XDeathDB (Gadepalli et al., 2021) was identified as an additional source by the external audit but had not yet been integrated at that stage. These gaps were closed by extracting 378 Reactome genes (3 pathways), 286 WikiPathways genes (4 pathways), and 5,487 XDeathDB genes (12 RCD forms) through manual CSV download, retaining 339 Reactome, 253 WikiPathways, and 895 XDeathDB unique genes in the matrix from the XDeathDB Shiny application https://pcm2019.shinyapps.io/XDeathDB/). The audit also identified that the *`gene_type`* column conflated cancer gene roles (driver/suppressor from multi-optosis) with RCD functional roles (EXECUTOR/REGULATOR compiled from Elicit Phase 3 and other sources), leading to a column split into *`cancer_gene_type`* and *`rcd_gene_type`*. Phase 2D (NCBI+DeepSeek) validated the resulting 1,341 novel unique pairs, and Phase 2E (Europe PMC+DeepSeek) performed full-text recovery on the remaining 1,724 non-MECHANISTIC pairs, recovering 588 MECHANISTIC associations through Europe PMC full-text validation.

### Phase 2F: Chain-of-Thought Full-Matrix Classification

The full run classified all 21,566 unique (gene, RCD-related entity) pairs using the deepseek-v4-pro model with thinking enabled and reasoning effort set to maximum. The two late-added MSigDB pairs (NLRX1 and PPIF, MPT-driven necrosis) were classified separately: their chain-of-thought columns by a five-vote mini-run, and their seven-class mechanistic classification by the Phase 2A/D two-call, yielding the final 21,568 unique pairs.

Four new columns were added to the unified matrix without overwriting any Phase 2A–2E values: *`cot_role`*, *`cot_direction`*, *`cot_mechanistic`*, and *`cot_evidence`* (the concatenated verdict). Processing was resumable via a checkpoint file saved every ten pairs, with an off-peak supervisor for autonomous restart.

Post-hoc reconciliation repaired 84 self-inconsistent rows (mechanistic NO with a confident role or direction), and 231 rows with genuine role/direction uncertainty, 178 regulators with unclear direction and 53 mechanistically-involved pairs with unresolved role, were retained as unresolved rather than forced into a confident category. Two gene-RCD pairs added to the matrix after the full run (NLRX1 and PPIF with MPT-driven necrosis) were classified separately with the same logic using a five-vote majority, completing 100% pair coverage (21,568/21,568).

#### Model-selection benchmark

To select the functional-role classifier, three DeepSeek model configurations were benchmarked on 26 well-established gene-RCD pairs (10 executors, 8 regulators, and 8 negative controls) (Supplementary Note S3; Figure 3A): the baseline *`deepseek-chat`* (81% combined accuracy, 38% false positives) and chain-of-thought *`deepseek-reasoner`* (73%, 25%). A governance-augmented tool-call loop with a hand-curated reference covering the 18 unique positive genes achieved 100% combined accuracy and zero false positives on all 26 pairs (the 18 positives plus 8 form-mismatch negatives), but that reference covers only a small fraction of the compendium (about 5% of pairs have a complete role, direction, and mechanistic classification to reference). A self-provided version of the same loop, which instead makes the model state its own molecular mechanism, known RCD forms, and the specific form its role belongs to, then self-checks those statements, also achieved 100% combined accuracy and zero false positives without any external reference. Because it is fully independent and scalable, the self-provided loop was adopted as the role-classification engine for the full 21,568-pair matrix. Structured-output mode (guaranteed JSON) was also adopted across the extraction pipeline to eliminate parsing failures.

**Figure 3.**
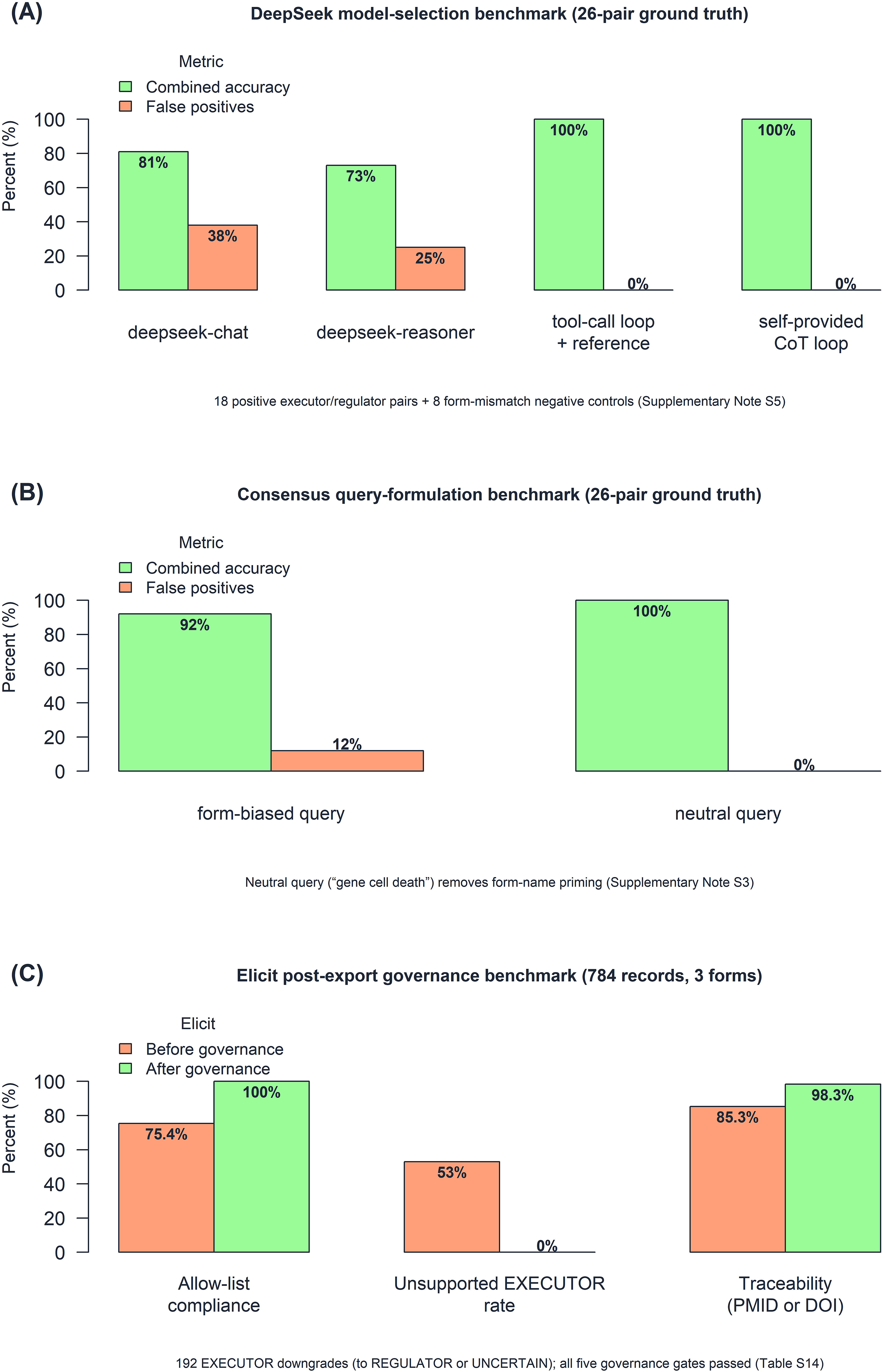
Benchmark evaluation of the three independent analyses. (A) DeepSeek model-selection benchmark against a fixed 26-pair benchmark set comprising 18 positive executor and regulator pairs and 8 form-mismatch negative controls: the baseline deepseek-chat model reached 81% combined accuracy with 38% false positives; chain-of-thought deepseek-reasoner reached 73% with 25% false positives; and the governance-augmented tool-call loop with a hand-curated reference, together with the self-provided chain-of-thought loop, each reached 100% combined accuracy with zero false positives; the self-provided loop was adopted because it requires no external reference. (B) Consensus query-formulation benchmark against the same 26-pair benchmark set: the form-biased query reached 92% combined accuracy with 12% false positives, whereas the neutral query, the gene symbol followed only by “cell death”, reached 100% combined accuracy with zero false positives. (C) Elicit post-export governance benchmark on the 784 records extracted across three forms (393 ferroptosis, 390 cuproptosis and 1 triaptosis): allow-list compliance improved from 75.4% to 100%, the unsupported EXECUTOR rate fell from 53% to 0%; after governance no unsupported EXECUTOR classification remained (the 192 records were downgraded to REGULATOR when positive regulator evidence (PROMOTES or PROTECTS) was present (∼88%), and to UNCERTAIN otherwise (∼12%)), and publication traceability rose from 85.3% to 98.3% (records with a valid PMID or DOI; descriptive comparison across distinct extraction sets). Green bars denote combined accuracy (panels A and B) or post-governance values (panel C); salmon bars denote false positives (panels A and B) or pre-governance values (panel C).

This benchmark selects the classifier against well-established associations; it is not an estimate of the error rate of the full 21,568-pair classification, whose quality control rests on the self-consistency majority vote, the semantic-contradiction audit, and the governance gates, with exact-string verbatim verification applied to the full-text extracted sentences where available.

### Phase 2G: Consensus Evidence-Channel Classification

A third, fully independent evidence channel was added through the Consensus academic search engine API. Each gene-RCD pair is queried with a neutral formulation, the gene symbol followed only by “cell death”, so that the model retrieves evidence without form-name priming. The channel returns a mechanistic verdict, role, and direction, benchmarked against the 26-pair benchmark set used for the DeepSeek model-selection benchmark; among seven query formulations, the form-biased query reached 92% combined accuracy with 12% false positives, whereas the neutral-query rule reached 100% combined accuracy with zero false positives (Supplementary Note S4; Figure 3B), confirming that form-name priming rather than the retrieval channel itself was the source of false positives. The full extraction runs over all 21,568 unique pairs with a checkpoint file for resumption, subject to a monthly API quota.

### Phase 3 Benchmark: Elicit Systematic Review Governance Validation

#### Rationale

Phase 3 of the RCDome pipeline uses Elicit’s systematic review API (https://elicit.com) to discover gene-RCD associations across 125 million academic papers, including paywalled content inaccessible to PubMed-based retrieval. However, AI-assisted extraction at scale introduces well-characterized risks: vocabulary drift (invention of non-standard terms), hallucination (fabrication of non-existent sentences), and allow-list leakage (extraction of genes outside a predefined target set). To quantify these risks and establish the effectiveness of our governance framework, we conducted a formal benchmark against three RCD forms of varying gene-set size.

#### Benchmark Design

This benchmark quantifies the extraction risks described above and tests whether post-export governance neutralizes them. Three RCD forms were selected to span the range of gene-set complexity in the RCDome matrix: Ferroptosis (1,724 HGNC-validated genes; a large, mechanistically well-characterized form), Cuproptosis (77 genes; a moderately studied, mechanistically distinct form), and Triaptosis (3 genes; a newly recognized form with minimal literature). Each form’s target gene list was extracted from the unified gene-RCD matrix and served as the closed allow-list for that form’s extraction (Figure 3C).

For each form, four complementary Elicit semantic searches were executed to maximize paper discovery while constraining results to manageable batch sizes (50 papers per search): (i) a broad search matching the RCD form name with “regulated cell death mechanism”; (ii) a genetic perturbation search pairing the form with “CRISPR knockout knockdown”; (iii) a pharmacological search for form-specific inhibitors; and (iv) a gene-expression correlation search. Abstract screening was auto-generated by Elicit. Extraction was configured with six structured columns, each with a controlled vocabulary to prevent free-text drift: *`gene_symbol`* (free text, post-filtered against the allow-list), *`mechanistic_role`* (EXECUTOR, REGULATOR, MARKER, UNCERTAIN), *`evidence_type`* (GENETIC_PERTURBATION, PHARMACOLOGICAL, EXPRESSION_CORRELATION, CITATION_ONLY), *`verbatim_sentence`* (direct quote from the paper), *`role_direction`* (PROMOTES, PROTECTS, ASSOCIATED, BIDIRECTIONAL, UNCLEAR), and *`required_for_death`* (YES, PARTIAL, NO, NOT_TESTED, UNCLEAR). Extraction was configured with ‘generate: Fals’ to ensure that the six columns, rather than Elicit’s default auto-generated schema, were applied to every paper.

#### Post-Export Governance

Four automated post-export rules were applied to every extracted record before it entered the benchmark audit: (Rule 1) Allow-list enforcement: any gene symbol not present in the pre-specified form-specific allow-list was rejected with a logged action. (Rule 2) Role normalization: non-standard role terms including SENSITIZER, RESISTANCE, and PROTECTIVE were mapped to REGULATOR, with the corresponding direction inferred (SENSITIZER -> PROMOTES, RESISTANCE/PROTECTIVE -> PROTECTS). (Rule 3) EXECUTOR validation: any record classified as EXECUTOR that lacked ‘*`required_for_death`* = YES’ was downgraded to REGULATOR only when its role_direction indicated PROMOTES or PROTECTS (positive regulator evidence), and otherwise reclassified as UNCERTAIN, reflecting the NCCD criterion that executioner genes must be demonstrably necessary for the death process. (Rule 4) Deduplication: records were deduplicated on (DOI, *`gene_symbol`*, first 50 characters of *`verbatim_sentence`*). Rules 2 and 3 together implement the governance principle that SENSITIZER and RESISTANCE are not ontological categories distinct from REGULATOR but rather directional modifiers of regulation. This normalization reduces a five-term vocabulary (EXECUTOR, REGULATOR, MARKER, SENSITIZER, RESISTANCE) to three, improving cross-form comparability without loss of granularity -- the directional information is preserved in the separate *`role_direction`* column.

#### Five-Gate Evaluation Framework

The benchmark assessed five independent compliance gates, each with a pre-registered threshold: The five Elicit governance gates (EG1–EG5) are defined in Supplementary Table S8. All five gates must pass for the benchmark to validate Phase 3 methodology for production deployment. The thresholds were set prior to data collection and were not adjusted post hoc. Governance actions (normalizations and downgrades) are reported per form and logged in the normalization audit trail.

#### Staged Elicit Extraction and Priority Tiers

Phase 3 Elicit systematic reviews are executed in three priority (P) tiers defined by the number of allow-list genes per RCD form in the unified matrix: P1 (≥100 genes; 20 forms), P2 (25–99 genes; 14 forms), and P3 (1–24 genes; 18 forms). A post-run reconciliation against the 53-form registry identified “Regulated Necrosis” as the omitted 53rd form, which was subsequently executed as a fourth batch (P4). Prioritization by gene count is justified because the allow-list size sets the gene-discovery ceiling per extraction: a form with a larger allow-list can yield more gene-RCD associations per Elicit review, and the largest forms (e.g., Apoptosis, 5,897 genes; Ferroptosis, 1,724 genes) carry the most associations awaiting independent validation.

Each form uses four semantic searches (broad mechanism, genetic perturbation, pharmacological, expression correlation), each at the Pro-plan ceiling of maxResults=1,000 results per search (the documented Elicit Pro limit; see Results, Search Depth Determination), with six governed extraction columns: *`gene_symbol`*, *`mechanistic_role`*, *`evidence_type`*, *`verbatim_sentence`*, *`role_direction`*, and *`required_for_death`*. Extraction proceeds in three priority phases (P1, P2, then P3), processing the largest allow-list first. Execution is sequential, resume-safe (progress saved after every form), and quota-aware: if the monthly API quota is exhausted mid-run, the pipeline halts gracefully after three consecutive session failures, preserving completed forms for resumption after quota replenishment.

#### Governance Literature Survey

To establish that RCDome’s governance safeguards reflect the broader scientific consensus rather than internal convention, we conducted a deep RAG of the AI-governance literature. A corpus of 1,500 papers was assembled by Europe PMC search and reference/citation snowballing, filtered to titles matching governance topics. Each paper was processed by DeepSeek (temperature 0) in a combined-pass extraction returning, per text chunk, governance principles, hallucination-control methods, and benchmark methods, every item anchored to a verbatim sentence copied from the source; items were deduplicated and synthesized into recurring themes, each carrying its supporting PMIDs.

RCDome integrates all 53 RCD-related entities from 20 association databases and resources into a single, queryable resource encompassing 9,265 HGNC-validated human genes and 22,473 individual gene-RCD claimed associations. The number of gene-RCD associations contributed by each source database ranged from 2 (Elicit Alert) to 10,131 (Rodrigues de Souza et al., 2025) (Supplementary Figure S1A). Every association is traceable to its source database and systematically evaluated against the published literature through an automated evidence-scoring framework that assigns confidence tiers (CT1–CT4). At its core lies a curated controlled vocabulary of 53 RCD-related entities, consolidated from 62 original name strings across 20 association databases and resources through systematic literature and database surveillance. Supplementary Table S9 compares RCDome against FerrDB v3 and RCDdb across the resources’ key features. The complete codebase is available for reproducibility and independent audit of every step from database extraction to evidence verification.

### AI-assisted mechanistic description, figure generation, and multi-model quality control

Following selection and ontological adjudication of the RCD-related entities, mechanistic information for each entity was systematically collected using Consensus in Deep mode. A standardized query framework was applied to reconstruct the available molecular evidence from pathway initiation to terminal cell death, including initiating signals, molecular regulators, participating genes and proteins, signaling intermediates, protective mechanisms, execution processes, morphological features, experimentally demonstrated crosstalk, and unresolved mechanistic steps. The framework required distinction between established mechanisms, context-dependent observations, and unresolved or proposed mechanisms, and prioritized traceable evidence from the scientific literature (Supplementary Prompt S1). The complete mechanistic-reconstruction framework is provided in Supplementary Prompt S1 rather than reproduced here.

For each RCD-related entity, the resulting mechanistic description and its associated references were retained together with the corresponding bibliographic Research Information Systems (RIS) file. The extracted text was subsequently harmonized using a second standardized prompt (Supplementary Prompt S2). This step was restricted to organization and linguistic standardization of the supplied material and was not intended to introduce additional mechanisms, molecules, or references. The harmonization framework explicitly required preservation of mechanistic uncertainty, separation of established from emerging evidence, and retention of the references supporting the source description. Each harmonized description was saved as an entity-specific plain-text (.txt) file, named according to the corresponding RCD-related entity, while the associated reference record and RIS file were retained as provenance artifacts.

The harmonized entity-specific descriptions were subsequently used as direct inputs for generation of mechanistic illustrations with ChatGPT (OpenAI; GPT-5.6 Sol) using a standardized figure-generation framework (Supplementary Prompt S3). The prompt imposed common scientific-illustration constraints across entities, including biologically ordered pathway representation, explicit molecular directionality, appropriate subcellular localization, consistent molecular labeling, and a publication-oriented biomedical illustration style. Importantly, the mechanistic description supplied for each individual entity constituted the biological input to the figure-generation step, whereas the standardized prompt controlled the graphical representation and visual organization.

Each initial figure underwent a sequential multi-model quality-control workflow using the standardized audit framework provided in Supplementary Prompt S4. In the first audit stage, the mechanistic description and corresponding figure were evaluated with Claude (Anthropic; Sonnet 5.5) for conceptual and mechanistic inconsistencies, including errors in causal sequence, molecular identity or localization, activation/inhibition directionality, pathway crosstalk, labeling, visual representation, and omission of biologically relevant components. The audit generated explicit element-level corrections rather than a general assessment. The resulting audit report was returned to ChatGPT, together with the original mechanistic description and figure, and used to generate a corrected version.

The revised figure was then subjected to an independent second-stage audit with Gemini (Google; 3.6 Reasoning) using the same quality-control framework. This step evaluated whether previously identified errors had been corrected and whether residual conceptual, mechanistic, directional, labeling, or graphical inconsistencies remained. Figures for which additional issues were identified were returned to ChatGPT for another correction cycle and subsequently reassessed with Gemini. This iterative correction–reaudit loop was continued until no additional correction was requested under the audit criteria. Throughout the workflow, AI-generated outputs were used for literature synthesis, graphical construction, and quality control rather than as primary evidence; the entity-specific mechanistic descriptions, their cited references, and the corresponding RIS records were retained as the traceable evidence layer.

#### Extraction Scripts

The complete extraction-pipeline scripts are catalogued in Supplementary Table S10. For completeness, the 6 omics annotation resources (TCGA, GEO, CPTAC, CCLE, GDSC, DepMap) and the 2 PPI annotation resources (STRING, BioGRID) are annotation resources, not gene-extraction sources.

Each script is self-contained, documented with input/output paths and source URLs, and reproducible with Python 3.8+ and standard dependencies (openpyxl, csv, json, urllib, subprocess, xml.etree, zipfile).

All extraction scripts are available in the code/extraction/ directory and are executable via the master orchestrator *`master_extract.py`*. The output CSVs in output databases/ contain all gene-RCD associations with full source traceability.

## RESULTS

### The RCDome Compendium Composition

The final canonicalized vocabulary comprises 53 RCD-related entities (Supplementary Note S5) spanning five ontological levels; importantly, canonicalization denotes vocabulary normalization and does not confer modality status, and only 19 of the 53 entities satisfy the BG0–BG6 hard-gate criteria for classification as bona fide RCD modalities. The 53 entities are organized into: 19 RCD modalities (bona fide cell death mechanisms with distinct molecular machinery, admitted through the BG0-BG6 hard gates), 7 subtypes (mechanistic subdivisions of parent modalities), 2 triggers (events that initiate cell death without themselves being death mechanisms), 13 functional/contextual categories (contextual programs, morphological outcomes, cell-type and differentiation programs), and 12 generic/umbrella/taxon concepts (umbrella, legacy, taxon-specific, and speculative emerging terms). All 53 entities carry gene-level evidence in the unified matrix (Dataset S1).

The compendium contains 22,473 gene-RCD association rows spanning 9,265 unique HGNC-validated human gene symbols and 53 RCD-related entities (62 original RCD form name strings, consolidated through 9 synonym-merge groups (11 name eliminations, 2 split names)) (Dataset S1). Because the matrix preserves every source-database claim, the 22,473 rows correspond to 21,568 unique gene-RCD associations plus 905 rows of genuine multi-database corroboration, retained as separate rows rather than consolidated because each source may legitimately assign a different mechanistic classification to the same gene-RCD association (e.g., Phase 2A/D versus Phase 2D verdicts); consolidating them into a single row would overwrite the source-specific classifications. Byte-identical duplicate rows arising from a single-source extraction artifact (3,874 XDeathDB key-gene entries) were removed as carrying no additional information, with one copy of each unique record retained for full traceability. Ten further rows for Calcioptosis, a form formally excluded by the five-criteria admission framework (see Materials and Methods), were removed, eliminating three genes (APMAP, ATP11C, CDC50A) that appeared only under that form. Gene biotypes are predominantly protein-coding (20,642, 91.9%), with non-coding RNA genes contributing 1,831 non-coding entries: 1,177 miRNAs (5.2%), 509 long non-coding RNAs (2.3%), 81 pseudogenes (0.4%), and 64 other non-coding RNAs (0.3%); the 1,836 rows that previously lacked a gene biotype (1,832 MSigDB-only plus 4 Elicit-only) were resolved via HGNC mapping, leaving no empty gene biotype.

Among the 53 RCD-related entities, gene coverage is highly heterogeneous: the largest forms, Apoptosis (6,229 association rows; 5,897 unique genes) and Autophagy (4,790 association rows), contain the most gene associations, followed by Ferroptosis (1,811) and Programmed cell death (1,710), while niche forms such as Mitoxyperilysis (13), Triaptosis (3), Pyronecrosis (1), Methuosis (2), and Spectosis (2) have fewer than 20. The median form contains 61 gene entries. Three forms contributed by the external audit (Reactome pathway-specific gene sets: Regulated Necrosis, 59; WikiPathways: Senescence and Autophagy in Cancer, 105; and XDeathDB: Mitotic death, 216) represent newly integrated categories not present in the original NCCD 2018-derived vocabulary. Every gene-RCD association carries full provenance to its source database, ten RCD-specific curated databases (FerrDB v3, RCDdb, iPCD, LncRCD, iLIR, HADb, HAMdb, ATD, AMT-DB, ncFO), five pathway and molecular databases (KEGG, Reactome, Gene Ontology, MSigDB/GSEA, WikiPathways), NCBI Gene, and six omic-annotation resources (TCGA, GEO, CPTAC, CCLE, GDSC, DepMap), supplemented by the AI-assisted Elicit extraction (20 association databases and resources, Supplementary Table S2), and is linked to a seven-class mechanistic classification derived from the Phase 2A/D independent LLM validation pipeline. Most associations rest on a single source (10,009 of 22,473 rows), 6,313 are corroborated by two sources, and corroboration declines steeply with additional sources; the long tail up to 14 sources marks the best-corroborated associations, such as the canonical executors (Supplementary Figure S1B).

Read at the level of mechanism, the compendium reveals a remarkably small execution core: only 225 gene-RCD pairs (4.7% of the role-resolved set) are executors, genes that directly pore, cleave, or permeabilize to kill the cell, while 4,455 pairs (93.3%) are regulators that promote or inhibit death without executing it (Figure 4A). Within the compiled RCDome association universe and under the evidence-first adjudication, the mechanistically involved set is dominated by regulatory associations surrounding a comparatively small executor subset.

**Figure 4.**
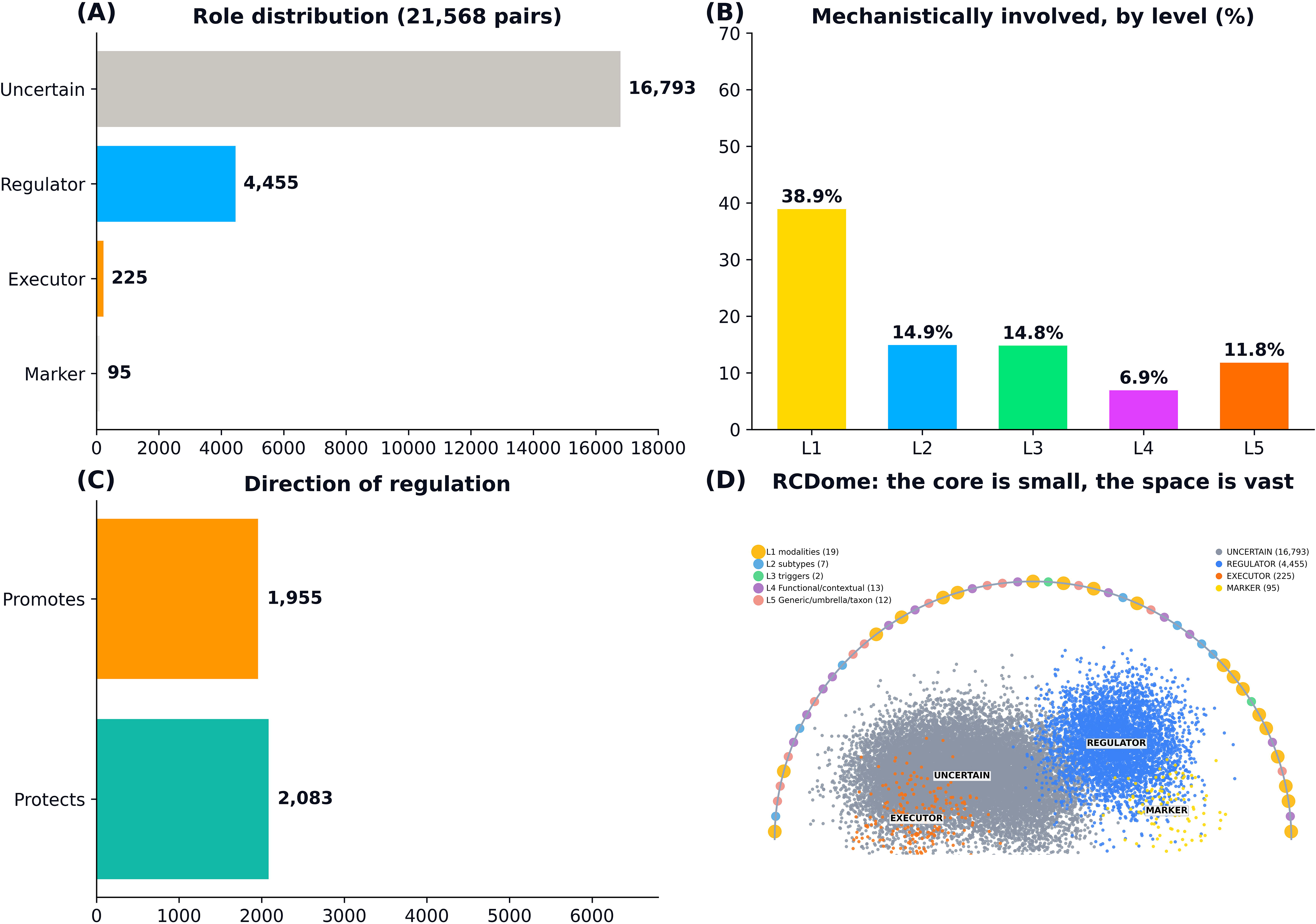
The RCDome: death core and regulatory network. (A) Role distribution across the 21,568 unique gene-RCD pairs classified by the evidence-first adjudication: 16,793 unresolved, 4,455 regulators, 225 executors, and 95 markers. (B) Mechanistically-involved fraction by ontological level (L1 modalities, L2 subtypes, L3 triggers, L4 Functional/contextual categories, L5 Generic/umbrella/taxon concepts); the modalities carry the densest mechanistic signal (38.9%) while the Functional/contextual categories (L4) are the most dilute (6.9%), with the Generic/umbrella/taxon concepts at 11.8%. (C) Direction across all role-resolved associations: 1,955 promote and 2,083 protect (among the 4,455 regulators, 1,730 promote and 2,083 protect), a near-even balance. (D) The RCDome: each dot is one association, drawn in role-colored clouds whose dot density reflects the count — the unresolved mass (grey, 16,793), the regulatory cloud (blue, 4,455), and the tiny executor (orange, 225) and marker (gold, 95) clouds — set inside a dome whose rim carries the 53 canonicalized RCD-related entities as colored nodes (L1 modalities larger, L2–L5 smaller), illustrating that the death-executing core is small and the space is vast.

### The RCD entity census and operational definitions

The RCDome identified 53 canonicalized RCD-related entities, represented by 22,473 association rows (21,568 unique gene-RCD pairs) in the unified evidence matrix. Of these, 20 carried the verbatim operational definitions of the NCCD (Galluzzi et al., 2018), 12 carried the expanded operational definitions of the multi-optosis dataset (Rodrigues de Souza et al., 2025), of which five (cuproptosis, disulfidptosis, erebosis, alkaliptosis, and oxeiptosis) were not described by the NCCD and had been AI-extracted from a 6,603-PDF corpus and human-validated, and 20 were drafted de novo by the RCDome (19 founding-draft plus one pre-PubMed term) and substantiated by their founding articles. One additional record (“Senescence and Autophagy in Cancer”) was a composite WikiPathways pathway label and was retained, per the provenance policy, with the evidence PMIDs of its two parent entities rather than a fabricated PMID of its own. Thirteen entities carried both an NCCD and a Rodrigues et al., 2025 definition; seven NCCD forms are represented in the matrix as finer-grained entities that Rodrigues et al., 2025 had grouped under a parent term. (Supplementary Table S11) The ontological adjudication of each entity, the pre- and post-adjudication levels and the Q1–Q13 evidence trail under the BG0–BG6 hard gates, is documented in Supplementary Table S12.

Among the 25 multi-optosis-anchored entities the overall score S ranged from 9.70 (apoptosis) to 4.15 (necrosis), and among the 28 newly-added entities from 7.20 (programmed cell death, an umbrella term) to 3.25 (vacuolar cell death). Across all 53 entities the distribution was right-skewed (mean 5.20, median 4.85, standard deviation 1.48, skewness +1.09): the drafts occupied the lower portion of the calibrated scale (mean 4.41 versus 6.09 for the anchors), consistent with the predominance of emerging, taxon-specific, legacy, and contextual forms among the newly defined entities.

Each entity’s operational definition is a 42-field record combining the definition, scoring, and provenance layers. For example, the eryptosis record comprises: the operational definition (“A cell-type-specific form of regulated cell death of erythrocytes, demarcated by cell shrinkage, membrane blebbing, calpain activation and phosphatidylserine exposure, and triggered by oxidative stress via prostaglandin E2/Ca2+ and phospholipase A2/ceramide pathways”); the key molecules (Ca2+ channels, ceramide, calpain, erythropoietin); the inflammatory status (no); the overlap description (phosphatidylserine-exposure clearance, apoptosis/efferocytosis-like); the six criteria RS1–RS6 (4/4/4/4/5/5) with overall score S = 4.15; and the full provenance chain (founding (Lang et al., 2006); authoring source (Rodrigues de Souza et al., 2025), RCDome; source databases Elicit_Phase3 and NCBI_Gene; channels B and E).

All 53 RCD entities were anchored to a reference PMID. All 52 verifiable founding PMIDs resolved in PubMed (Knock Test). Verbatim integrity was confirmed for 20/20 NCCD terms and 25/25 multi-optosis records. Consensus cross-validation corroborated each de novo definition against coinage, mechanism, and PubMed-consensus evidence streams, yielding 13 fully corroborated and 13 partially corroborated drafts — the 19 de novo FOUNDING_DRAFT definitions plus 7 NCCD-verbatim entities with draft complements (subtypes, umbrellas, and legacy terms without a single coining paper), plus one NCCD-locked definition whose draft complement only was unscored, for 27 of the 53 entities. DeepSeek full-text validation of the 33 non-NCCD entities (32 not in the NCCD 2018 source plus the composite pathway label), using PMC full text where available and Consensus full-text-derived takeaways otherwise, returned 6 fully validated, 9 with minor wording gaps, 4 criterion-pending, 1 umbrella-scope, and 1 abstract-limited (the composite pathway label); the remaining 12 were verbatim-locked Rodrigues definitions. Eight entities were initially flagged, of which six were verbatim NCCD/Rodrigues et al. (2025) definitions flagged only by a source-mismatch (the committee text had been checked against the founding paper rather than the committee source) and were therefore reclassified as verbatim-invariant; the remaining genuine flags are the criterion-pending and umbrella-scope drafts described below.

Four of the de novo drafts, lysoptosis, PANoptosis, regulated necrosis, and triaptosis, were flagged as criterion-pending because their founding articles describe the mechanism but not the MC2 trigger and/or the MC3 morphological phenotype required by the NCCD definitional logic; these elements remain to be characterized in the primary literature. Metalloptosis was flagged as umbrella-scope: like mitotic catastrophe and cellular senescence in the NCCD itself, it is an umbrella category rather than a discrete RCD form and therefore lacks a single trigger and morphology by definition. These flags are recorded as explicit definition-status annotations rather than as errors.

Two ontology corrections were applied following the biological curation: MPT-driven necrosis was reclassified from the trigger level (L3) to the modality level (L1), while the mitochondrial permeability transition (MPT) itself remains a trigger (L3), correcting a trigger-centered placement; and vacuolar cell death was reclassified from L4 to L5 as a taxon-specific morphological category (plant + protist; not human). Following the adoption of strict hard gates (BG0-BG6; Materials and Methods), the ontological levels were recalibrated across the 53 entities: L1 was restricted to the 19 bona fide modalities, while morphology, umbrella, contextual, taxon-specific, legacy, and speculative emerging forms were re-leveled to L2-L5 (Supplementary Table S11).

### Compendium Completeness

Following the consolidation, 50 of 100 matrix columns are populated with a non-empty value in all 22,473 rows (i.e., applicable to every association). This completeness profile reflects the multi-source architecture rather than a compilation gap: the fully populated columns cover gene identity, ontological placement, and classification, whereas the remaining 50 columns are source-specific evidence fields that were filled only where the corresponding source contributed evidence. Gene biotypes, Ensembl IDs, and UniProt IDs were mapped from the HGNC complete set: gene biotype is complete for all 22,473 rows; Ensembl IDs are set to NO_VALUE_FOUND for 38 rows (33 loci) that lack an HGNC Ensembl cross-reference; UniProt IDs are present for all protein-coding rows and set to NOT_AVAILABLE for 1,699 non-protein-coding genes. Eight Elicit-derived ontology columns (rcd type, parent relationship, operational notes) classify all 53 RCD-related entities into an evidence-anchored taxonomy with literature-justified operational notes. Five multi-optosis operational definition columns provide PMID-referenced definitions for the 25 forms across the 9,188 genes associated with those forms; the remaining 77 genes belong to entities outside the multi-optosis ontology and carry no multi-optosis definition. Ancillary annotation columns use the sentinel value NOT_AVAILABLE for non-protein-coding genes. The Europe PMC recovery pipeline (see Materials and Methods) populated 22,187 of 22,473 rows (98.7%) with at least one traceable PMID across four evidence channels (pmid_list, llm_pmids, epmc_pmids, elicit_pmids). A targeted Phase 2E extension searched Europe PMC full-text with DeepSeek LLM for all pairs lacking a co-occurrence PMID; none showed additional evidence, confirming these 286 DATABASE_ONLY (1.3%) represent the irreducible uncertainty of RCD knowledge: gene-RCD associations from curated databases with no independent verification in the indexed literature. rcd_gene_type and pmid_list remain empty only for database-only rows outside the Elicit and deep-RAG layers (17,578 and 20,030 of 22,473, respectively). No critical analytical column is empty. A formal governance contract with matrix provenance protocol ensures traceability, reproducibility, and auditability of the compendium. Separately, 19,410 rows (86.4%) were classified as non-MECHANISTIC by Phase 2A/D, prompting the development of a full-text recovery pipeline to identify hidden evidence in the literature. As a refinement of the retrieval strategy, this pipeline replaces PubMed’s gene-indexed search with Europe PMC’s full-text search engine.

### Phase 2A/D Validation: Scale and Coverage

Validation was performed on all 21,568 unique gene-RCD combinations. Of the compendium’s 22,473 total rows, 22,187 carry PMIDs (98.7%), corresponding to 21,283 of these 21,568 unique combinations. Each was independently assessed through a two-call LLM architecture against PubMed abstracts.

### Validation Performance

The two-call majority-vote architecture reclassified evidence tiers, and the *`mechanistic_class`* versus *`evidence_tier`* concordance was 79.2%. The resulting tier distribution is reported below. Up to the first 100 retrieved PMIDs per pair used for content verification are retained in the unified matrix, and the full retrieved paper count is stored separately as the evidence force.

### Evidence Tier Distribution

After Phase 2A/D validation, the unified matrix evidence tiers (as defined in the Materials and Methods section, Evidence Tier Assignment) were distributed across the 22,473 matrix rows as follows:

ET1 (Gold Standard), 2,864 rows (12.7%), reflects multi-database consensus with RCDdb PMID support

ET2 (Strong Evidence), 5,686 rows (25.3%), reflects three or more independent sources or LLM-confirmed

ET3 (Moderate Evidence), 8,212 rows (36.5%), reflects two sources with LLM partial confirmation

ET4 (Weak Evidence), 5,711 rows (25.4%), reflects single source without LLM confirmation, or LLM unable to verify

### Mechanistic Classification

While evidence tiers classify the strength of support for each gene-RCD association, the mechanistic classification independently categorizes the nature of the association by combining LLM status with quantitative evidence force. The following seven-class classification resolves the ambiguity of raw LLM outputs.

Two measures describe mechanism and are not interchangeable. The evidence-first adjudication labels a pair mechanistically involved when the evidence supports a mechanistic role, giving 23.8%. The stricter seven-class scheme reserves the label MECHANISTIC for rows assessed by both LLM knowledge and PubMed abstracts in the abstract-level pass, giving 13.6%, and full-text recovery adds further mechanistic support in later phases. The difference is the stringency of the criterion, not a disagreement between the analyses. The per-row verdict shown in the Gene Catalog module of the RCDome Shiny web application (https://rcdome.org/) is the seven-class label; the 23.8% figure is an aggregate summary of the evidence-first adjudication.

Applied to all 22,473 validated rows, the seven classes are:

MECHANISTIC, 3,063 rows (13.6%), assessed by both LLM knowledge and PubMed abstracts; LITERATURE_GAP, 1,519 rows (6.8%), LLM training recognizes but abstracts lack explicit statements;

CONTEXT_GAP, 6,170 rows (27.5%), PubMed confirms but LLM training does not recognize; CO_OCCURRENCE, 2,416 rows (10.8%), high paper counts but no mechanistic evidence; INSUFFICIENT_DATA, 9,019 rows (40.1%), few papers; LLM cannot determine involvement; DATABASE_ONLY, 286 rows (1.3%), gene and RCD term never co-occur in PubMed; TECHNICAL_FAILURE, 0 pairs (0%), no API failures in the final run.

Association record here means that a database or a search recorded the gene and the death form together; it does not by itself establish a mechanistic role. Rows classified DATABASE_ONLY or CO_OCCURRENCE are association records without mechanistic support, and the web application displays them as such so they cannot be mistaken for curated mechanism.

Together, the seven mechanistic classes account for all 22,473 validated rows.

Phase 2A/D validation of the 1,832 MSigDB-only gene-set pairs (predominantly the ‘Programmed cell death’ gene set, n=1,512) assigned mechanistic classifications and PMIDs to every pair. Of these, 282 were confirmed mechanistic, 265 regulators (155 pro-survival, 110 pro-death) and 16 executors, while 150 were literature gaps, 471 context gaps, 428 co-occurrence, 316 insufficient-data, and 185 database-only. Overall, 1,576 of the 1,832 pairs (86.0%) now carry at least one PMID.

### Phase 3 Benchmark Results

#### Extraction Summary

The Elicit systematic review pipeline extracted gene-RCD associations for all three benchmark forms in a single automated run. Across the three forms, 154 unique papers yielded 784 records before post-export governance, comprising 393 Ferroptosis records, 390 Cuproptosis records, and 1 Triaptosis record (Supplementary Table S13). The low yield for Triaptosis reflects its status as a newly named RCD form (2024) with only three genes currently recorded in the RCDome matrix and minimal published literature; this result is consistent with the form’s emerging evidence grade C (on the A-D evidence-grade scale: A, well-established; B, established; C, emerging; D, proposed).

The raw and post-governance record counts are equal (784) because the post-export governance normalizes roles (192 EXECUTOR downgrades (to REGULATOR or UNCERTAIN)) rather than removing records; zero records were rejected by the allow-list filter.

#### Gate Results

All five Elicit governance gates passed at their pre-registered thresholds (Supplementary Table S14). Allow-list compliance was 100%: no gene symbol outside the pre-specified form-specific lists entered the final record set, demonstrating that the post-export allow-list filter fully compensates for Elicit’s tendency to extract genes beyond the instructed scope. Vocabulary compliance was 100%: all 784 records used only the permitted role vocabulary (EXECUTOR, REGULATOR, MARKER, UNCLEAR), with zero instances of non-standard terms such as SENSITIZER, RESISTANCE, or Suppressor entering the final audit. No unsupported EXECUTOR classification remained: the 192 records initially classified as EXECUTOR by Elicit were downgraded to REGULATOR when positive regulator evidence (PROMOTES or PROTECTS) was present (∼88%), and to UNCERTAIN otherwise (∼12%) by the ‘*`required_for_death`*’ validation rule, leaving zero unvalidated EXECUTOR claims in the final record set. This result is structurally expected: Elicit’s extraction engine tends to classify any gene with published experimental manipulation as an EXECUTOR; the governance rule corrects this by requiring explicit evidence of death requirement. Traceability was 98.3% (770/784 records with a valid PMID or DOI), reflecting Elicit’s native use of DOIs as paper identifiers.

#### Governance Impact

The 192 EXECUTOR downgrades (88 Ferroptosis, 104 Cuproptosis) represent the single most high-impact governance action. Without the ‘required_for_death’ validation rule, the benchmark would have reported 192 EXECUTOR claims, none of which met the NCCD criterion of demonstrable necessity for the death process. This finding is consistent with the cuproptosis pilot audit, where 53% of EXECUTOR classifications were inflated, and validates the utility of embedding NCCD-compliant definitions directly into the extraction governance layer rather than relying on post hoc expert review.

#### Scalability Assessment

The benchmark extracted 784 governed records from 154 papers. Production search depth was determined empirically (see Search Depth Determination below) rather than extrapolated; the measured Ferroptosis scaling series from 50 to 1,000 results per search is reported in Supplementary Table S15. The benchmark validates the methodology for production deployment; the scaling strategy prioritizes the P1 forms (those with 100+ genes in the matrix) for full extraction, with P2 and P3 forms following as capacity allows.

#### Comparison with Pre-Governance Baseline

The Cuproptosis FULL pre-fix review (372 papers, 1,143 records, processed before governance rules were implemented) serves as a baseline for measuring governance impact. In that pre-governance extraction, only 75.4% of records complied with the allow-list, 53% of EXECUTOR classifications lacked *`required_for_death`* evidence, and 85.3% of records had a traceable PMID or DOI. The benchmark demonstrates that post-export governance eliminates all three failure modes simultaneously: allow-list compliance rose from 75.4% to 100%; the unsupported EXECUTOR rate fell from 53% to 0%; and traceability (records with at least one valid PMID or DOI) rose from 85.3% in the baseline to 98.3% in the governed benchmark (Figure 3C). Because the baseline and the governed benchmark comprise distinct extraction sets, these percentages are reported as a descriptive comparison between separate executions rather than a paired, controlled estimate of the governance effect. The governance framework does not merely improve extraction quality; it transforms an unreliable AI extraction into a verifiable, auditable evidence pipeline.

#### Search Depth Determination

To determine the production search depth, we measured how Elicit’s paper yield scales with the maxResults parameter across a five-point series using Ferroptosis (1,724 allow-list genes) as the test form. Elicit’s API documentation specifies a default of 200 results per search (valid range 1-10,000) subject to plan-specific limits; the Pro plan used here caps results at 1,000 per search and 5,000 total per review. Omitting maxResults therefore does not produce an uncapped search -- it silently applies the 200-result default. Elicit’s semantic search uses vector-similarity retrieval, returning the top-N papers in descending relevance to the query.

The series reveals steeply diminishing returns. The marginal gene yield declines monotonically with each depth increment: 50 to 100 adds 85 genes over 81 papers (1.05 new genes per paper), 100 to 200 adds 116 genes over 167 papers (0.69), 200 to 500 adds 212 genes over 561 papers (0.38), and 500 to 1,000 adds 157 genes over 852 papers (0.18). The default (200) captures 424 unique genes from 315 papers (1.35 genes per paper), while the Pro ceiling (1,000) captures 793 genes from 1,728 papers (0.46 genes per paper). Gene discovery therefore does not scale linearly with search depth; marginal papers increasingly re-confirm already-known associations. Production extraction was configured at maxResults=1,000 (the Pro ceiling) to maximize recall within plan limits, while noting that the default of 200 already captures 53% of the ceiling’s gene yield (424/793) at 18% of the paper count (315/1,728).

The Elicit literature-mining layer yielded 4,601 gene-RCD associations across 50 RCD forms, triaged into 3,803 genuine (each carrying a body-level verbatim sentence and a DOI, with a PMID in 98.3% of cases) and 798 association_only (database-backed associations whose Elicit classification was not retained). The three forms without Elicit results were out of scope by design: Erebosis (Drosophila), Hypersensitive response (plant), and Regulated Necrosis (an umbrella term). Among the 3,803 genuine rows, the functional classification was dominated by regulators: REGULATOR 3,347; EXECUTOR 323; MARKER 119; UNCERTAIN 10; CANDIDATE_MODULE 3; and 1 unassigned. Direction of effect was predominantly PROMOTES (2,397) versus PROTECTS (963), with 1,392 genes required for death. Evidence type: GENETIC_PERTURBATION 2,079; EXPRESSION_CORRELATION 801; PHARMACOLOGICAL 670; CITATION_ONLY 3; and 250 (6.6%) unassigned, where the extraction model left the field blank because no clean evidence type was determinable from the source abstract.

Phase 2B processed 167,868 free PMC full-text articles across 21,283 gene-RCD pairs carrying at least one PMID, yielding deep evidence for 1,383 rows (1,118 unique pairs). Deep evidence is full-text, sentence-level support: for each pair, the pipeline recorded the papers examined (open-access versus paywalled), the strongest verbatim body sentence, the functional role, the evidence type, the direction of effect, a multi-paper confidence tier, and the proportion of papers with direct experimental support. Functional roles were dominated by regulators (878), followed by markers (297), interactors (88), executors (78), and substrates (42). Evidence strength was expression-correlation (529), genetic intervention (371), citation-only (304), direct experimental (139), and pharmacological (40). Confidence was low for 821 pairs, medium for 366, and high for 196. Direction was promoting (492) versus inhibiting (390), with 430 unknown, 70 context-dependent, and one row without a recorded direction. Because deep evidence only upgrades or confirms, never downgrades, it promoted 142 gene-RCD pairs from a moderate to a strong evidence tier and elevated 131 pairs from a literature-gap to a mechanistic classification. These promotions are conservative by design: the deep layer strengthens a claim only when the full-text body independently supports it, leaving all weaker or uncertain classifications untouched.

Phase 2C full-text Europe PMC recovery added 3,058 MECHANISTIC classifications, in addition to the 3,063 MECHANISTIC rows from Phase 2A/D abstract validation. Across the two evidence layers, the deep full-text pipeline and the Elicit systematic review corroborated 387 gene-RCD pairs jointly, out of 4,863 unique pairs covered by at least one of the two layers: 1,118 pairs had deep full-text evidence and 4,132 had Elicit evidence. The two layers drew on nearly disjoint literature, with 1,013 unique full-text papers in the deep layer versus 8,253 PMIDs and 9,146 DOIs in the Elicit layer, totaling 10,138 unique papers. Only 21 papers were cited by both layers, confirming that the 387 jointly corroborated pairs rest on independent rather than redundant literature. Overall, the AI evidence layers covered 4,169 gene-RCD pairs (19.3% of the compendium’s unique pairs) with genuine verbatim support, rising to 4,863 (22.5%) when association-only claims are included.

The 387 jointly corroborated pairs were strongly enriched for high confidence: 211 (54.5%) carried the strongest evidence tier and 235 (60.7%) were classified as mechanistic, with 145 (37.5%) meeting both criteria simultaneously. Functional roles were dominated by regulators (243; 62.8%), followed by markers (79), executors (33), interactors (23), and substrates (9). Most (334; 86.3%) were genuine Elicit findings, with 53 (13.7%) association-only. Dual-layer agreement therefore concentrates the highest-confidence, mechanistically characterized links: a 4.3-fold enrichment of strong-tier and a 4.5-fold enrichment of mechanistic pairs relative to the compendium as a whole.

#### Governance Literature Survey Results

The survey recovered 27,852 verbatim-anchored items from 1,308 papers, synthesized into 419 themes, of which 255 were AI-governance, 71 human-clinical (excluded), and 93 ambiguous. The AI-relevant literature converged on five recurring principles, evaluation and benchmarking, detection and mitigation, human oversight and accountability, transparency and explainability, and ethics and regulation, all of which RCDome implements (exact-string verification, curator flags, provenance traceability, the five-gate benchmark, and the governance contract). No single consensus framework exists: 78% of extracted themes appear in a single paper, positioning RCDome’s explicit governance as an early, principled implementation rather than a follower of a mature standard. The full ranked governance framework (51 themes with supporting PMIDs) is provided in the study GitHub repository.

#### Full-Matrix Classification and Evidence-First Adjudication

Chain-of-thought (CoT) classification was applied to the complete matrix to add a mechanistic verdict, functional role, and direction to every gene-RCD pair, information the two-call binary validation (Phase 2A/D) and the full-text recovery phases did not capture uniformly. Rather than a single model answer, each pair received three separately sampled chain-of-thought runs of the same production classifier with a self-governance check (verify-role), merged by majority vote, so that one inconsistent sample could not dominate the final verdict.

The evidence-first adjudication resolved 5,138 (23.8%) of the 21,568 unique gene-RCD pairs as mechanistically involved (YES) and 16,236 (75.3%) as not mechanistically involved (NO). Among the role-resolved pairs, 225 (4.7%) were executors, genes that directly execute the death pathway, and 4,455 (93.3%) were regulators that promote or inhibit it. Across the 21,568 unique gene-RCD pairs, roles resolved as 4,455 regulators, 225 executors, 95 markers, and 16,793 with an unresolved role; direction (across all role-resolved associations) resolved as PROMOTES for 1,955, PROTECTS for 2,083, and UNCLEAR for 737, a near-even pro-death versus anti-death balance. Overall, 4,000 of the 9,265 genes represented in RCDome (43.2%) had at least one mechanistically involved association. The number of mechanistic associations varied markedly by source database, ranging from 1 (Elicit Alert) to 3,872 (Rodrigues de Souza et al., 2025) (Supplementary Figure S2).

The classification was internally audited: 84 self-inconsistent rows (mechanistic NO with a confident role or direction) were repaired, and 231 rows with genuine uncertainty, 178 regulators with unclear direction and 53 mechanistically-involved pairs with unresolved role, were retained as unresolved rather than forced into a confident category. Two gene-RCD pairs (NLRX1 and PPIF with MPT-driven necrosis) were classified separately with a five-vote majority, completing full pair coverage (21,568/21,568).

#### The RCDome Web Application

The compendium is exposed through the RCDome Shiny web application (https://rcdome.org/) with a multi-layer navigation. The Home page renders the atlas as a girded-cupola dome, 53 entity nodes colored by ontological level (L1 gold through L5 orange) along the rim, with a gene constellation beneath; a modules grid on the same page opens the nine data modules. Clicking an entity node opens a detail page showing its operational definition, associated genes (filterable by direction, role, mechanistic class, evidence databases, and PMID availability), and the literature evidence (association PMIDs and co-occurrence force). A jump-to-entity selector permits direct navigation among the 53 entities without returning to the dome.

The dome is the visual signature of the platform: its cupola rim carries the 53 canonicalized RCD-related entities as nodes colored by their adjudicated ontological level, L1 modalities (19, gold), L2 subtypes (7, blue), L3 triggers (2, green), L4 Functional/contextual (13, purple), and L5 Generic/umbrella/taxon (12, orange), over a constellation of the 9,265 HGNC-validated genes. This color-coding makes the ontology’s core claim visible at a glance: only 19 of the 53 entities are bona fide RCD modalities; the remaining 34 are subtypes, triggers, contextual programs, or umbrella concepts, and their level color encodes that distinction.

RCDome Platform Architecture. The RCDome web application is a pure R Shiny application; the compendium matrix is stored as an in-memory R data file (RDS), with no external microservice or database backend. The interface comprises a Home landing page, an Explore RCDome educational guide, and the data modules Operational Definitions, Compare Definitions, RCD entities, Gene Catalog, Discrepancy Radar, Discovery Timeline, the Digital Molecular RCD Board, Network, and 3D RCDome, plus an About page with data provenance. The Operational Definitions module is a browser for the 53 entities’ operational definitions, sourced directly from the RCDome Decoder: a searchable list on the left and the full 42-field record (definition, scoring, and provenance) on the right.

The Compare Definitions module places two to four operational definitions side by side in adjacent columns, so that shared wording and differences are visible at a glance.

The RCD entities module presents the deep-dive pages: the 53 audited descriptions verbatim with functional citation hyperlinks, together with a full-page cartoon and its legend for each entity.

The application source code and compendium are available at https://github.com/BioCancerInformatics/RCDome. The Gene Catalog module exposes the complete 22,473-row × 100-column matrix (Dataset S1) as a filterable, sortable table with ten columns, gene, biotype, RCD entity, cancer gene, confidence tier, mechanistic, role, direction, source databases, and evidence, and a Direction/Role source selector that switches the displayed annotation among the DeepSeek chain-of-thought, Elicit, Consensus, GPT-4o, and evidence-first adjudication sources. Each analysis is preserved in its own columns, so the interface never conflates provenance. A Death core preset filters to the 225 executor pairs; a per-entity comparison table presents Elicit versus DeepSeek executor and regulator counts side by side for all 53 entities; and four download buttons export the filtered table, the complete matrix, the entity comparison, and the death core. https://github.com/BioCancerInformatics/RCDome.

The Discrepancy Radar module exposed 7,905 classifier disagreements — 7,892 with a third-opinion arbitration and 13 delta-adjudicated — (Supplementary Note S2), faceted by the role, direction, and mechanistic axes. Each row was color-coded by its label and by concordance (full agreement, 2-of-3, or discordant) and carried the GPT-4o arbitration verdict with its reasoning. The evidence-first adjudication column reports the resolved value and its decision path.

The Discovery Timeline module plotted the 53 entities on five horizontal lanes, one per ontological level, from 1859 to 2025, each entity placed by its discovery year and colored by level. Clicking an entity opened a discovery card with its discovery year and the founding paper, its title, authors, journal, and clickable PMID and DOI links.

The application also includes a Digital Molecular RCD Board, a prose-synthesis module that answers gene-RCD queries strictly from the RCDome database and the RCDome decoder under the governance rules described in the Materials and Methods section, with a retrieval-augmented Q&A submodule that answers free-form questions across all 53 entities and all genes. A nine-test board benchmark (Supplementary Note S6) validated the board at 100% golden accuracy, 100% census accuracy, 100% fact-stability, a 0.047% governance violation rate, and 98.7% articulation conformity; a two-test Q&A benchmark (n = 100 per test) validated the Q&A at 100% positive accuracy and 100% negative controls, Wilson 95% confidence interval (CI) 96.3–100%, two-sided binomial p = 0.010; a seven-layer reporting benchmark (n = 200 pairs × 7 layers; 1,238 scored) measured 93.2% overall layer accuracy (95% CI 91.7–94.5%) and 100%/98% Evidence-first/DeepSeek precision against cross-layer substitution.

The Network module renders the gene-RCD associations as an interactive network: entity nodes colored by ontological level, gene nodes colored by dominant mechanistic role, with hover details (gene symbol and HGNC name; entity name, level, and definition) and click-through from a gene to its literature evidence.

The 3D RCDome module rendered the atlas as a rotatable three-dimensional sphere: the 53 entities on the shell, colored by level, and the gene constellation inside. By default the constellation displayed the top 400 genes for legibility, with a control to switch to the full 9,265-gene constellation. Network lines connected each gene to its top entity. Clicking a level in the legend highlighted that level’s entities, genes, and lines; clicking an entity opened a focused view of its gene constellation, where each star could be hovered and clicked to open the evidence trail for that gene in that entity.

The interface is built for both education and scientific accountability. Every column, filter, and control carries a hover explanation; pills are colored by concept (level, biotype, tier, role, direction); entity cards are level-colored; and every PMID shown as evidence resolves to a verified article title (10,141 curated titles, 100% accuracy against PubMed). The nine data modules (Operational Definitions, Compare Definitions, RCD entities, Gene Catalog, Discrepancy Radar, Discovery Timeline, the Digital Molecular RCD Board, Network, and 3D RCDome) are exposed as cards on the Home page, with the interactive dome as the primary entry point.

The application was built iteratively with AI assistance (DeepSeek and Consensus classification, AI-generated pathway figures and descriptions) under the governance rules described in the Materials and Methods section. Every value exposed in the interface was verified against the compendium matrix (Knock Test), and every AI-generated element is traceable to its source. Detailed AI usage, token, runtime, and cost figures for every AI-assisted phase are reported in Supplementary Note S7 (AI usage, tokens, and cost).

### Practical utility of the web application

The interface is built for extraction as much as for browsing: in a single session, a researcher can filter to a mechanism of interest, download the underlying matrix, and trace any claim back to its source, with every association PMID resolving to a verified article title.

## DISCUSSION

RCDome was built to answer a single question: “Where did this gene-RCD association come from, and what is the evidence?” Every architectural decision follows from that question. The five-channel architecture ensures that no single source type dominates: Channel A (PubMed Boolean) maximizes recall, Channel B (curated databases) provides precision, Channel C (AI-assisted screening) captures recent discoveries, and Channel D (manual curation) serves as the ultimate arbiter, and Channel E (Elicit systematic review) recovers full-text associations. The two-call independent LLM validation architecture prevents any single model from biasing the verification: one call draws on internal knowledge, the other on PubMed abstracts, and only concordance between them yields a CONFIRMED classification. The five-criteria admission framework ensures that the L1 modality subset is a set of mechanistically distinguishable entities, while the broader 53-entity vocabulary also contains subtypes, triggers, and conceptual categories retained for provenance. Every design choice was made to maximize one thing: traceability.

These design principles are implemented as executable checks in the pipeline. The traceability contract rejects ambiguous patterns (“and others,” “etc.”) at integration. The anti-hallucination framework verifies every LLM-extracted sentence against its source document by exact string matching; sentences not found verbatim are rejected and logged. The *`ontology_curator_verified`* flag was set to TRUE for all 53 entities following individual entity-by-entity review and approval (logged 2026-09-05). Automated quality filters exclude non-HGNC symbols and NCBI Gene keyword-only associations following a spot audit that revealed zero overlap with known RCD executioner and regulator genes, a decision that removed 10,857 rows but protected the compendium from systematic noise. Governance, in RCDome, is not a statement of values; it is executable code. RCDome uses a benchmarked, multi-agent/LLM evidence architecture with human adjudication and executable governance to transform heterogeneous biomedical database assertions into a traceable, literature-validated human gene knowledge atlas.

The RCDome compendium is a diagnostic of the field, and several findings took us by surprise. The sheer number of RCD entities, 53 RCD-related entities consolidated from 62 original name strings, was not something we anticipated when this project began. The list includes 19 bona fide RCD modalities, but also subtypes, trigger events, functional/contextual categories, and generic/umbrella/taxon concepts. Regulated cell death is far more diverse than the textbooks suggest, and the vocabulary continues to expand.

The disparity in gene coverage is striking. Apoptosis accounts for 6,229 association rows (5,897 unique genes). Pyronecrosis has one. This is not a flaw in the compilation; the coverage pattern likely reflects research intensity, field maturity, database representation, and source/indexing depth. Apoptosis has been studied for over fifty years. Several of the most recently described forms are known from a single discovery paper. The compendium makes this imbalance explicit, and it is sobering. The mechanistic classification produced an even more striking result. Of the 22,473 gene-RCD associations, only 3,063 (13.6% of the 22,473 validated rows) are MECHANISTIC, meaning both LLM knowledge and PubMed abstract evidence support a functional role. For the remaining 86.3% of validated rows, the literature is either insufficient, ambiguous, or silent. A further 2,416 rows (10.8%) reflect CO_OCCURRENCE: the gene and RCD term appear together in published articles, but without evidence of a functional relationship. For 9,019 rows (40.1%), the literature is insufficient to determine the nature of the association. An additional 1,519 rows (6.8%) represent a LITERATURE_GAP: the LLM recognizes the association, but PubMed abstracts lack explicit confirmation. In other words, for the vast majority of gene-RCD associations reported across 20 resources, the published record does not yet explain how the gene relates to the cell death form. Finally, 286 rows (1.3%) are classified as DATABASE_ONLY. These are gene-RCD associations contributed by 10 independent curated databases for which no supporting PMID was found in any indexed literature source, despite exhaustive search via NCBI PubMed, Europe PMC full-text recovery, and a targeted DeepSeek LLM campaign. They represent unresolved evidence gaps within the literature sources and retrieval strategy used in this study. They are flagged for expert manual review. Taken together, these findings suggest that the RCD field has outgrown its information infrastructure. Databases have accumulated associations faster than the literature has verified them. RCDome does not solve this problem, but it makes it quantitatively measurable.

RCDome does not replace FerrDB v3, RCDdb, or any of the 20 association databases and resources from which it draws. It integrates their gene-RCD pair associations into a single, traceable context where every claim can be verified against its source. The compendium transforms distributed, heterogeneous knowledge, 62 original name strings across 20 resources, into a unified resource of 53 RCD-related entities, each with gene-level evidence, source provenance, and a seven-class mechanistic classification. The name RCDome reflects the architectural intent: a structure that provides comprehensive overhead coverage, unified enclosure, and a single vantage point from which to survey the entire RCD landscape.

For the researcher asking “Is gene X involved in ferroptosis, and what is the evidence?”, RCDome replaces hours of database cross-referencing with a single query that returns the answer, the provenance, and the evidence tier. The public codebase and the formal governance contract ensure that every step from database extraction to evidence verification is reproducible and auditable.

### What the atlas reveals about the biology of regulated cell death

The full-matrix classification, read as a whole, paints a picture that surprised us. The death-executing machinery is remarkably small: only 225 gene-RCD pairs (4.7% of the role-resolved set) are executors, the genes that directly pore, cleave, or permeabilize to kill the cell. Among the role-resolved pairs, 4,455 (93.3%) were regulators that promote or inhibit death without executing it. Within the compiled RCDome association universe and under the evidence-first adjudication, the mechanistically involved set is dominated by regulatory associations surrounding a comparatively small executor subset (Figure 4A).

The signal is sharply stratified by ontological level. The L1 modalities, the bona fide death forms, carry 38.9% of their pairs as mechanistically involved, whereas the L5 concepts (Autophagy, Programmed cell death) carry only 11.8%, despite being the largest association hubs. Most genes associated with autophagy are associated with the process, not with its lethal outcome; the evidence-first adjudication makes that distinction explicit at this scale (Figure 4B). This stratification is biologically informative, but it should not be read as an independent validation of the ontology: L1 was defined precisely to concentrate mechanistic forms, so a higher mechanistic fraction in L1 is in part expected by construction.

The direction of regulation is nearly balanced, 1,955 promoters versus 2,083 protectors (across all role-resolved associations), consistent with death as a decision that cells weigh in both directions, not a one-way cascade (Figure 4C). And 43.2% of the 9,265 genes have at least one mechanistically-involved association, confirming that RCD is woven through a large fraction of the human genome. Taken together, these numbers change how the field should be read: the core of regulated cell death is small, and most of the compiled network is regulation of death, not execution of it (Figure 4D). Gene coverage is highly heterogeneous across the 53 entities, from Apoptosis (5,897 unique genes) and Autophagy (4,634 unique genes) to Pyronecrosis (1 gene) (Figure 5). Where a subtype carries more genes than its parent, Autosis (1,105 genes) against Autophagy-dependent cell death (110), the difference reflects the depth of database indexing, not ontological inheritance; parent–child gene inventories are not expected to be nested subsets.

**Figure 5.**
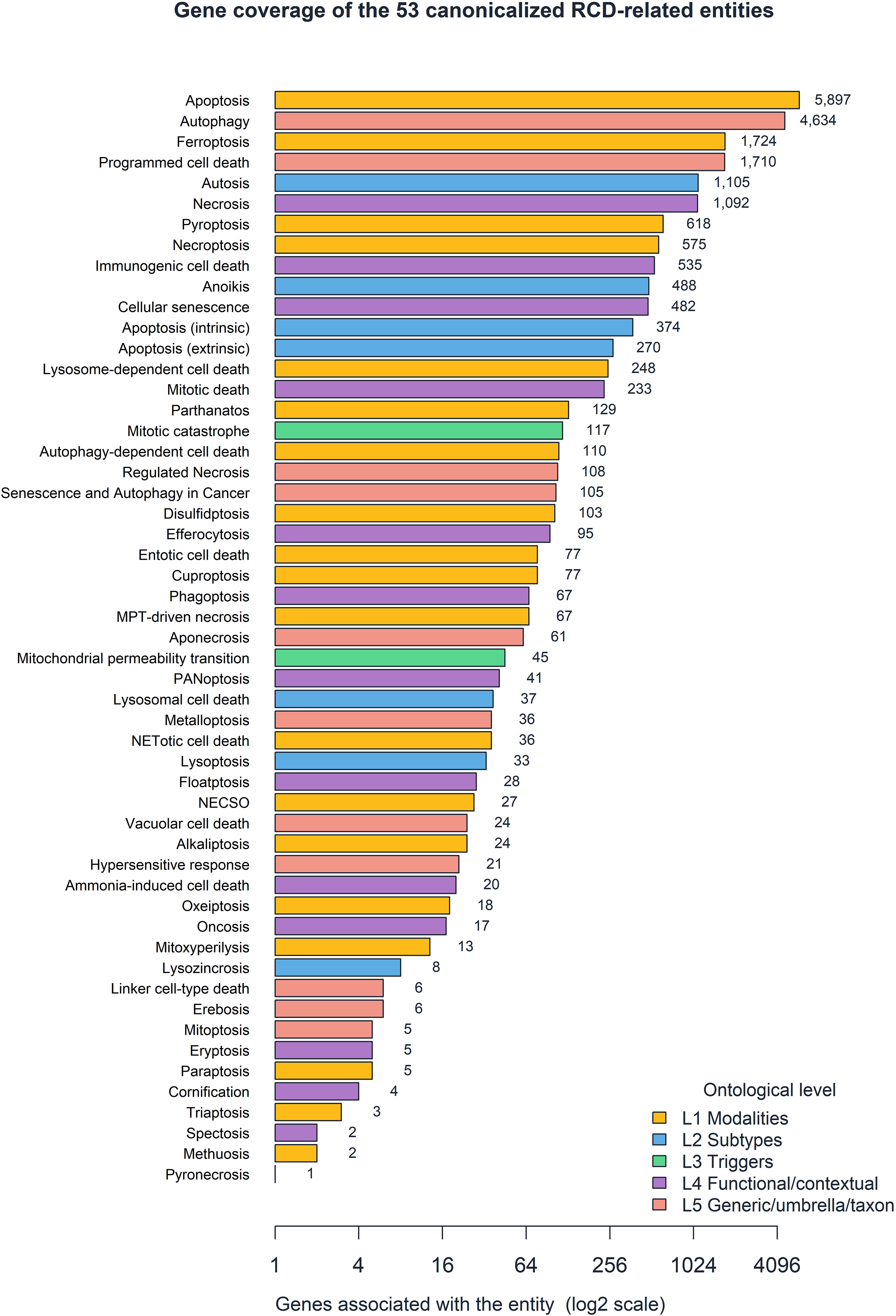
Gene coverage of the 53 canonicalized RCD-related entities. Each horizontal bar shows the number of distinct genes associated with one entity in the unified matrix (a gene linked to several entities is counted in each), colored by ontological level (L1 modalities, L2 subtypes, L3 triggers, L4 Functional/contextual categories and L5 Generic/umbrella/taxon concepts). The gene-count axis is log2-scaled so that the many entities with fewer than ten genes remain visible, and the exact gene count is printed at the end of each bar. Coverage is highly heterogeneous, from Apoptosis (5,897 unique genes) and Autophagy (4,634 unique genes) to Pyronecrosis (1 gene).

### The value and limits of the automated classifications

The three independent classifications, DeepSeek (the chain-of-thought full-matrix classification), Elicit (the original AI-assisted extraction), and Consensus (the third independent evidence channel), are preserved separately and can be compared. DeepSeek and Consensus each provide full 21,568-pair coverage; Elicit covers the subset where its literature-mining evidence is available. The evidence-first adjudication resolved all 21,568 pairs: 5,138 (23.8%) mechanistically involved, 16,236 (75.3%) not, and 194 (0.9%) out of context, with 225 executors and 4,455 regulators. The RCDome interface exposes these side by side through a source selector, and a researcher can extract the filtered table, the complete 100-column matrix, the per-entity comparison, or the 225-pair death core for their own work.

This utility should be used with eyes open. Scaling a majority-vote classifier to 21,568 pairs necessarily leaves a residual error rate. We repaired 84 self-inconsistent rows after the fact, and we deliberately retained 231 rows as uncertain rather than forcing them into a category, because a wrong-but-confident label is worse than an honest unresolved. The benchmark record is strong, and it was built to be method-independent: the four landmark papers, the 17-association accuracy ground truth, and the 26-pair benchmark set, were all fixed reference sets defined independently of the classifiers themselves, the classifications were never graded against their own outputs. On these fixed development reference sets (not independent held-out validation), the DeepSeek chain-of-thought classifier and the Consensus neutral-query rule both reached 100% combined accuracy with zero false positives. But those reference sets are well-established associations; they cannot guarantee that every one of 21,568 automated classifications is correct. Some percentage of the classifications will be wrong, and we want to find them.

We therefore envision RCDome not as a finished archive but as an active application that improves through use. We invite correction through two channels: the contact option built into the application itself, and the issue tracker of the GitHub repository. Every report will be traceable to its source and reconciled in a subsequent release.

The 53 RCD-related entities are explicitly heterogeneous, and we therefore refer to them as RCD-related entities rather than 53 established human modalities. Several are comparative or cross-species programs (hypersensitive response, linker cell-type death, eryptosis, cornification), others are morphological outcomes or umbrellas (necrosis, oncosis, regulated necrosis, metalloptosis), and others are emerging or legacy forms. Recognizing this heterogeneity strengthens rather than weakens the atlas: it reports the field as it is.

In this light, nomenclature in this field is dynamic, not binary. PANoptosis, eryptosis, and several emerging forms are presented with their evidence maturity (established, supported, emerging, legacy, or provisional) rather than as settled verdicts, so that the atlas tracks consensus as it evolves.

The automated channels (DeepSeek, Elicit, Consensus) preserve association and evidence for every pair, but the final ontological adjudication, what counts as a modality, subtype, trigger, or umbrella, is curator-governed, consistent with the governance rules of this project.

Taken together, RCDome answers the question it was built to answer (where each gene-RCD association comes from and what supports it) while making explicit how much of the field remains unverified. It is best read not as a finished archive but as infrastructure for a field still forming: a traceable substrate on which the community can correct, extend, and re-adjudicate as the biology, the evidence, and the nomenclature mature.

### Limitations

Four limitations are explicitly acknowledged. First, twenty-eight of the 53 RCD-related entities were not covered by the previous multi-optosis operational-definition set and therefore required RCDome-specific definition work; in the current decoder, all 53 entities have an operational definition anchored to a reference PMID. Second, the automated classifications rely on LLM calls whose residual error rate cannot be guaranteed across all 21,568 pairs; we repaired 84 self-inconsistent rows and retained 231 as explicitly uncertain, but some fraction of the remaining automated verdicts will be wrong and is open to community correction. Third, 286 DATABASE_ONLY pairs (1.3%) lack any verifiable paper trail in the indexed literature, and 86.3% of validated rows are not mechanistically confirmed, limits of the published record itself rather than of the platform. Fourth, the controlled vocabulary is a governed snapshot of the published record at the matrix freeze date (Dataset S1 freeze date: 2026-09-01: forms named after the freeze (e.g., ruptosis, 2026) or described only in non-human systems are documented as explicit exclusions in the Materials and Methods rather than silently omitted from the 53-entity vocabulary.

## Supporting information

Supplementary Tables S1 to S15 workbook

Dataset S1 matrix

## STATEMENTS

## Ethics approval and consent to participate

Ethics approval is not applicable to this study.

## Data and Code Availability Statement

The RCDome compendium matrix (Dataset S1; 22,473 rows × 100 columns; freeze date: 2026-11-01), extraction pipeline, and the RCDome web application source code are available at https://github.com/BioCancerInformatics/RCDome. The 20 association databases and resources are publicly accessible at the URLs listed in Supplementary Table S2. All extraction scripts, integration code, and the RCDome Shiny web application source code are available at https://github.com/BioCancerInformatics/RCDome. The complete extraction pipeline is orchestrated by master_extract.py, which executes all 23 channel_*.py extraction scripts in sequence (23 of the 56 scripts in the full pipeline catalog, Supplementary Table S10), producing timestamped, traceability-compliant CSV outputs. The repository also includes automated smoke tests with continuous integration that verify the mechanistic classification mapping, the chain-of-thought governance rules, and the benchmark scoring. RCDome is freely available at https://rcdome.org/.

## Author contributions

E.R.S. conceptualized the study; developed, validated, and implemented the R-based and Python computational pipeline; performed integrative analyses; contributed to manuscript revision; and developed the LLM-based RCD entity module of the Shiny application; and generated the AI-assisted figures.

V.dS.L. conceptualized the study, curated data, performed cross-referencing, and edited the manuscript.

H.A.C.N. curated data and edited the manuscript.

E.M.A. conceptualized and supervised the study; contributed to methodological design; developed, validated, and audited the computational pipeline; curated data; performed cross-referencing; drafted the manuscript, generated the conventional figures, and approved the final version.

## Acknowledgements

The authors acknowledge the public databases and resources used in this study, which provided the data for the RCDome gene-RCD compilation. If any gene or association was interpreted incorrectly, the authors request contact so the record can be assessed and amended in the live RCDome application. Generative artificial intelligence tools were used in a supervised, author-controlled capacity as detailed in the Generative AI Statement.

## Consent for publication

Not applicable.

## Generative AI Statement

Generative artificial intelligence tools were used in a strictly supervised, author-controlled capacity throughout the study and during manuscript preparation. The authors directed, audited, and approved every AI-assisted step, and no AI system exercised independent scientific judgment.

DeepSeek (deepseek-chat, deepseek-reasoner, and deepseek-v4-pro) served as the core language model of the compilation and validation pipeline. It executed the two-call independent validation of gene-RCD associations, in which one call drew on internal knowledge and the other on PubMed abstracts, and only concordance between the calls produced a confirmed classification. It executed the Europe PMC full-text deep retrieval-augmented evidence extraction, the full-text recovery of PMIDs, and the chain-of-thought classification of the complete matrix, which classified all 21,568 unique pairs with a mechanistic verdict, functional role, and direction. It also drafted the operational definitions of the 21 RCDome-drafted entities, which were substantiated by a three-stream validation composed of a DeepSeek claim-level check, a Consensus cross-reference, and a PubMed E-utilities verification, and then subjected to assistant-draft/human-approval review with every decision logged. An anti-hallucination framework verified every extracted sentence by exact string matching against the source document, and sentences not found verbatim were rejected and logged. During manuscript preparation, DeepSeek executed the predefined consistency checks and the nine-component compliance audit specified by the authors.

Consensus (the academic search engine) was used for the AI-assisted literature screening of Channel C, for the independent cross-validation of each de novo operational definition against coinage, mechanism, and PubMed-consensus evidence streams, and as the third independent evidence channel, which classified each gene-RCD pair from a neutral query composed of the gene symbol followed only by the term cell death.

Elicit was used for the Phase 3 systematic review and extraction of gene-RCD associations across 50 RCD forms, with post-export governance comprising allow-list enforcement, role normalization, and publication-traceability checks.

The generative-AI extraction of five multi-optosis operational definitions from the prior study, performed with *GPT-4o*, is retained in RCDome and was human-validated.

The RCD entity descriptions of the RCD Entities module were developed with the assistance of Consensus and DeepSeek, and were validated through a multi-phase anti-hallucination audit that extracted 2,812 claims across the 53 descriptions, verified each claim against its cited source, and classified each as supported (1,994), not supported (284), or uncertain (534). A subsequent re-anchoring pass re-anchored 669 of the not-supported and uncertain claims to peer-reviewed sources and removed 140 claims without support (Supplementary Note S8). The cartoon representations and logos of the RCD entities were generated from these descriptions under the figure-generation (Supplementary Prompt S3), figure-legend generation (Supplementary Prompt S5), logo creation (Supplementary Prompt S6), and scientific-accuracy audit (Supplementary Prompt S4) frameworks.

Every AI-extracted association was flagged as *`curator_verified`* false until manual expert review. The ontology of the 53 entities was adjudicated individually by the primary curator and independently reviewed entity-by-entity by two additional curators. The traceability contract, the hard-gate admission framework, and the anti-hallucination controls were enforced as executable code.

Verification does not rest on one model judging another; it rests on deterministic string matching against the source text and on human adjudication of every entity, neither of which is a language model.

The scientific rationale, study design, analytical strategy, auditing framework, verification criteria, manuscript organization, interpretation of results, scientific conclusions, and all editorial decisions were conceived, directed, and approved exclusively by the authors. All AI outputs were critically reviewed, manually verified, and, where appropriate, modified by the authors before incorporation. The authors assume full responsibility for the accuracy, integrity, and scientific validity of all text, analyses, code, and conclusions presented in this work.

## Publisher’s note

All claims expressed in this article are solely those of the authors and do not necessarily represent those of their affiliated organizations, or those of the publisher, the editors and the reviewers. Any product that may be evaluated in this article, or claim that may be made by its manufacturer, is not guaranteed or endorsed by the publisher.

## Significance Statement

Artificial intelligence is increasingly used to curate scientific knowledge, yet most such systems remain opaque and unverifiable: they produce a claim without showing whether it is grounded in its source. This work addresses that problem directly. RCDome demonstrates a verifiable, human-governed artificial intelligence pipeline for scientific curation: a two-call large language model architecture, an exact-string anti-hallucination framework, three independent evidence channels, and a governance contract enforced as executable code. Applied to regulated cell death, a field in which dozens of databases record gene associations under conflicting names and most have never been checked against the primary literature, this pipeline integrates 20 public databases and resources into a single traceable atlas of 53 curator-adjudicated entities, systematically evaluates every gene-RCD association, and states plainly what is supported and what is not. The atlas is delivered through the RCDome Shiny web application (https://rcdome.org/), a one-stop compilation that lets any researcher, in a single place, query which genes execute or regulate a given RCD form and inspect the evidence behind each answer. The result is a general demonstration that machine-assisted scientific knowledge can be made auditable, reproducible, and honest about its own uncertainty, a contribution that bears directly on the central challenge of applied artificial intelligence: producing machine-generated knowledge that researchers can trust. By making the evidence behind each gene-RCD association traceable and auditable, RCDome provides the curated foundation on which therapeutic targeting of cell death pathways can be built, for example in cancer and degenerative disease. As a free, one-stop, traceable atlas, RCDome lowers the access barrier to curated cell-death knowledge, giving researchers without specialized computational infrastructure access to traceable and audited gene-RCD evidence.

## Supplementary material

The Supplementary Material for this article can be found online at: https://github.com/BioCancerInformatics/RCDome.

