## Supplementary information for "RCDome: A Multi-Database Human Gene Atlas of Regulated Cell Death"

### Supplementary Notes

### Supplementary Note S1

### Canonical-Terminology Pre-Audit and 53-Entity Expert Nomenclature and Ontological Adjudication

#### Purpose

This note documents the governance and quality-control framework applied to RCDome's terminology and ontology: a terminology pre-audit performed before entity-level adjudication, a 53-entity expert nomenclature and ontological adjudication, and a project-wide propagation verification that closes each audit.

#### 1. The canonical-terminology pre-audit

Before any biological adjudication, RCDome terminology was subjected to a dedicated **pre-audit** of the word *"canonical"* and its variants across the manuscript, supplementary information, tables, decoder, matrix metadata, and the web application.

The risk being managed was semantic conflation: a reader could interpret *"canonical"* (vocabulary normalization) as *"canonical"* (biological establishment), and thereby read the 53-entity vocabulary as a list of 53 established cell-death modalities.

The pre-audit therefore required that **every** occurrence of *"canonical"* be adjudicated semantically into one of four categories:

• **vocabulary normalization**: the term was renamed *"canonicalized"* or *"normalized"*;

• **neutral entity designation**: the term was replaced by *"RCD-related entity"*;

• **biological canonicality***: "canonical"* was retained only where the claim of biological establishment was scientifically justified; or

• **bona fide modality status**: the term *"bona fide RCD modality"* was used for L1 entities admitted through the hard gates.

A global find-and-replace of *"canonical"* → *"canonicalized"* was deliberately prohibited, because it would have altered legitimate biological uses (e.g., *"canonical executors"*, *"canonical apoptosis"*, *"canonical inflammasome"*) and corrupted scientific meaning.

#### 2. The 53-entity expert nomenclature and ontological adjudication

Each of the 53 canonicalized RCD-related entities was then assessed against a five-level ontology and a set of hard gates.

**Ontology levels.** L1 (bona fide RCD modality: distinct causal machinery and functional necessity); L2 (subtype: mechanistic subdivision of a parent modality); L3 (trigger: an initiating event that is not itself a complete death mechanism); L4 (functional/contextual: context-dependent programs, morphological outcomes, cell-type-specific and differentiation-associated phenomena); L5 (generic/umbrella/taxon).

Hard biological gates (BG0–BG6). BG0 whole-cell death; BG1 identifiable causal machinery (obligatory for L1); BG2 functional necessity and distinguishability (obligatory for L1); BG3 supportive phenotype/trigger; BG4 published naming/provenance; BG5 independent replication (a maturity input); BG6 species scope. Naming alone (BG4) is not evidence of mechanistic independence.

**Association vs modality.** Two dimensions are kept separate: (i) whether a gene is associated with an RCD-related entity in the evidence base, and (ii) whether the entity itself qualifies as a bona fide RCD modality. Evidence of the former does not establish the latter. Accordingly, the associated-gene list is not described as a *"death signature"* or *"death machinery"* unless the evidence classification supports that statement; the preferred formulation is *"genes associated with [entity] in RCDome"*.

**Uncertainty and provenance.** Entities that fail L1 admission are not deleted. They are retained at an appropriate level (contextual category, morphology, subtype, trigger, umbrella, taxon-specific concept, legacy term, or provisional/emerging entity) with explicit provenance. Database inclusion is never treated as biological endorsement.

#### 3. The propagation governance gate

The audit established a rule that has since governed all subsequent corrections: a change is not considered implemented when the authoritative source has been corrected alone.

Every accepted adjudication must be propagated:

• **horizontally**: to every artifact that reproduces the affected information (manuscript, supplementary information, supplementary tables, decoder, unified matrix, database fields, figures and legends, application components, landing pages, explanatory text, and downloadable outputs); and

• **vertically**: through every dependent representation derived from it (counts, group distributions, summaries, labels, colors, tooltips, and narrative prose).

For example, reclassifying one entity from L4 to L5 is not complete until the entity's classification, the level counts (e.g., the L4/L5 totals), the figures, the legends, the landing-page summaries, the supplementary tables, and the application display have all been updated and independently verified.

Any residual disagreement between the authoritative adjudicated source and a downstream artifact is treated as a propagation failure and recorded (original state, adjudicated state, affected dependency, stale representation, corrective action, and post-correction verification), never silently repaired.

#### 4. Traceability and reproducibility

Every change carries an audit trail: what changed, why it changed, the prior state, the evidence, and where the consequences were applied. Versions are archived rather than overwritten, preserving the history of the project. This propagation gate (adjudicated → propagated → independently verified) is the final governance check before an audit is declared complete, and it is re-run after any subsequent modification to guarantee internal concordance across the entire RCDome ecosystem.

- The evidence-quality audit.
- The contradiction audit and the repair logs.
- The fetched abstracts.
- The vote outputs (one file per evidence source).

### Supplementary Note S2

### Label-Error Benchmark: Disagreement Arbitration, Evidence-First Adjudication, and Semantic-Contradiction Repair

#### 1. Purpose

this note explains how RCDome measures the error rate of its classification labels where the independent classifiers disagree, and how internal semantic contradictions in the matrix were found and repaired. The benchmark arbitrates discordance with an independent vote and a human verdict, preserving the original classifications, and the audit-and-repair removes self-contradictions by conservative downgrade.

#### 2. The classification layers and the disagreement set

The study carries **three classification outputs** and **one adjudication layer**:

| Layer | Role | Labels (vocabulary) |
| --- | --- | --- |
| DeepSeek | classification | role (EXECUTOR/REGULATOR/MARKER/SUBSTRATE/INTERACTOR/UNCERTAIN) · direction (PROMOTES/PROTECTS/UNCLEAR) · mechanistic (YES/NO) |
| Consensus | classification | role (EXECUTOR/REGULATOR/MARKER/UNCERTAIN) · direction (PROMOTES/PROTECTS/UNCLEAR) · mechanistic (YES/NO) |
| Elicit | classification | role_direction (PROMOTES/PROTECTS/UNCLEAR/ASSOCIATED/BIDIRECTIONAL) · evidence_type (GENETIC_PERTURBATION/PHARMACOLOGICAL/EXPRESSION_CORRELATION/CITATION_ONLY/CURATED) — no role, no mechanistic |
| GPT + human | adjudication | verdict (DEEPSEEK/CONSENSUS/NEITHER/UNCLEAR/OUT_OF_CONTEXT) — arbitrates, does not classify |

A pair is a disagreement when two classifiers that share a label are both determined and their values differ. Across the shared labels this yields 7,905 label-disagreements (role 320, direction 935, mechanistic 6,650) on 7,831 unique pairs. The disagreements concentrate on three boundaries: role is the executor–regulator boundary, direction the promote–protect boundary, and mechanistic the involved–not-involved boundary.

A disagreement between classifiers does not necessarily indicate an inconsistency or an error: the classifiers extract from different evidence sources, so the same pair may be judged on different papers. The two causes are distinguished below.

##### What is comparable, per label

| Label | Compared between | Shared vocabulary | Arbitrable? |
| --- | --- | --- | --- |
| role | DeepSeek vs Consensus | EXECUTOR / REGULATOR / MARKER / UNCERTAIN | yes (Elicit excluded — no role) |
| direction | DeepSeek vs Consensus vs Elicit | PROMOTES / PROTECTS | yes (all three emit direction) |
| mechanistic | DeepSeek vs Consensus | YES / NO | yes (Elicit excluded — no mechanistic) |

##### Why some labels are not arbitrable

- Elicit has no role and no mechanistic, so it cannot enter those

comparisons; they remain DeepSeek-vs-Consensus.

- DeepSeek's SUBSTRATE and INTERACTOR have no Consensus counterpart:

vocabulary differences, not disagreements.

- Elicit's ASSOCIATED and BIDIRECTIONAL have no DeepSeek/Consensus

counterpart: likewise vocabulary differences.

##### The two distinct causes of disagreement

1. 1. Source-coverage disagreement: the instruments consult different evidence

(LLM knowledge plus PubMed abstracts for DeepSeek; the Consensus API corpus; the Elicit systematic review). The same pair may be judged on different papers.

2. 2. Interpretation disagreement: the same evidence is read, but different

labels are assigned (one reads a pore-forming gene as EXECUTOR, another as REGULATOR).

The benchmark does not eliminate either. It reports both, and arbitrates only the interpretation disagreements; source-coverage differences are documented as design.

##### The evidence

Every disagreement is arbitrated against real source evidence. Where a stored verbatim sentence exists, it is used directly; where it does not, the source abstract is fetched from the DeepSeek PMIDs (Europe PMC), falling back to the Consensus DOI. Of the pairs that required a fetch, 98.5% resolved to a readable abstract; the remainder were unadjudicable.

#### 3. The third-opinion vote

Each disagreement is submitted to `deepseek-reasoner` with the disputed label, the two classifier values, and the evidence. The model votes one of five verdicts: DEEPSEEK, CONSENSUS, NEITHER (both wrong, with the correct value stated), UNCLEAR (evidence cannot decide), or OUT_OF_CONTEXT (the Consensus evidence concerns a different gene or context). The vote guides the human curator, who remains the gold standard.

The model returns a single fused line, a verdict followed by its reason. A semantic consistency check then reads the reason against the verdict and, where they contradict, flags the row (FLAGGED) for the human curator; it never auto-overrides a vote.

The vote is the model's guide only. The human curator's final verdicts are collected separately and are complete (all curators reviewed); the numbers below are the third-opinion vote, not the final arbitration.

The adjudicated truth produced by this process is a synthesis, not a ground truth: it is the best available resolution of discordance, produced transparently, and it never replaces the original classifications.

#### 4. Results

The results below report the model's third-opinion vote. The final human arbitration is complete (all curators returned their verdicts).

**Cross-classifier agreement.** Among pairs where both classifiers were determined, DeepSeek and Consensus agree on role in 87.6% (1,011/1,154), on direction in 85.4% (986/1,155), but on the binary mechanistic verdict in only 65.0% (2,796/4,304).

**Third-opinion verdicts (n = 7,892).**

| Verdict | Count | % |
| --- | --- | --- |
| DEEPSEEK | 3,430 | 43.5% |
| CONSENSUS | 2,533 | 32.1% |
| UNCLEAR | 1,314 | 16.7% |
| OUT_OF_CONTEXT | 590 | 7.5% |
| NEITHER | 25 | 0.3% |

The model committed to a side in 75.6% of disagreements and abstained honestly in 24.4% (UNCLEAR 16.7%, OUT_OF_CONTEXT 7.5%, NEITHER 0.3%).

**Consistency.** The semantic check flagged 285 verdict-vs-reason contradictions for the human curator (never auto-corrected) and confirmed 1,174 rows as consistent.

Interpretation. The two classifiers converge on who and how (role/direction, 86–88%) but diverge on the loosest claim: is the gene involved at all (65%). A quarter of the disputes cannot be cleanly arbitrated from the available evidence, and the atlas reports that honestly rather than forcing a call.

#### 5. Semantic-contradiction audit and repair

An independent audit of the matrix found **337 internal contradictions** (237 DeepSeek, 100 Consensus): verdicts violating their own governance rules (role vs direction vs mechanistic). All were repaired by conservative downgrade-only rules and logged:

- 181 REGULATOR-with-bad-direction → UNCERTAIN/UNCLEAR (Rule 2)
- 237 mechanistic YES-with-UNCERTAIN-role → NO (Rule 5, including the cascade)
- 80 mechanistic NO-with-confident-role → UNCERTAIN (Rule 4; already present)
- Consensus: 75 downgraded; 19 umbrella cases kept and logged

Post-repair re-audit: 0 contradictions remaining in both the matrix and the Consensus results. The repaired matrix (100 columns, including the 11 Consensus variables).

---

#### 6. Evidence-quality finding

A deterministic audit of the disagreements found:

- 42.1% have a bullet-fragment verbatim (a gene-list item, not a sentence),

but 100% still carry the full Consensus title/takeaway: so they remain arbitrable.

- 7.9% have the gene absent from both sides.
- 20.4% have the RCD form absent from both sides.

The benchmark therefore runs on the full disagreement set, and the model votes UNCLEAR/OUT_OF_CONTEXT where the evidence does not support a clear call.

---

#### 7. Key decisions

3. 1. Verbatim-grounded scope: stored evidence arbitrated directly; missing

evidence fetched, not skipped.

4. 2. Label-level granularity: one row per (pair, label).
5. 3. Human is the gold standard: the model vote is a guide only.
6. 4. Resume-safe and logged: every run checkpoints; every flag and repair

is flagged with its rule and date.

7. 5. Never replace, always add: the original classifications are preserved

byte-identical; the benchmark adds new variables (vote, reason, consistency, human verdict, adjudicated truth) alongside them.

---

#### 8. Traceability files

- The vote outputs (one file per evidence source).
- The fetched abstracts.
- The contradiction audit and the repair logs.
- The evidence-quality audit.

**9. Evidence-First Adjudication**

The final evidence-first adjudication (the final_* columns reported in the manuscript) resolves every pair across three pools under the evidence hierarchy above.

The three pools (21,568 unique pairs): Agreement 13,737 (AGREED_LABEL_ABSTRACT); Disagreement 6,883 (UNIFORM_PIPELINE); Delta 948 adjudicated (DELTA_ADJUDICATED), 0 remaining (PENDING_DELTA).

The central rule — a vote is not a vote. Adjudication is not an agreement poll. The final verdict is decided by the evidence (verbatim > abstract), never by who agrees with whom. A verdict, including a unanimous one, is overturned only by evidence of equal or greater strength (full-text verbatim).

The seven rules. (1) Verbatim-grounded: deciding_evidence records the exact fragment that decided each row, labelled [verbatim] or [abstract]. (2) Support rubric: each reason is scored APOIADA / NAO_APOIADA / PARCIAL against that evidence. (3) Asymmetric ceiling: verbatim can carry the verdict; abstract-only support is at most PARCIAL. (4) OUT_OF_CONTEXT must be demonstrated (a gene/form/process mismatch), itself checked against the evidence. (5) Negative defaults: mechanistic → NO (no causal claim); direction → UNCLEAR (no directional claim). (6) Abstract never cancels: a fetched abstract is recorded CONFIRMED / INSUFFICIENT / DIFFERENT_VIEW, never a cancellation. (7) Additive, never overwrite: the original cot_* / consensus_* columns remain byte-identical; the adjudication only adds columns.

GPT-4o arbitration and the two curators. Each disagreement is submitted to GPT-4o (independent of DeepSeek and Consensus), which votes DEEPSEEK / CONSENSUS / NEITHER / UNCLEAR / OUT_OF_CONTEXT. Two human curators (Curator 1, Curator 2) record verdicts independently. The triple vote is classified CORROBORATED / HUMANS_AGREE / GPT_PLUS_ONE / TOTAL_DISAGREEMENT; the final label follows the majority, and TOTAL_DISAGREEMENT remains pending.

Final counts (unique pairs). mechanistic: YES 5,138 (23.8%) · NO 16,236 (75.3%) · OUT_OF_CONTEXT 194 · pending 0. role: EXECUTOR 225 · REGULATOR 4,455 · MARKER 95 · UNCERTAIN 16,793 · pending 0. direction: PROMOTES 2,102 · PROTECTS 2,083 · UNCLEAR 17,382 · pending 0.

Agreement pool. The agreement evidence is a DeepSeek label plus a Consensus abstract (not full-text). The agreement stands; the fetched-abstract re-check is a non-cancelling view, recorded CONFIRMED 481 / INSUFFICIENT 13,019 / DIFFERENT_VIEW 237.

Delta round. 948 pairs were adjudicated (DELTA_ADJUDICATED) through the same evidence-first pipeline. no pairs remain PENDING_DELTA, because their original abstract fetch was unadjudicable (no readable abstract): PEAR1/Efferocytosis, MAGED1/Apoptosis (extrinsic), MIR24-1/Programmed cell death. Their full-text verbatim has since been retrieved, and their verdicts were subsequently adjudicated in the curator round.

### Supplementary Note S3

### RCDome: Model-Selection Benchmark (CoT + Self-Provided Loop)

This note documents the design and results of the model-selection benchmark, which compared three functional-role classifiers on 26 well-established gene-RCD pairs and selected a self-provided governance loop as the role-classification engine.

#### Design

Twenty-six well-established gene-RCD pairs with known functional roles, in three strata:

10 true EXECUTORS: *BAX*, *CASP3*, *CASP8*, *CASP9*, *CYCS* (Apoptosis); *MLKL*, *RIPK3* (Necroptosis); *GSDMD*, *GSDME*, *CASP1* (Pyroptosis).

8 true REGULATORS: *BCL2*, *BCL2L1*, *XIAP*, *TP53* (Apoptosis); *GPX4*, *ACSL4*, *AIFM2* (Ferroptosis); *BECN1* (Autophagy).

8 NEGATIVE controls: *GPX4*/Pyroptosis, *MLKL*/Ferroptosis, *CASP3*/Necroptosis, *BAX*/Necroptosis, *GSDMD*/Ferroptosis, *CASP1*/Apoptosis, *MLKL*/Apoptosis, *GSDMD*/Necroptosis.

The 26 pairs comprise 18 positive pairs (the 18 unique positive genes: 10 executors plus 8 regulators) plus 8 form-mismatch negatives that reuse 8 of those genes with an incorrect form.

Each pair was submitted to all three models in the same run with the identical regulatory-aware prompt (mechanistic, role, direction).

Two loop variants were benchmarked. The reference-based variant (v7) used a hand-curated 18-gene reference (one entry per unique positive gene: known form + mechanism) and achieved 100% combined accuracy with zero false positives, but that reference cannot scale: only about 5% of the compendium's pairs have a complete role, direction, and mechanistic classification to reference (1,102 rows; 573 genes; 875 pairs). The self-provided variant (v11) instead makes the model state its own mechanism, known forms, and role-specific form, then self-checks those statements; it achieved the same 100% combined accuracy and zero false positives with no external reference. Because it is fully independent and scalable, v11 was adopted for the full 22K-pair matrix.

#### The self-provided loop (adopted)

The winning arm is a tool-call loop on *deepseek-v4-pro* (thinking mode, *reasoning_effort*=max) with: three-fold self-consistency voting; six logic rules; and three self-provided fields the model must state in every call - its molecular mechanism, its known RCD form(s), and the single form its role belongs to (*role_form*). Three self-consistency rules then check those statements: an executor-keyword rule, a form-mismatch rule, and a role-form-specificity rule. No external reference is used; the gene-form pairs are drawn from the matrix, but the loop never consults the matrix's stored role or direction columns, so its classification is independent of that reference.

#### Results (same run, 26 pairs)

| Metric | deepseek-chat | deepseek-reasoner | self-provided loop (v11) |
| --- | --- | --- | --- |
| Combined accuracy | 21/26 (81%) | 19/26 (73%) | 26/26 (100%) |
| EXECUTOR sensitivity | 9/10 (90%) | 6/10 (60%) | 10/10 (100%) |
| EXECUTOR precision | 9/11 (82%) | 6/7 (86%) | 10/10 (100%) |
| REGULATOR sensitivity | 7/8 (88%) | 7/8 (88%) | 8/8 (100%) |
| REGULATOR precision | 7/10 (70%) | 7/13 (54%) | 8/8 (100%) |
| Coherent rejection | 5/8 (62%) | 6/8 (75%) | 8/8 (100%) |
| False positives | 3/8 (38%) | 2/8 (25%) | 0/8 (0%) |
| Role-mech coherence | 23/26 (88%) | 26/26 (100%) | 26/26 (100%) |
| Direction accuracy | 7/8 (88%) | 8/8 (100%) | 8/8 (100%) |

#### Conclusion

The self-provided governance loop achieved 100% combined accuracy, 100% sensitivity and precision on both roles, and zero false positives - exceeding both baselines on every metric, without any external reference. It was therefore adopted as the role-classification engine for the compendium.

#### Code

*cot_benchmark.py* (v1) -> v2 -> v3 (loop) -> v4 (self-consistency + rules) -> v5 (coherence) -> v6 (executor-keyword) -> v7 (form-mismatch, with reference) -> v8 (agnostic) -> v9 (self-provided mechanism/*known_forms*) -> v10 (*role_form*) -> v11 (broadened executor words). All scripts reside in the "extraction code snippets" directory.

#### Chain-of-thought full-run benchmark (*deepseek-v4-pro* + self-governance loop)

Before the full-matrix run, the same 26 ground-truth pairs (10 executors, 8 regulators, 8 negative controls) were re-benchmarked against the production classifier: *deepseek-v4-pro* with thinking enabled, the verify-role self-governance loop, and the role-form specificity gate. The gate rejects a mechanistic-YES verdict whenever the model's self-stated role-form does not match the queried RCD form, targeting the form-over-generalization false positives (GPX4/Pyroptosis and MLKL/Apoptosis) that persisted in earlier iterations.

Results: the production classifier reached 26/26 (100%) combined accuracy and 0/8 (0%) false positives, against 2 false positives for *deepseek-chat* and 2 for *deepseek-reasoner* on the same set.

### Supplementary Note S4

### RCDome: Consensus API Classification Benchmark:

This note documents the design and results of the Consensus API classification benchmark, which tuned the Consensus evidence-channel classifier against the same 26 ground-truth pairs used for the DeepSeek model-selection benchmark (Supplementary Note S3).

#### Design

Twenty-six well-established gene–RCD pairs (10 executors, 8 regulators, 8 negative controls) served as ground truth. Seven iterations (v1–v7) tested query formulations: v1–v4 tested reference-query formulations; v5 combined reference and agnostic queries; v6 mirrored the DeepSeek self-governance loop; v7 introduced the neutral-query rule (the gene symbol followed only by "cell death"), removing form-name priming.

#### Results

The neutral-query rule (v7) reached 26/26 (100%) combined accuracy and 0/8 (0%) false positives, against 1/8 false positives for v6 (mechanistic 25/26; role 17/26). The neutral-query rule was therefore adopted for the full Consensus extraction.

| Iteration | Combined accuracy | False positives (of 8 negatives) |
| --- | --- | --- |
| v6 | mechanistic 25/26 (role 17/26) | 1/8 |
| v7 (neutral query) | 26/26 (100%) | 0/8 (0%) |

### Supplementary Note S5

### Canonicalization Logic and Separation from Biological Modality Status

#### Purpose

This note defines the term *"canonicalized RCD-related entity"* as used throughout RCDome, and explains why vocabulary canonicalization must be kept rigorously distinct from biological recognition of an entity as a canonical or bona fide regulated cell-death (RCD) modality.

#### 1. What canonicalization means in RCDome

RCDome consolidates gene–RCD association records recovered from 20 heterogeneous sources (curated databases, pathway resources, and literature-mining channels, plus omic-annotation compendia used for annotation only), each using its own nomenclature. A single biological concept therefore arrives under multiple names (e.g., "Entotic-cell-death", "Entotic cell death", and "Entosis" for the same entity).

**Canonicalization** is the vocabulary operation that maps these heterogeneous source terms onto a single preferred RCDome identifier while the original source term is preserved in a parallel field (`*rcd_form*`, alongside the normalized identifier `*rcd_form_canonical*`). The result is a controlled vocabulary of 53 canonicalized RCD-related entities.

Canonicalization is therefore an informatics operation on nomenclature, not a biological verdict.

#### 2. What canonicalization does NOT mean

Canonicalization does not imply that an entity is:

• a canonical or established human cell-death modality;

• mechanistically independent;

• validated in human cells;

• accepted by the Nomenclature Committee on Cell Death (NCCD);

• supported by a causal execution machinery; or

• equivalent in evidence maturity to apoptosis, necroptosis, ferroptosis, or pyroptosis.

Membership in the controlled vocabulary records only that the entity's terminology has been normalized into RCDome. Biological and ontological status is adjudicated separately and independently.

#### 3. Terminology hierarchy

Four terms are used deliberately, with distinct meanings:

a. **Canonicalized RCD-related entity**: one of the 53 preferred vocabulary identifiers produced by normalization and consolidation of heterogeneous source terminology. This is a vocabulary-normalization statement.

b. **RCD-related entity**: the neutral umbrella term for any member of the 53-entity controlled vocabulary, used when the normalization history is not the point of the sentence.

c. **Bona fide RCD modality**: an entity admitted to level L1 through the RCDome BG0–BG6 hard-gate framework (identifiable causal machinery and functional necessity/distinguishability). In the current adjudicated ontology, 19 of the 53 entities satisfy this criterion.

d. **Canonical RCD modality**: reserved for contexts that genuinely require a claim of biological establishment or authoritative mechanistic recognition, and only where that claim is supported by the evidence. This term is never used merely because an entity has been canonicalized in RCDome.

#### 4. The 53 ↔ 19 relationship

The 53 canonicalized RCD-related entities span five ontological levels:

• 19 bona fide RCD modalities (L1);

The relationship between the 19 L1 modalities and the NCCD's 20 canonical definitions is assessed in Methods ("Relationship to the NCCD canonical ontology").

• 7 subtypes (L2);

• 2 triggers (L3);

• 13 functional/contextual categories (L4);

• 12 generic/umbrella/taxon concepts (L5).

Thus, the headline count of 53 is a count of **normalized vocabulary entries**, not of established modalities. Only the 19 L1 entities are described as bona fide RCD modalities.

#### 5. Preservation of provenance

Canonicalization never overwrites the original source terminology. Every association row retains its source form (`*rcd_form*`) in parallel with the normalized identifier (`*rcd_form_canonical*`), so the provenance of each name (and therefore of each association) remains traceable to its source database or literature origin.

#### 6. Avoiding semantic leakage

Because *"canonical"* in informatics conventionally denotes a preferred normalized identifier, whereas in biology it can imply authoritative establishment, RCDome uses "canonicalized" for the vocabulary operation and reserves "canonical" and "bona fide" for biological/ontological status. This lexical distinction is maintained consistently across the manuscript, supplementary material, figures, tables, and the web application so that a reader cannot conflate inclusion in the controlled vocabulary with biological endorsement.

### Supplementary Note S6

### RCDome: Digital Molecular RCD Board Benchmark

This note documents the design and results of the Digital Molecular RCD Board benchmark, which evaluated a prose-synthesis module that answers gene–RCD queries strictly from the RCDome database and the RCDome decoder, with every claim traceable to a matrix row, a decoder field, or a recorded PMID.

#### Board governance

The board emits a single continuous prose synthesis with no headers, no bullet lists, and no markdown bold. Four provenance categories govern how each sentence is written without being printed as labels: facts with a PMID ("the RCDome database records … (PMID …)"), model annotation ("within the RCDome database …"), decoder context ("the RCDome decoder defines/states …"), and hypothesis ("Hypothesis:" or "may suggest" or "is consistent with"). Seven rules are enforced in the prompt and by a deterministic post-generation scrub, never by a second model: (1) only "RCDome database" and "RCDome decoder" are used, with the bare words "database", "decoder", and the internal term "matrix" banned (the matrix rule is biology-safe, ignoring extracellular matrix, mitochondrial matrix, and matrix metalloproteinase); (2) assertive causal verbs are replaced with hedging equivalents in all tenses; (3) genuine uncertainty phrasing is preserved; (4) the exact verbatim disclaimer is appended at the very end; (5) the synthesis body ends with exactly three numbered "Suggested Queries:" immediately before the disclaimer (auto-numbered, never invented); (6) only PMIDs present in the data may be cited, and the founding PMID is always cited when the entity's operational definition is described; and (7) a pair absent from the database is reported as "not recorded" without inventing a role or direction.

#### Q&A retrieval strategy

The Q&A submodule answers free-form questions across all 53 entities and all 9,265 genes. The complete gene–form matrix (22,473 rows, each carrying mechanistic class, role, direction, evidence tier, and PMID) exceeds the model context, so the Q&A uses a retrieval-augmented strategy that never deletes data. A static layer always injects the complete gene–entity index (all 22,473 associations, names only) and the full 53-entity decoder. A dynamic layer then detects, for each question, the entities mentioned (by normalized name or its stem, so "apoptotic" matches Apoptosis) and any gene symbols, and injects the full seven-layer records for exactly those rows. Genes named explicitly in the question are always placed first so the budget can never displace them. The dynamic layer is capped so the full prompt stays within the model context (about 58K tokens), with a reply token limit applied. No entity or gene is ever excluded from the index, and every classification (MECHANISTIC, LITERATURE_GAP, CONTEXT_GAP, CO_OCCURRENCE, DATABASE_ONLY, INSUFFICIENT_DATA) remains retrievable, so a gene recorded as executor in one form and regulator in another is shown with both records.

#### Benchmark design

The benchmark (*run_board_benchmark.py* and *audit_canonical.py*) evaluates the board on nine tests, each with its own statistically determined sample size, 95% Wilson confidence interval, and significance test. The deterministic scrub is applied to the raw model output before the audit, so the audit measures the final, scrubbed text; scrub corrections are logged separately. The audit is a fixed 28-point checklist per synthesis report: 10 governance checks (causal verbs, hedging, disclaimer, three queries), 14 policy checks (terminology and relationship terms), and 4 accuracy checks (cited PMIDs exist, founding PMID cited, role matches, direction matches). Census reports use a 24-point lookup audit without the query requirement, and negative controls use a 29-point audit with inverted role and direction checks. Sample sizes follow the standard formulas: n = z²·p·(1−p)/E² for a proportion at 95% confidence (z = 1.96); n = (zα+zβ)²·2·p̄·(1−p̄)/Δ² to detect a difference at 80% power (zβ = 0.84); and the Wilson 95% interval (p + z²/2n ± z·√(p(1−p)/n + z²/4n²))/(1 + z²/n).

#### Results

The benchmark ran on 2026-09-09 with seed 42, temperature 0.7, prompt v2-smoke-aligned, and the *deepseek-chat* model. Golden accuracy was 26/26 (100%; Wilson 95% CI 87.1–100%; binomial against 0.95, p = 0.639). Census accuracy was 53/53 (100%; CI 93.2–100%; binomial p = 0.114). Fact-stability on *BAX*-Apoptosis was 73/73 (100% per fact for gene, entity, and founding PMID; CI 95.0–100%). Governance showed 1 violation across 2,142 checks (0.047%; CI 0.008–0.264%), broken down as causality 0, hedge 0, disclaimer 0, queries 1, terminology 0, and accuracy 0. Anti-hallucination caught 4 of 4 traps (100%; CI 51.0–100%). Articulation conformed in 78 of 79 reports (98.7%; CI 93.2–99.8%). Model comparison favored *deepseek-chat* (26/26, 100%) over *deepseek-reasoner* (22/26, 84.6%). Chat governance passed 10/10 (100% across five question types). The single governance violation was *MLKL*-Necroptosis failing the exactly-three-queries check; on direct regeneration that same report passed 30/30, confirming this is temperature-driven prose variability rather than a systematic defect. The deterministic scrub applied 163 corrections across the run (55 bare database or decoder terms, causal-verb hedging, markdown-bold removal, and disclaimer and query normalization).

#### Q&A benchmark description and validation

The Q&A submodule was benchmarked with the same statistical machinery as the board benchmark (point estimate, Wilson 95% CI, binomial test against a 0.95 null), using *deepseek-chat*, temperature 0.3, a 1500-token reply cap, and the full retrieval-augmented prompt. Two tests were run. T10 positive accuracy (n = 100; 25 golden executor or regulator gene–form pairs, each queried on role, direction, evidence tier, and supporting PMID) audited five checks per answer: the correct fact is cited, the RCDome source is named, the interpretation is hedged, the disclaimer is appended, and no PMID absent from the data is cited; the result was 100/100 = 100% (Wilson 95% CI 96.3–100%, binomial p = 0.010). T11 negative controls (n = 100; 8 not-recorded pairs queried on the same four types, plus 68 out-of-scope or fabrication queries) were all reported as absent or refused without a hallucinated PMID; the result was 100/100 = 100% (Wilson 95% CI 96.3–100%, binomial p = 0.010). The significance of the Q&A submodule is that it makes the board a full-coverage interface rather than a single-entity viewer: the static layer keeps the complete gene–entity index and the dynamic layer retrieves the full seven-layer records on demand, so the Q&A answers across all 53 entities and all 9,265 genes without funneling the data. Every classification (MECHANISTIC, LITERATURE_GAP, CONTEXT_GAP, CO_OCCURRENCE, DATABASE_ONLY, INSUFFICIENT_DATA) remains reachable, and a gene recorded as an executor in one form and a regulator in another is shown with both records (for example, *BAX* as a mechanistic executor in apoptosis with PMID 17235284, together with its co-occurrence annotations in necroptosis and pyroptosis and its insufficient-data annotations in autosis, erebosis, and oncosis). The 100% positive and negative results recorded zero hallucinated PMIDs across the benchmark, demonstrating that full coverage and zero observed hallucination were achieved together in the same module.

**Statistical design.** The Q&A tests use the same sample-size formulas as the board benchmark: proportion 95% CI n = z²·p·(1−p)/E² (z = 1.96); detect a defect Δ with 80% power n = (zα+zβ)²·2·p̄·(1−p̄)/Δ² (zβ = 0.84); Wilson 95% CI lower bound for 100% observed n/(n + z²). A claim of accuracy at least 95% with 95% confidence requires the Wilson lower bound to reach 95%, which occurs at n = 73 (73/76.84 = 0.950). For two-sided significance against the 0.95 null, n = 100 gives p = 0.010. In power terms (80%): n = 77 detects a 15% absolute defect (p = 0.80), n = 142 detects 10%, and n = 436 detects 5%. The chosen n = 100 therefore detects an absolute defect of about 12.5% at 80% power, holds the Wilson lower bound at 96.3%, and gives a one-sided binomial p = 0.006 and a two-sided p = 0.010. T10 breakdown (25 pairs × 4 question types, all 100 passed): role 25/25, direction 25/25, evidence tier 25/25, supporting PMID 25/25. T11 breakdown: not-recorded 32/32 (8 pairs × 4 types), out-of-scope and fabrication refusal 68/68. Audit volume: T10 = 100 answers × 5 checks = 500; T11 = 32 not-recorded checks plus 68 refusal checks × 2 = 168; 668 checks in total, all passed.

#### Code corrections

Six defects were found and fixed during validation: (1) the violation-type mapping had mislabeled the hedging and query checks and contained a dead branch, so hedge and query counts were reported incorrectly; (2) every failed check is now logged with its pair, check identifier, name, and exact detail rather than only tallies; (3) the scrub now removes markdown bold markers; (4) entities with the sentinel category CORE_PENDING are no longer falsely flagged, which corrected census accuracy from 51/53 to 53/53; (5) the query-count articulation check now applies only to synthesis and negative reports, not to census lookups; and (6) the matrix policy check now ignores extracellular matrix, mitochondrial matrix, and matrix metalloproteinase.

#### Post-review regression benchmarks

The Board and Q&A were benchmarked on random, seed-reproducible samples of n = 100 gene–entity pairs or queries, with the exact binomial 95% confidence interval (Clopper–Pearson).

**Reference discipline.** Because a synthesis may cite both gene–RCD association evidence and entity-level definitions, the two citation levels must never be interchanged: a gene–RCD statement must cite the gene-specific association PMID, and a definition statement must cite the entity founding PMID. Across 100 associations, the synthesis separated the two levels correctly in all 100 cases (100%; 95% CI 96.4–100%).

**Context fidelity**. The Board receives the structured payload (role, direction, mechanistic class, evidence tier, *required_for_death*) for the queried gene–entity pair, and the prose must not contradict those fields. Across 100 pairs, no contradiction was observed (100%; 95% CI 96.4–100%).

**English-only policy**. The Board and Q&A answer only in English; a query in any other language must receive the exact refusal sentence. Across 100 non-English queries (Portuguese, Spanish, French), all 100 were refused correctly (100%; 95% CI 96.4–100%).

**Minimal source and insufficient-data directness**. For a gene–entity pair with no recorded PMID, the synthesis must state that no PMID is recorded and name the source database; for a pair with an INSUFFICIENT_DATA mechanistic classification, it must say so directly rather than expand empty fields. Across 100 cases of each, the correct behavior was observed in all 100 (100%; 95% CI 96.4–100%).

Held-out validation. To confirm that the post-review results generalize beyond the development sample, the reference-discipline and context-fidelity benchmarks were re-run on a fresh held-out sample (seed 20260919, n = 100 each, excluding the development cases). Both held-out runs were 100% (100/100; exact binomial 95% CI 96.4–100%), matching the development results and validating the "≥96% with 95% confidence" claim on unseen cases.

**Classification-layer reporting benchmark (Q&A).** The Q&A submodule reports each gene–form record across seven classification layers and must return the layer the user actually asked for rather than a different layer’s values. The retrieved record now carries the full seven-layer spectrum — Evidence-first (final_role / final_direction), DeepSeek (cot_role / cot_direction), Compiled gene role (rcd_gene_type / role_direction), Consensus (consensus_role / consensus_direction), GPT (gpt_mapped_role / gpt_mapped_dir), Curator 1 (curator_1_mapped_role / curator_1_mapped_dir), and Curator 2 (curator_2_mapped_role / curator_2_mapped_dir) — and the prompt instructs the model to lead with Evidence-first in the synthesis and to report the requested layer verbatim when asked, listing divergent layers in separate rows. Curator vote labels (NOT_NEEDED, OUT_OF_CONTEXT, UNCERTAIN, UNCLEAR) are literal values reported verbatim and never substituted. This was benchmarked on a random, seed-reproducible sample of n = 200 gene–entity pairs (seed 7; 54 divergent and 146 agreeing pairs, matching the 27.2% population divergence rate), each queried for all seven layers (1,400 questions; 1,238 scored after excluding unpopulated layers), with deepseek-chat at temperature 0.3. Overall layer-reporting accuracy was 93.2% (1,154/1,238; Wilson 95% CI 91.7–94.5%). Per layer (n = 200 unless noted): Consensus 98.0% (196/200), Evidence-first 97.0% (194/200), DeepSeek 94.0% (188/200), Curator 1 92.0% (184/200), GPT 88.5% (177/200), Curator 2 88.5% (177/200), and Compiled gene role 100% (38/38 populated). Cross-layer contamination — reporting one layer’s values in answer to another layer’s question — was measured on 100 divergent pairs for the Evidence-first and DeepSeek layers: Evidence-first precision 100% (0/100 substitutions), DeepSeek precision 98% (2/100 substitutions). Residual error concentrated in the GPT and Curator 2 vote layers, the least-populated and least-verbose of the seven.

The board achieves 100% golden accuracy, 100% census accuracy, 100% fact-stability, a 0.047% governance violation rate, and 98.7% articulation conformity, with every violation traceable and logged.

### Supplementary Note S7

### RCDome: AI Usage, Tokens, and Cost:

This note records the compute, token, runtime, and cost values for every AI-assisted phase of the RCDome pipeline. These figures were removed from the main manuscript (Methods, Results, and Discussion) and are documented here for full transparency and reproducibility.

#### 1. Phase 2A/D: Two-call LLM validation (full matrix)

| Rows validated | 21,568 unique gene-RCD pairs |
| --- | --- |
| Runtime | ~35 hours (dedicated server, sequential) |
| DeepSeek API cost | ~US$12 |
| Tokens | 59.6 million |
| Model | deepseek-chat (temperature 0) |

#### 2. Phase 2A/D: MSigDB extension (1,832 pairs)

| Pairs validated | 1,832 MSigDB-only gene-set pairs |
| --- | --- |
| Runtime | ~1–2 hours (8-thread parallel) |
| DeepSeek API cost | ~US$3–5 (estimated) |
| Tokens | 8.2 million |
| Model | deepseek-chat (temperature 0) |

#### 3. Phase 2B: Europe PMC full-text deep RAG

| Articles processed | 167,868 open-access full-text articles |
| --- | --- |
| Pairs with deep evidence | 1,383 rows (1,118 unique pairs) |
| Tokens | 1,383.7 million (phase2b_audit_report.json) |
| Runtime | multi-day (Zima, 8-thread) |
| Cost | not itemized per phase (deepseek-chat) |

#### 4. Phase 2C: Europe PMC full-text recovery (DeepSeek)

| Pairs recovered | 3,058 MECHANISTIC classifications |
| --- | --- |
| Tokens / cost | to be recorded |

#### 5. Phase 2D: NCBI + DeepSeek novel-pair validation

| Pairs validated | 1,341 novel unique pairs |
| --- | --- |
| Tokens / cost | to be recorded |

#### 6. Phase 2E: Europe PMC + DeepSeek full-text recovery

| Pairs processed | 1,724 non-MECHANISTIC pairs |
| --- | --- |
| Recovered | 588 MECHANISTIC associations |
| Tokens / cost | to be recorded |

#### 7. Phase 3: Elicit systematic-review benchmark (3 forms)

| Records extracted | 784 governed records from 154 papers |
| --- | --- |
| Runtime | 5 minutes |
| API cost | ~US$18 (3 forms x ~$6 per form) |

#### 8. Phase 3: Elicit full extraction (50 forms)

| Associations extracted | 4,602 across 50 RCD forms (4,601 retained in Dataset S1) |
| --- | --- |
| Triaged | 3,803 genuine, 798 association_only, 1 retracted |
| Elicit Pro license | US$600.00 |
| Elicit token consumption | US$5,400.00 |

#### 9. Governance literature extraction (DeepSeek, separate track)

| Corpus | 1,500 papers (search + snowball) |
| --- | --- |
| Papers processed | 1,308 |
| Items extracted | 27,852 verbatim-anchored items |
| Raw themes | 3,313 |
| Framework themes | 419 (255 AI-governance, 71 human-clinical, 93 ambiguous) |
| Core themes | 70 |
| Deduplicated themes | 51 |
| Status | COMPLETE |
| Tokens / cost | to be recorded (Zima run logs) |

#### Combined DeepSeek token usage (to date)

| Phase 2A/D (full matrix) | 59.6 M |
| --- | --- |
| Phase 2A/D (MSigDB extension) | 8.2 M |
| Phase 2B deep RAG | 1,383.7 M |
| Phase 2C recovery | included in total |
| Phase 2D NCBI validation | included in total |
| Phase 2E Europe PMC recovery | included in total |
| Governance track | included in total |
| Total (recorded phases) | 7,066,900,638 |

#### Total AI cost summary

| Elicit Pro license | US$600.00 |
| --- | --- |
| Elicit token consumption (full 50-form run) | US$5,400.00 |
| DeepSeek tokens (all phases) | US$2,368.77 |
| Consensus (neutral-query channel) | US$859.25 (US$850.00 paid + US$9.25 upcoming) |
| Total | US$9,228.02 |

Note: token counts are as reported by the DeepSeek API usage fields and the per-run logs. Costs are direct API and license charges; they exclude human time and server infrastructure. The combined DeepSeek usage across the full project (including the manuscript preparation and the audits) totals 7,066,900,638 tokens and US$2,368.77. Phase 2G (Consensus): all 21,568 gene–RCD pairs classified (100%); the 11 consensus columns are merged into the unified matrix, at a Consensus API cost of US$859.25.

### Supplementary Note S8

### RCDome: Anti-Hallucination Strategy for the RCD Entity Descriptions

#### 1. Verification strategy

This note documents the post-hoc anti-hallucination verification of the 53 RCD entity descriptions used in the RCD Entities module, and the re-anchoring pass that followed it. It records the strategy, the steps, the final disposition, and the re-anchoring outcome. No assistant output was accepted on the basis of the assistant alone; every disposition rests on the cited source or on human review.

Every citation-bearing sentence was extracted from the 53 descriptions, yielding 2,812 claims, and each cited reference was resolved to full text through the first available route (Europe PMC open-access XML, OpenAlex, Unpaywall, preprints, Crossref, and PubMed). Each claim was then confronted with the full text of its cited source by an independent checker that quoted the supporting or contradicting sentence.

The verification matched the generation mode. Text extracted verbatim from a source was checked literally. Text synthesized by the assistant was checked by meaning. A first pass was rejected because it used a 6,000 character cap and a one-word answer; the rigorous pass used the complete full text and required the evidence sentence.

#### 2. Seven phases

| Phase | Step | Result |
| --- | --- | --- |
| 1 | Claim extraction and full-text resolution | 2,812 claims |
| 2 | First pass (rejected as shallow) | 1,322 supported, 0 not supported, 1,490 uncertain |
| 3 | Full-text evidence-citing verification | 1,532 supported, 538 not supported, 742 uncertain |
| 4 | Semantic re-verification of paraphrased claims | 120 recovered |
| 5 | Re-sourcing of paywalled-only claims | 198 re-anchored, 213 still unsupported, 153 no public source |
| 6 | Targeted re-anchoring of the unsupported | 141 re-anchored |
| 7 | Semantic Scholar sweep | 3 recovered |

Final disposition: 1,994 supported (70.9%), 284 not supported (10.1%), 534 uncertain (19.0%).

#### 3. Re-anchoring pass

The 284 not-supported and 534 uncertain claims were then re-anchored, reformulated, or removed. The pass verified each claim against the cited source, re-cited claims to references already present in the entity when they supported the claim, re-anchored claims to free peer-reviewed sources when a new source was required, reformulated partial claims to match what the source states, and removed claims without support. Bibliographic flags (retraction, erratum, new version) were also handled.

Result: 669 claims corrected (re-anchored, re-cited, or reformulated), 140 removed, and 9 kept unchanged (false negatives). The final disposition of the 2,812 claims is therefore 2,672 supported (95.0%) and 140 removed (5.0%), with no not-supported or uncertain claim remaining. An independent integrity check then verified the citation-to-reference mapping across the 53 descriptions, and corrected one orphan citation and duplicate references in 15 entities. The module holds 53 descriptions with canonicalized names and 2,129 references, with no orphan citations and no duplicate references.

#### 4. Governance

Every change preserved the original classification, never replaced a reference without archiving, and used only peer-reviewed free sources. The module curator performed the re-anchoring. The four descriptions pending at the time of this note (Autophagy, Programmed cell death, Hypersensitive response, and Eryptosis) were written under the same policy and added to complete the 53 entities.

### Supplementary Prompts

#### Supplementary Prompt S1. Consensus mechanistic-reconstruction framework

\# Regulated Cell Death Mechanism: [PATHWAY NAME]

Provide a comprehensive, mechanistically detailed, evidence-based description of **[PATHWAY NAME]**, reconstructing the process from the earliest initiating signal to the final irreversible death of the cell.

The objective is to explain **how the cell dies as a continuous molecular cause-and-effect sequence**, not merely to list genes, proteins, biomarkers, morphological features, or associated pathways.

Base the answer on peer-reviewed primary studies, authoritative reviews, and established regulated cell death consensus. Prioritize recent evidence when it updates previous models, but include landmark discoveries when necessary. Clearly distinguish established mechanisms, context-dependent findings, hypotheses, and controversies.

1. Definition and classification

Define **[PATHWAY NAME]** and explain:

* what distinguishes it from other regulated cell death mechanisms;

* whether it is an established modality, subtype, or emerging/proposed mechanism;

* its accepted biological definition and main diagnostic criteria;

* the biological contexts in which it occurs.

Separate mechanistically defining features from associated biomarkers or morphology.

2. Initiating signals and sensing

Describe the earliest extracellular or intracellular stimuli that can initiate **[PATHWAY NAME]**.

Include, when relevant, receptor ligands, cellular or oxidative stress, DNA damage, infection, immune signaling, metabolic perturbation, nutrient deprivation, organelle dysfunction, mechanical stress, ion imbalance, and toxic stimuli.

For each major trigger, explain **how the cell senses it** and how the stimulus connects molecularly to downstream death signaling.

Use:

**trigger → sensor/receptor → first intracellular signaling event.**

3. Pathway initiation and propagation

Trace the mechanism step by step after sensing.

Identify the important receptors, adaptor proteins, enzymes, kinases/phosphatases, proteases, transcription factors, metabolites, lipids, ions, organelles, and multiprotein complexes.

For each major component explain:

* what activates or inhibits it;

* its molecular function;

* relevant post-translational modifications;

* changes in localization, oligomerization, or complex formation;

* substrates or interacting partners;

* immediate downstream consequence.

Do not state that a protein simply “regulates” the pathway. Explain **what it physically or biochemically does**.

4. Molecular complexes and biochemical reactions

Describe the central molecular complexes or biochemical reactions required for the pathway.

For each one explain:

* composition;

* cellular location;

* trigger for assembly or activation;

* biochemical activity;

* substrates or targets;

* how the reaction advances the cell toward death.

If multiple complexes form sequentially, describe their chronological and causal relationship.

5. Regulatory checkpoints and protective systems

Identify mechanisms that normally prevent inappropriate activation of **[PATHWAY NAME]**, including relevant endogenous inhibitors, anti-death proteins, antioxidant systems, metabolic defenses, repair systems, degradation pathways, and feedback loops.

For each, explain:

**protective factor → molecular target → inhibitory mechanism → consequence of failure.**

State whether loss of the protective mechanism directly triggers death or only sensitizes the cell.

6. Commitment and reversibility

Identify the known or proposed **point of no return**.

Explain:

* which stages represent reversible stress or pathway activation;

* which events represent commitment to death;

* evidence that cells can or cannot recover after these events;

* genetic or pharmacological interventions that rescue cells before or after commitment.

If no single commitment point is known, state this explicitly.

7. Execution phase

Describe the execution phase in greatest mechanistic detail.

Answer:

**What molecular process physically causes the cell to lose viability in [PATHWAY NAME]?**

Explain sequentially all relevant events, such as plasma membrane permeabilization or rupture, pore formation, proteolysis, mitochondrial dysfunction, ATP depletion, metabolic collapse, lipid peroxidation, ROS accumulation, ion dysregulation, osmotic imbalance, lysosomal or ER damage, DNA/chromatin alterations, cytoskeletal collapse, and organelle failure.

Include only processes supported for this mechanism and explain how one event causes the next.

8. Terminal cause of death

Explicitly identify the **proximate cause of cellular death**.

Explain what ultimately makes recovery impossible.

Distinguish:

* molecular execution mechanism;

* irreversible loss of homeostasis;

* morphological consequences;

* biomarkers;

* events occurring after commitment or death.

Clarify whether the terminal event is membrane rupture, catastrophic lipid damage, proteolytic dismantling, energetic failure, osmotic collapse, organelle failure, or another mechanism.

9. Morphological progression

Describe morphology chronologically and connect each feature to its molecular cause:

**healthy cell → early response → commitment → execution → terminal morphology.**

Include relevant features such as swelling or shrinkage, membrane blebbing, chromatin condensation, nuclear fragmentation, vacuolization, mitochondrial or other organelle changes, apoptotic bodies, and membrane rupture.

State which features are characteristic and which are shared with other death mechanisms.

10. Canonical and alternative routes

If more than one molecular route leads to **[PATHWAY NAME]**, describe each separately using:

**trigger → sensor → signaling intermediates → molecular complex/reaction → executioner → terminal event.**

Explain where routes diverge and converge and whether they are cell-type-, stimulus-, or context-dependent.

Use “canonical” or “non-canonical” only when supported by the literature.

11. Crosstalk with other regulated cell death mechanisms

Describe experimentally demonstrated crosstalk with other death modalities.

Focus on shared signaling proteins, enzymes, metabolites, organelles, stress responses, or execution machinery.

For each connection, specify whether it represents pathway switching, shared upstream signaling, shared execution machinery, reciprocal inhibition, or simultaneous activation.

Do not treat correlation or co-occurrence as mechanistic crosstalk.

12. Experimental evidence

Explain how researchers demonstrate that death occurs specifically through **[PATHWAY NAME]**.

Discuss the strongest relevant approaches, including:

* genetic knockout/knockdown;

* gain-of-function;

* pathway-specific inhibitors;

* rescue experiments;

* biochemical assays;

* microscopy;

* membrane integrity assays;

* organelle-specific assays;

* metabolomics, lipidomics, or proteomics when relevant.

Explain which experiments distinguish this pathway from alternative death mechanisms.

Separate causal evidence from biomarkers and morphology.

13. Molecular hierarchy

Classify the principal molecules as:

1. **Initiators/sensors** – detect the stimulus.

2. **Signal transducers** – propagate signaling.

3. **Core machinery** – required molecular components.

4. **Executioners** – directly produce lethal damage.

5. **Negative regulators/protective factors** – suppress the pathway.

6. **Biomarkers** – indicate activation but may not be causal.

7. **Context-dependent regulators** – important only in specific systems.

For every core molecule, state its precise mechanistic role.

14. Final integrated mechanistic sequence

Conclude with one continuous chronological causal chain:

**Initiating stimulus**

→ cellular sensor/receptor

→ proximal signaling

→ molecular complex or biochemical reaction

→ amplification/regulatory events

→ commitment to death

→ executioner mechanism

→ structural/metabolic failure

→ irreversible loss of cellular homeostasis

→ cell death.

**This final sequence must be detailed enough to be converted directly into a scientific pathway diagram, mechanistic figure, or biological animation.**

**Evidence standards**

**Throughout the answer:**

* cite major mechanistic claims;

* prioritize primary experimental evidence;

* distinguish established facts from proposed mechanisms;

* identify unresolved or controversial steps;

* do not infer causality from gene-expression correlations alone;

* state when evidence is species-, cell-type-, or stimulus-specific;

* prioritize mammalian and human evidence when available;

* do not describe biomarkers as causal machinery without experimental support.

The final objective is to reconstruct [PATHWAY NAME] from the first biological signal to the physical and irreversible death of the cell, explaining every major causal molecular step in between.

#### Supplementary Prompt S2. Harmonization framework

Using only the text and references provided below, write a detailed, publication-ready mechanistic description of [PATHWAY NAME] in English for the RCDome**.**

Follow this structure:

1. Definition and Classification — state whether it is established, emerging, provisional, subtype, process, trigger, or RCD-related concept. Clearly distinguish what is consensus from what is recent, context-dependent, or unresolved.

2. Initiation and Triggers — explain the physiological/pathological context, trigger, sensor/receptor, and earliest known intracellular event. If the trigger or sensor is unknown, state this explicitly.

3. Core Mechanistic Machinery — describe the pathway continuously from trigger to signaling, molecular complexes, organelles, biochemical reactions, and structural changes. Explain the precise role of each major molecule; do not call a regulator an executor unless supported.

4. Propagation and Amplification — explain important branches, feedback loops, metabolic/ionic changes, organelle remodeling, and context-dependent regulators.

5. Protective/Negative Regulators — describe mechanisms that suppress, delay, repair, or reverse the pathway.

6. Commitment / Point of No Return — identify the best-supported irreversible step. If no universal commitment point is known, say so and give the strongest current candidate.

7. Execution and Terminal Cause of Death — answer explicitly: What physically kills the cell? Distinguish the core executor from downstream morphology or biomarkers. If the terminal mechanism is unresolved, do not invent one.

8. Morphological Sequence — describe the chronological cellular changes from a normal cell to terminal death.

9. Crosstalk and Differential Diagnosis — distinguish this pathway mechanistically from the most similar RCD modalities.

10. Experimental Validation — explain the strongest experiments required to prove this pathway, including genetic perturbation, pharmacological rescue, functional assays, imaging, and exclusion of competing death mechanisms.

11. Molecular Hierarchy — summarize: initiators/sensors → signal transducers → core machinery → executors → negative regulators → biomarkers.

12. Integrated Canonical Sequence — finish with one continuous arrow-style mechanistic sequence from the first trigger to terminal death.

13. Final Mechanistic Interpretation — provide the most defensible operational definition for RCDome, including evidence status, biological context/species if relevant, core machinery, terminal lesion, and major unresolved question.

14. References — use numbered citations [1], [2], etc. throughout the text and provide a numbered reference list with DOI at the end.

**Critical rules**

* Do not invent mechanisms, molecules, receptors, executors, or points of no return.

* Separate established core, context-dependent/emerging evidence, and hypotheses/unresolved mechanisms.

* Morphology or biomarkers alone must not be treated as proof of mechanism.

* Do not infer active cell death from gene-expression signatures alone; when relevant, describe transcriptomic data as pathway-associated or susceptibility states.

* Avoid tables unless absolutely necessary.

* Preserve mechanistic precision and write as a continuous biological story rather than a list of disconnected facts.

* If the supplied references conflict, explain the uncertainty instead of choosing one without justification.

* Do not use outside references unless explicitly requested.

**SOURCE TEXT AND REFERENCES**

**[PASTE TEXT + REFERENCES HERE]**

#### Supplementary Prompt S3. Figure-generation framework

**# PROMPTS**

**1. FIGURE CREATION**

Create a professional scientific illustration of [MECHANISM NAME] for publication in a high-impact peer-reviewed journal such as *Nature Cell Biology*, *Cell Death & Differentiation*, or *Nature Reviews Cancer*. The visual style must follow the standards of professional biomedical scientific illustration — precise, informative, and elegant.

**CONCEPT TO ILLUSTRATE:**

**[PASTE MECHANISM DESCRIPTION HERE]**

**MANDATORY VISUAL REQUIREMENTS:**

* White background, no gradients, no dark themes, no cinematic effects, no glow, no lens flares

* Clean vector-like illustration style, semi-3D organelle shapes with soft shading for depth — reference style: *Nature Reviews* figures, BioRender premium, or Servier Medical Art

* All molecular components and cellular structures must be anatomically and conceptually accurate — no decorative invention

* Color palette: use distinct, harmonious, muted scientific colors — one consistent color per molecular actor throughout the figure (e.g., always the same color for caspases, for mitochondria, for the nucleus, etc.)

* All key structures must be labeled with clean black sans-serif text (Arial or Helvetica style) connected by thin leader lines

* Arrows must indicate directionality of the pathway (activation = solid arrow $\rightarrow$, inhibition = flat-head bar $\dashv$, crosstalk = dashed arrow)

* The cell must be shown in cross-section, revealing internal organelles relevant to the mechanism

* Include a small inset panel in the bottom-right corner showing the cancer relevance described in the concept: either evasion, promotion, or therapeutic exploitation — labeled "Cancer Context"

* If the mechanism shares pathways with other cell death forms, show these connections as light gray dashed arrows with small labels at the periphery of the figure, without overloading the main illustration

* Layout must be organized left to right or top to bottom following the biological sequence of the pathway

* Final image must look publication-ready at 600 DPI, suitable as a figure panel or journal cover candidate.

**STYLE REFERENCES (mandatory):**

> "Reference style: scientific figure illustration from Nature Reviews Cancer or Cell journal, drawn by a professional biomedical illustrator, white background, flat-to-semi-3D organelle style, no dark themes, no futuristic aesthetics, no glowing effects. BioRender premium quality."

**DO NOT include:**

* Dark or black backgrounds

* Neon or glowing effects

* Futuristic or sci-fi aesthetics

* Decorative elements not grounded in the biology

* Generic "cell death" imagery not specific to the described mechanism

* Text outside of labels (no titles, no legends inside the image)

**2. AUDIT**

**SCIENTIFIC ACCURACY AUDIT for [MECHANISM NAME]**

Act as a senior peer reviewer specializing in regulated cell death (RCD), at the standard of high-impact journals such as *Nature Cell Biology*, *Cell Death & Differentiation*, *Molecular Cell*, or *Nature Reviews Cancer*. Your task is to rigorously audit the conceptual, logical, and sequential accuracy of the RCD pathway described/illustrated below, prior to approval for publication**.**

**RCD PATHWAY UNDER AUDIT:**

[PATHWAY NAME — e.g., Necroptosis, Pyroptosis, Ferroptosis, PANoptosis, Intrinsic Apoptosis, etc.]

**CONTENT TO BE AUDITED (textual description and/or generated image):**

[PASTE MECHANISM DESCRIPTION AND/OR DESCRIBE WHAT THE IMAGE SHOWS HERE]

**MANDATORY AUDIT SCOPE — verify each of the following axes:**

1. Causal sequence (central pathway)

Are events in the correct chronological/causal order (sensor $\rightarrow$ adaptor $\rightarrow$ effector $\rightarrow$ execution)? Does every activation or inhibition arrow reflect a real molecular relationship documented in the primary literature? Are there no "logical jumps" (omitted obligatory intermediate steps that would change the mechanistic meaning)? Do multiprotein complexes (e.g., necrosome, inflammasome, apoptosome, ripoptosome) show the correct stoichiometric composition and correct partners?

2. Molecular identity and specificity

Is each protein/molecule cited specific to this pathway, or is it shared/redundant with another RCD pathway (and is this correctly flagged as crosstalk rather than exclusive)? Are isoforms and activation states (cleaved vs. precursor; phosphorylated vs. non-phosphorylated; oligomerized vs. monomeric) correctly differentiated? Is the subcellular localization of each component anatomically correct (cytosol, outer mitochondrial membrane, plasma membrane, nucleus, ER, etc.)?

3. Directionality and arrow type

Is every activation shown as a solid arrow ($\rightarrow$) and every inhibition as a flat-head bar ($\dashv$)? Is there no sign inversion (e.g., an inhibitor shown as an activator, or vice versa)? Are bidirectional or feedback relationships (positive/negative) explicitly marked as such?

4. Peripheral context (crosstalk with other pathways)

Are connections to other forms of cell death (apoptosis, necroptosis, pyroptosis, ferroptosis, autophagy, PANoptosis, etc.) biologically validated and not speculative? Are decision/branch points (e.g., caspase-8 as a switch between apoptosis and necroptosis) correctly positioned within the pathway, rather than presented as exclusively central to this pathway? Is there no improper overlap between "central pathway" and "peripheral crosstalk" (i.e., a central component cannot be treated as peripheral, and vice versa)?

5. Cancer context (inset)

Is the described role (evasion, promotion, or therapeutic exploitation) consistent with current literature and not an overgeneralization? Is the cited oncological relevance specific to this pathway (and not a generic cell-death effect)? If a drug/inhibitor/agonist is cited as therapeutic exploitation, does it actually act on the indicated molecular target, at the correct step of the pathway?

6. Terminological consistency

Does the nomenclature follow the most recent Nomenclature Committee on Cell Death (NCCD) guidelines? Are terms not being used as synonyms when they actually describe distinct processes (e.g., "necrosis" vs. "necroptosis"; "immunogenic cell death" treated as a separate pathway rather than a phenotype)?

7. Critical omissions

Is there any essential regulator (key activator or inhibitor) missing whose absence would change the interpretation of the pathway? Is there any relevant feedback loop that was omitted?

**#MANDATORY OUTPUT FORMAT:**

For each of the 7 axes above, answer in table format with the columns:

| Axis | Status (✅ Correct / ⚠️ Imprecise / ❌ Incorrect) | Identified problem | Why it is wrong (conceptual rationale) | How to correct it (specific, actionable fix) |

Rule: Every row marked ⚠️ or ❌ MUST include a concrete, actionable correction in the last column — never leave a problem flagged without a corresponding fix. The correction must specify exactly what changes (e.g., *"replace inhibitory bar with activating arrow between X and Y"*, *"reposition complex Z from cytosol to outer mitochondrial membrane"*, *"relabel as cleaved-caspase-3 instead of pro-caspase-3"*, *"move this connection from central pathway to peripheral crosstalk and mark with gray dashed arrow"*).

At the end, provide:

1. An overall verdict: `PUBLISHABLE` / `PUBLISHABLE WITH MINOR CORRECTIONS` / `REQUIRES MAJOR REVISION` / `REJECT`

2. If any inconsistency was found: a fully corrected and rewritten version of the mechanistic description, with every fix from the table already incorporated, ready to be used as the new input for the `[CONCEPT TO ILLUSTRATE]` field of the image-generation prompt.

3. If any inconsistency was found: a short bullet-point "fix list" of only the elements that need to be physically changed in the image itself (position, arrow type, label text, color, removal or addition of a component) — written so it can be pasted directly as an instruction for image regeneration.

4. If no inconsistency was found: explicitly state *"No corrections needed — pathway is accurate as described"* and skip points (2) and (3).

*Be rigorous and skeptical by default: treat any claim that is not broadly consolidated in the literature as "imprecise" until proven otherwise, even if it seems plausible. Do not soften critiques out of courtesy — the standard is that of a demanding Q1-journal reviewer. Never flag a problem without simultaneously providing its solution.*

#### Supplementary Prompt S4. Multi-model scientific-accuracy audit framework

UNIVERSAL FIGURE AUDIT & CORRECTION PROMPT

Act as a senior peer reviewer and scientific illustration editor for a high-impact journal (Nature Cell Biology, Cell Death & Differentiation, Molecular Cell, Nature Reviews Cancer), with expertise in regulated cell death (RCD) pathways and in reviewing biomedical scientific illustrations.

Your task has two parts, to be executed in sequence:

(1) Audit the figure below against the current literature and against the mandatory visual standards of this journal.

(2) Based strictly on the errors you find, produce a corrected, ready-to-use version of both the mechanistic description and the image-generation prompt, so the figure can be regenerated as conceptually and visually accurate.

**INPUT:**

- Mechanism name: [MECHANISM NAME]

- Mechanistic description used to generate the figure: [PASTE DESCRIPTION USED]

- Original image-generation prompt used: [PASTE ORIGINAL IMAGE PROMPT, IF AVAILABLE]

- Figure to audit: [ATTACH/DESCRIBE THE IMAGE]

MANDATORY VISUAL STANDARD (the corrected figure must always comply with this, regardless of mechanism):

- White background, no gradients, no dark themes, no glow, no lens flares, no cinematic or sci-fi effects

- Clean vector-like, semi-3D organelle style (BioRender premium / Nature Reviews Cancer standard)

- One consistent color per molecular actor throughout the entire figure

- All structures labeled with clean black sans-serif text and thin leader lines

- Activation = solid arrow (→); inhibition = flat-head bar (⊣); crosstalk with other cell-death pathways = light gray dashed arrow at the periphery only

- Cross-sectional cell view, anatomically correct subcellular localization for every component

- Correct stoichiometry and composition of multiprotein complexes (e.g., necrosome, inflammasome, apoptosome, ripoptosome)

- Layout follows the biological sequence, left-to-right or top-to-bottom

- Small labeled inset, bottom-right, titled "Cancer Context," showing evasion, promotion, or therapeutic exploitation relevant to this specific mechanism

- No decorative or invented elements without biological grounding; no text beyond labels

- Terminology consistent with the current NCCD (Nomenclature Committee on Cell Death) guidelines

**AUDIT**

Check every element of the figure against these axes, treating any claim not solidly established in the literature as unverified until confirmed:

1. Causal sequence — correct sensor → adaptor → effector → execution order; no omitted obligatory steps; correct complex composition

2. Molecular identity — correct activation state (cleaved vs. precursor, phosphorylated vs. not, oligomerized vs. monomeric) and correct subcellular localization for every labeled component

3. Arrow accuracy — for every single arrow/connector: correct type (activation vs. inhibition), correct direction, correctly marked if bidirectional/feedback

4. Crosstalk — connections to other RCD pathways are peripheral, gray, dashed, and biologically validated (not central, not speculative)

5. Cancer context — inset content is specific to this mechanism and consistent with current literature, not generic

6. Color and label consistency — one color per actor throughout; every label attached to the correct structure via its leader line

7. Style compliance — matches the mandatory visual standard above with no exceptions

8. Critical omissions — any essential regulator, feedback loop, or step missing that would change the interpretation of the pathway

Be rigorous and skeptical by default. Flag anything imprecise, even if plausible, until it is clearly supported. Never note a problem without also stating exactly how to fix it.

**OUTPUT FORMAT (always structured exactly like this):**

**Audit Findings**

| Axis | Status (✅/⚠️/❌) | Element affected | Problem | Why it is wrong | Exact fix |

**Overall Verdict**

**PUBLISHABLE / PUBLISHABLE WITH MINOR CORRECTIONS / REQUIRES MAJOR REVISION / REJECT**

**Corrected Mechanistic Description**

(Full rewritten description, only if any issue was found — with every fix already incorporated. If none found, write: "No corrections needed — mechanism and figure are accurate as described.")

**Corrected Image-Generation Prompt**

(A complete, ready-to-submit prompt combining: the corrected mechanistic description above + the full mandatory visual standard listed above + a short explicit "Corrections applied" section listing only what changed from the previous version, so the differences are traceable. Skip this section if no issue was found.)

Do not simplify, shorten, or soften any part of the corrected prompt — it must be complete and self-sufficient for direct resubmission to the image generator, requiring no further editing.

#### Supplementary Prompt S5. Figure-legend generation framework

\# PROMPT: FIGURE LEGEND GENERATION FOR REGULATED CELL DEATH (RCD) ILLUSTRATIONS

You are a senior scientific editor and author specializing in regulated cell death (RCD), writing at the standard of *Nature Cell Biology*, *Cell Death & Differentiation*, *Molecular Cell*, and *Nature Reviews Cancer*. Your task is to write a publication-ready figure legend for the scientific illustration provided, following the technical conventions of scientific writing described in *How to Write and Publish a Scientific Paper* (Gastel & Day).

**INPUTS (all three are mandatory)**

1. FIGURE: the attached image of the mechanism.

2. FIGURE-GENERATION PROMPT: the prompt used to create the figure (visual conventions: arrow types, color logic, inset "Cancer Context", gray dashed crosstalk, left-to-right/top-to-bottom flow).

[PASTE OR REFERENCE THE FIGURE-CREATION PROMPT]

3. MECHANISM DESCRIPTION: the text used as the [CONCEPT TO ILLUSTRATE].

[PASTE MECHANISM DESCRIPTION HERE]

RCD PATHWAY: [e.g., Ferroptosis / Alkaliptosis / Disulfidptosis / Necroptosis / Pyroptosis / PANoptosis ...]

TARGET JOURNAL (optional): [journal name]

WORD LIMIT (optional): [e.g., 250–350 words; if not specified, aim for 200–350 words]

FIGURE NUMBER (optional): [e.g., Figure 1]

**SOURCE HIERARCHY AND ANTI-HALLUCINATION RULES**

- Use ONLY information present in the figure and the mechanism description. Do not add molecules, steps, drugs, citations, or claims that appear in neither.

- If the figure and the description disagree (a label, arrow type, component, or localization), do NOT silently choose one. Follow the mechanism description for biological content, follow the figure for what is physically displayed, and report every discrepancy in the "Discrepancy Report" (see output).

- Preserve the evidentiary qualifiers present in the figure or description ("proposed", "candidate", "context-dependent", "conceptual", "not established", "preliminary", "not required for initiation"). Never convert a hedged claim into an established one. Never upgrade correlation to causation.

- Do not invent citations, PMIDs, or reference numbers. Place "(refs)" markers only if the user explicitly requests them.

**STRUCTURE OF THE LEGEND (follow this order)**

1. Title (bold, one sentence, ≤ 20 words). Informative and specific to the pathway, stating the main message of the figure, not merely naming the topic (e.g., "Ferroptosis is driven by iron-dependent phospholipid peroxidation when GPX4-independent defenses fail"). No period issues, no abbreviations unless universally known.

2. Overview sentence (1–2 sentences). State what the figure shows as a whole: the trigger, the executor/mechanism of death, and the outcome. Use the present tense for the model (schematic figures depict established or proposed models, not experimental results).

3. Panel-by-panel (or region-by-region) description. Describe each numbered panel/module in the same sequence as the biological flow of the figure (e.g., induction → regulation → execution → death). For each panel:

- Identify it by its number/letter exactly as shown in the figure (e.g., "(1)", "(a)").

- Name the key molecular actors and their subcellular localization as drawn (plasma membrane, cytosol, lysosome, mitochondrion, nucleus, ER, etc.).

- State causal relationships with correct verbs (activates, inhibits, promotes, is required for, amplifies, is dispensable for), consistent with the arrow types in the figure.

- Clearly distinguish core pathway components from modulators, amplifiers, and non-essential/peripheral events, mirroring how the figure separates them.

- Distinguish activation states when shown (cleaved vs. precursor, phosphorylated, oligomerized, oxidized/reduced).

4. Crosstalk statement (1–2 sentences). Describe the connections to other RCD forms exactly as depicted (light gray dashed arrows, peripheral boxes). State explicitly whether crosstalk is validated, reciprocal, conceptual, or absent, and that these are not part of core execution when the figure says so.

5. Cancer Context statement (1–3 sentences). Summarize the inset: whether the pathway is exploited for evasion, promotion, or therapy; name only the tumor types, drugs, and targets shown; keep the evidence-level qualifiers (e.g., "best-established", "preclinical", "preliminary/associative").

6. Symbol and color key (1–2 sentences, placed inside the legend, NOT inside the image). Define every graphical convention used: solid arrows = activation/flow; flat-head bars = inhibition; dashed arrows = crosstalk or conceptual/indirect relationships; gray dashed = connections to other RCD forms; consistent color for each molecular actor class; any special symbols (P = phosphate, S–S = disulfide bond, red cross = blocked/not required, "?" = uncertain/unestablished). Include only conventions actually present in the figure.

7. Abbreviations. Define every abbreviation at first use in the legend or, if more than ~8 are needed, provide a single alphabetized sentence at the end: "Abbreviations: ...". Use official HGNC gene/protein symbols as they appear in the figure (italicize gene symbols only if the figure treats them as genes; keep protein symbols in roman type). Follow current NCCD nomenclature for cell death terms and do not use distinct processes as synonyms (e.g., necrosis vs. necroptosis).

8. Adaptation/attribution line (optional). If the figure is original and created by the authors, add: "Figure created by the authors." If adapted from other work, add: "Adapted from [source], with permission." Include only if the user supplies this information.

**WRITING RULES (Gastel & Day principles)**

- The legend must be self-explanatory: a reader should understand the figure without reading the main text.

- Be concise, precise, and non-redundant: do not repeat information already readable from the labels in the image, do not restate general background, and do not discuss methods or results beyond what the schematic shows.

- Use simple, direct sentences and the present tense for model descriptions; avoid vague words ("various", "several", "etc.") and avoid unnecessary adjectives or promotional language ("remarkable", "crucial", "novel").

- Use consistent terminology with the figure: the same name for the same entity throughout.

- Write full words for pathway names at first use, with the abbreviation in parentheses.

- Use correct units, ion notation (Ca²⁺, Fe²⁺, H⁺, HCO₃⁻), and symbols (↑, ↓, →, ⊣) consistently.

- Do not include a "Figure X." prefix duplication; if a figure number is provided, begin with it in bold followed by the title.

- Respect the word limit; if the content cannot fit, prioritize in this order: core pathway → regulation/modulators → cancer context → crosstalk → abbreviations.

**OUTPUT FORMAT (mandatory)**

**A. FINAL FIGURE LEGEND**

The complete legend, ready to paste into a manuscript, as continuous prose (title in bold, then paragraph text; no bullet points inside the legend).

**B. ABBREVIATIONS (only if not already integrated into the legend)**

**C. DISCREPANCY REPORT**

A table with the columns:

| Item | What the figure shows | What the description says | Resolution adopted in the legend | Recommended action (fix in image or in text) |

**If there are no discrepancies, write: "No discrepancies detected between figure, generation prompt, and mechanism description."**

**D. QUALITY CHECKLIST**

Confirm each item with ✅ / ⚠️ and a short note:

- Title is informative and specific

- Panel order follows the biological flow of the figure

- All labeled structures relevant to the mechanism are covered

- All arrow types and colors are explained

- All abbreviations are defined

- Evidence-level qualifiers preserved (no overstatement)

- Crosstalk described as shown (peripheral, not central)

- Cancer Context consistent with the inset

- NCCD-consistent terminology

- No information added beyond figure + description

- Word limit respected (state the final word count)

Be rigorous and conservative: when in doubt, preserve the hedge, flag the issue in the Discrepancy Report, and never add unsupported information.

#### Supplementary Prompt S6. Logo (icon) creation framework

Create a new image from scratch.

You are a senior biomedical illustrator creating ONE minimalist scientific icon for a coherent collection of Regulated Cell Death (RCD) modalities intended for publication in a high-impact biomedical journal.

You will receive only the biological description of one RCD pathway.

First interpret the biology silently. Then generate the icon directly.

**IMPORTANT:**

This is NOT a mechanistic pathway diagram.

This is NOT an infographic.

This is NOT a molecular illustration.

This is NOT a logo in the commercial-branding sense.

It is a SIMPLE CELLULAR PICTOGRAM representing the characteristic phenotype or defining biological event of the RCD modality.

**==================================================**

**CORE VISUAL LANGUAGE — MUST REMAIN CONSTANT**

**==================================================**

The entire RCD collection follows the same visual grammar:

- flat 2D vector illustration;

- front-facing or top-down cellular view;

- simple soft cartoon-like biomedical style;

- rounded organic shapes;

- thick, smooth, uniform outlines;

- no perspective;

- no isometric view;

- no 3D;

- no realistic rendering;

- no gradients;

- no shadows;

- no textures;

- pure white background;

- square canvas;

- one centered composition;

- generous white margin;

- simple enough to remain readable at thumbnail size.

The image must look like a member of ONE unified scientific icon family.

**==================================================**

**DEFAULT CELL TEMPLATE**

**==================================================**

Unless the biology clearly requires another arrangement, represent the RCD as ONE large simplified cell occupying approximately 70–85% of the canvas.

The default cell should have:

- a large rounded, oval, or mildly irregular outer silhouette;

- one thick continuous membrane outline;

- one simple nucleus when biologically appropriate;

- a few highly simplified intracellular structures;

- sparse small circular cytoplasmic dots when useful;

- approximately two simplified mitochondria when appropriate;

- several broad curved ER-like bars around the nucleus when appropriate.

Internal organelles are SECONDARY visual elements.

They should be subtle and simplified rather than competing with the defining RCD feature.

Do NOT automatically add every organelle.

**==================================================**

**PATHWAY-SPECIFIC MORPHOLOGY**

**==================================================**

Read the supplied biological description and identify the SINGLE most characteristic morphological, structural, or cellular event of this RCD.

Modify the default cell template only as much as necessary to express that event.

Possible changes include, only when supported by the description:

- swelling;

- shrinkage;

- irregular contour;

- membrane rupture;

- membrane discontinuity;

- blebbing;

- fragmentation;

- vacuolization;

- organelle alteration;

- nuclear alteration;

- extrusion;

- engulfment;

- cell-in-cell formation;

- extracellular release;

- DNA/chromatin release;

- detached fragments;

- loss of membrane integrity;

- accumulation of a specific cellular compartment.

The defining event must be immediately visible.

However, preserve the SAME simplified visual language used for the rest of the collection.

**==================================================**

**EXCEPTION TO THE ONE-CELL TEMPLATE**

**==================================================**

Use more than one cell, extensive extracellular material, or an unusual composition ONLY when the defining biology cannot be communicated correctly with one cell.

Examples include processes involving:

- cell-in-cell structures;

- engulfment;

- phagocytosis/efferocytosis;

- extrusion of extracellular chromatin;

- interaction between two cells;

- massive release of cellular material.

Even in these cases, preserve the exact same flat, rounded, simplified cartoon-like visual grammar.

Do NOT transform the image into a mechanistic diagram.

**==================================================**

**COLOR LANGUAGE**

**==================================================**

Color may vary between RCD modalities.

Do NOT force all icons to use the same color.

Choose a visually distinctive dominant color family appropriate for THIS icon.

Examples may include:

blue,

teal,

green,

yellow,

orange,

brown,

red,

pink,

magenta,

purple,

gray,

or another restrained scientific color.

The icon may be:

- predominantly monochromatic with several related tonal values; OR

- use 2–3 harmonious colors when biologically useful.

Prefer ONE dominant color family.

Use additional contrasting colors only when necessary to distinguish an important pathway-specific structure or event.

Do NOT create a highly multicolored illustration.

The cell body, nucleus, ER, mitochondria, particles and other structures should usually be rendered as tonal variations of the dominant palette.

Outlines must always be a darker, more saturated version of the corresponding fill.

Never use pure black outlines.

**==================================================**

**OUTLINE AND SHAPE STYLE**

**==================================================**

Use a clearly visible thick outer contour.

The outer contour should be approximately 1.5–2.5% of the main cell width.

Use smooth rounded curves.

Internal structures should use the same rounded visual language.

Avoid:

- delicate linework;

- thin technical lines;

- realistic membranes;

- complex bilayers;

- excessive nested borders;

- sharp geometric shapes unless biologically necessary.

**==================================================**

**NUCLEUS**

**==================================================**

When present, the nucleus should be:

- one large simple circle, oval, or softly irregular shape;

- filled with a darker or lighter tone from the same palette;

- surrounded by one simple outline.

Do NOT draw chromatin detail unless nuclear alteration is itself the defining feature.

**==================================================**

**MITOCHONDRIA**

**==================================================**

When mitochondria are present as background organelles:

- show only approximately 1–2;

- use simple bean-shaped silhouettes;

- use a few short rounded internal cristae;

- keep them visually subtle and tone-on-tone.

If mitochondrial morphology is the defining feature of the pathway, it may become more prominent.

Do NOT create a detailed standalone mitochondrion unless absolutely required by the biological description.

**==================================================**

**ENDOPLASMIC RETICULUM**

**==================================================**

When appropriate, represent the ER as a few broad smooth curved bars arranged around part of the nucleus.

Use approximately 3–6 rounded segments.

Keep them tone-on-tone and visually secondary.

Do NOT draw realistic ER networks.

**==================================================**

**SMALL CYTOPLASMIC ELEMENTS**

**==================================================**

Small molecules, granules, vesicles or particles may be represented using a small number of:

- circles;

- rounded dots;

- simple blobs;

- compact geometric shapes.

Use them sparingly.

They are decorative-supporting biological elements, not the main subject unless explicitly required by the pathway.

**==================================================**

**MEMBRANE DAMAGE OR CELL RUPTURE**

**==================================================**

When membrane integrity is lost:

- break the outer contour locally;

- use a few simplified released fragments or soft extracellular blobs;

- optionally use a dashed contour in damaged regions if appropriate;

- preserve the same thick rounded line style.

Do NOT render realistic membrane debris.

Do NOT create explosive visual effects.

**==================================================**

**EXTRACELLULAR RELEASE**

**==================================================**

**When cellular material is released:**

**- use broad soft pale shapes;**

**- use only a limited number of fragments, dots, vesicles or strands;**

**- maintain ample white space;**

**- keep the composition clean.**

**For extracellular DNA or chromatin, use a few simple thin wavy strands only when biologically central.**

**==================================================**

**VISUAL HIERARCHY**

**==================================================**

The illustration should contain:

1. ONE dominant cell or cellular event;

2. ONE immediately recognizable RCD-specific morphological hallmark;

3. only a few supporting structures.

The defining event must carry the identity of the pathway.

Background organelles must remain secondary.

Do NOT attempt to explain the complete signaling mechanism.

**==================================================**

**STRICTLY PROHIBITED**

**==================================================**

NO text.

NO pathway name.

NO protein names.

NO gene names.

NO chemical names.

NO ion labels.

NO abbreviations.

NO NH3, Ca2+, ROS, ATP, DNA labels, etc.

NO letters.

NO numbers.

NO legends.

NO annotations.

NO arrows.

NO arrowheads.

NO cause-to-effect diagram.

NO signaling pathway.

NO flowchart.

NO isolated organelle connected to another structure by arrows.

NO realistic molecular structures.

NO detailed molecular interactions.

NO perspective.

NO tilted cells.

NO isometric illustration.

NO 3D organelles.

NO depth.

NO gradients.

NO shadows.

NO highlights.

NO gloss.

NO texture.

NO photorealism.

NO decorative elements unrelated to the biological description.

**==================================================**

**SCIENTIFIC INTERPRETATION**

**==================================================**

Use ONLY the information contained in the supplied RCD description.

Do not invent morphological characteristics.

If the description contains many molecular mechanisms but only one characteristic cellular phenotype, represent the CELLULAR PHENOTYPE rather than attempting to draw all molecular mechanisms.

If no unique morphology is described, choose the simplest cellular representation of the most distinctive biological event.

Translate molecular mechanisms into visible cellular consequences whenever scientifically justified.

For example:

a molecular alteration should NOT automatically be represented by labeled molecules and arrows.

Instead, represent the cellular or organellar state produced by that alteration when supported by the description.

**==================================================**

**SERIES CONSISTENCY**

**==================================================**

Imagine that this icon will appear directly beside dozens of other RCD icons.

It must have the same:

- line thickness;

- degree of simplification;

- cartoon-like softness;

- organelle abstraction;

- visual density;

- flatness;

- white background;

- overall scale;

- margin;

- level of biological detail.

The PATHWAY changes.

The STYLE does not.

Different RCDs should be distinguished mainly by:

- cell morphology;

- defining structural event;

- extracellular behavior;

- specific organelle alteration;

- silhouette;

- dominant color family.

They must NOT be distinguished by changing illustration style.

**==================================================**

**FINAL QUALITY CONTROL**

**==================================================**

Before generating the image, silently ask:

1. Does this look like a simple cell icon rather than a mechanistic diagram?

2. Is there one dominant cellular phenotype?

3. Are arrows, labels and molecular notation completely absent?

4. Is the illustration strictly flat 2D?

5. Is the outline thick, smooth and rounded?

6. Is the palette restrained, preferably one dominant color family?

7. Are internal organelles simple and secondary?

8. Could this image sit beside a blue, green, red, brown, purple or pink version of the same visual family without looking stylistically different?

9. Is the biological hallmark recognizable without text?

10. Would the icon remain readable when strongly reduced in size?

If any answer is NO, simplify and correct the image before generating it.

**==================================================**

**OUTPUT**

**==================================================**

ONE isolated RCD scientific icon.

Square canvas.

Pure white background.

Centered composition.

Large cellular subject.

Generous clean margin.

Flat 2D vector.

Soft rounded shapes.

Thick smooth colored outlines.

Restrained palette.

Minimalist biomedical cartoon style.

Publication-quality consistency.

**PATHWAY DESCRIPTION:**

[PASTE THE COMPLETE RCD DESCRIPTION HERE]

### SUPPLEMENTARY FIGURE LEGENDS

**Supplementary Figure S1. Database contribution and multi-database corroboration.** (A) Number of gene–RCD associations contributed by each of the 20 source databases and resources (log2-scaled axis; exact counts printed at the end of each bar). (B) Number of source databases per association: 10,009 associations rest on a single source, 6,313 are corroborated by two sources, and the count declines steeply with additional sources; the long tail up to 14 sources marks the best-corroborated associations (e.g., the canonical executors), which carry the strongest multi-database agreement.

**Supplementary Figure S2. Mechanistic associations per source, stacked by RCD entity ontological level.** Each bar shows the number of gene–RCD associations classified as mechanistically involved by the evidence-first adjudication (final_mechanistic = YES), segmented by the ontological level of the corresponding RCD entity: L1 modalities (19 entities), L2 subtypes (7), L3 triggers (2), L4 functional/contextual (13), and L5 generic/umbrella/taxon (12). Colors follow the same concordant palette as Figure 5. The total number of mechanistic associations is printed at the end of each bar, ranging from 1 (Elicit Alert) to 3,872 (Rodrigues de Souza et al., 2025). Source totals are non-additive: a gene–RCD pair with multiple sources contributes to each pertinent source.
