## Supplementary Figure S1 for "RCDome: A Multi-Database Human Gene Atlas of Regulated Cell Death"

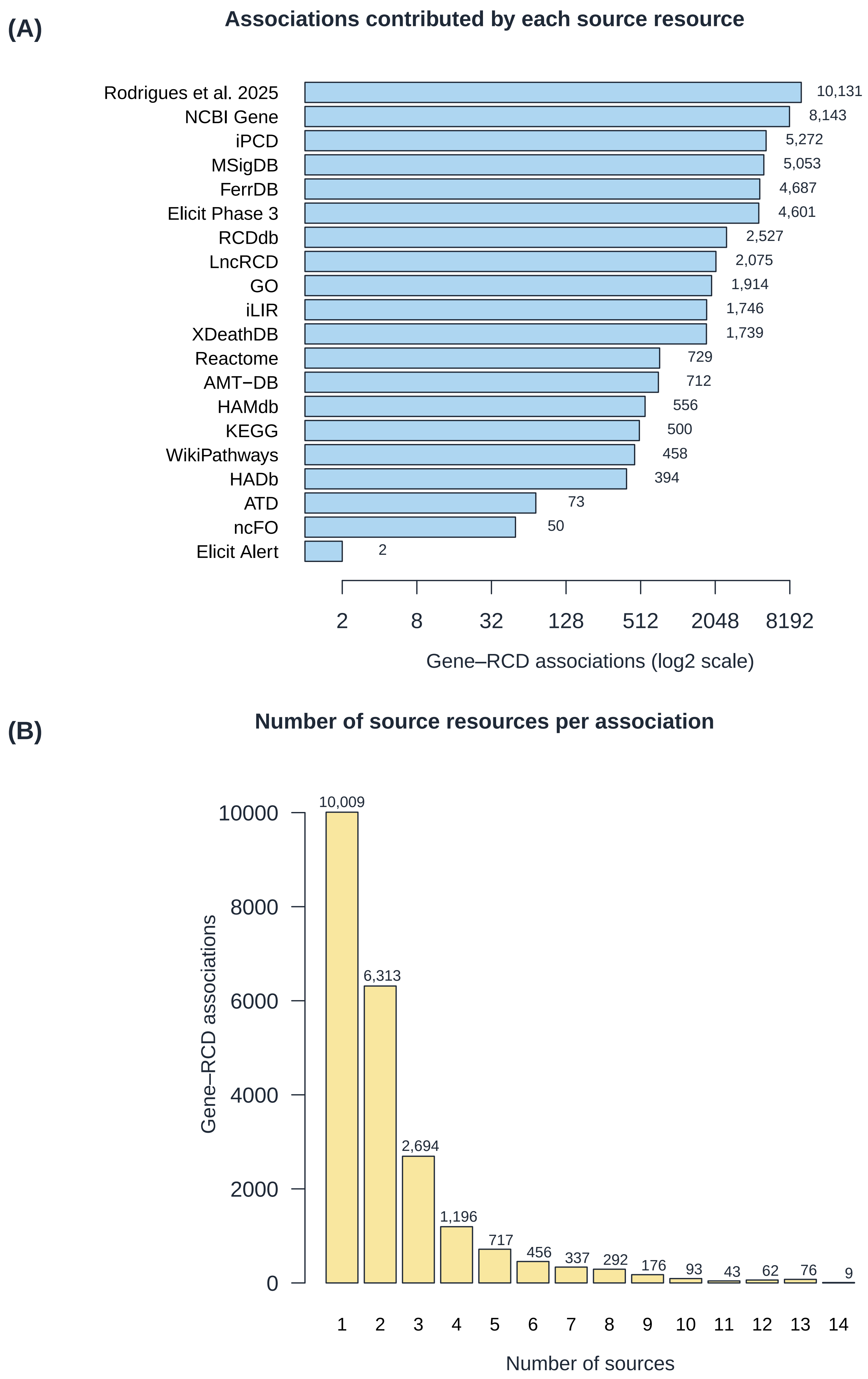

(A)
Associations contributed by each source resource
Rodrigues et al. 2025
NCBI Gene
iPCD
MSigDB
FerrDB
Elicit Phase 3
RCDdb
LncRCD
GO
iLIR
XDeathDB
Reactome
AMT−DB
HAMdb
KEGG
WikiPathways
HADb
ATD
ncFO
Elicit Alert
10,131
8,143
5,272
5,053
4,687
4,601
2,527
2,075
1,914
1,746
1,739
729
712
556
500
458
394
73
50
2
2
8
32
128
512
2048
8192
Gene–RCD associations (log2 scale)
(B)
Number of source resources per association
Gene–RCD associations
0
2000
4000
6000
8000
10000
10,009
6,313
2,694
1,196
717
456
337
292
176
93
43
62
76
9
1
2
3
4
5
6
7
8
9
10
11
12
13
14
Number of sources
