## Supplementary Figure S2 for "RCDome: A Multi-Database Human Gene Atlas of Regulated Cell Death"

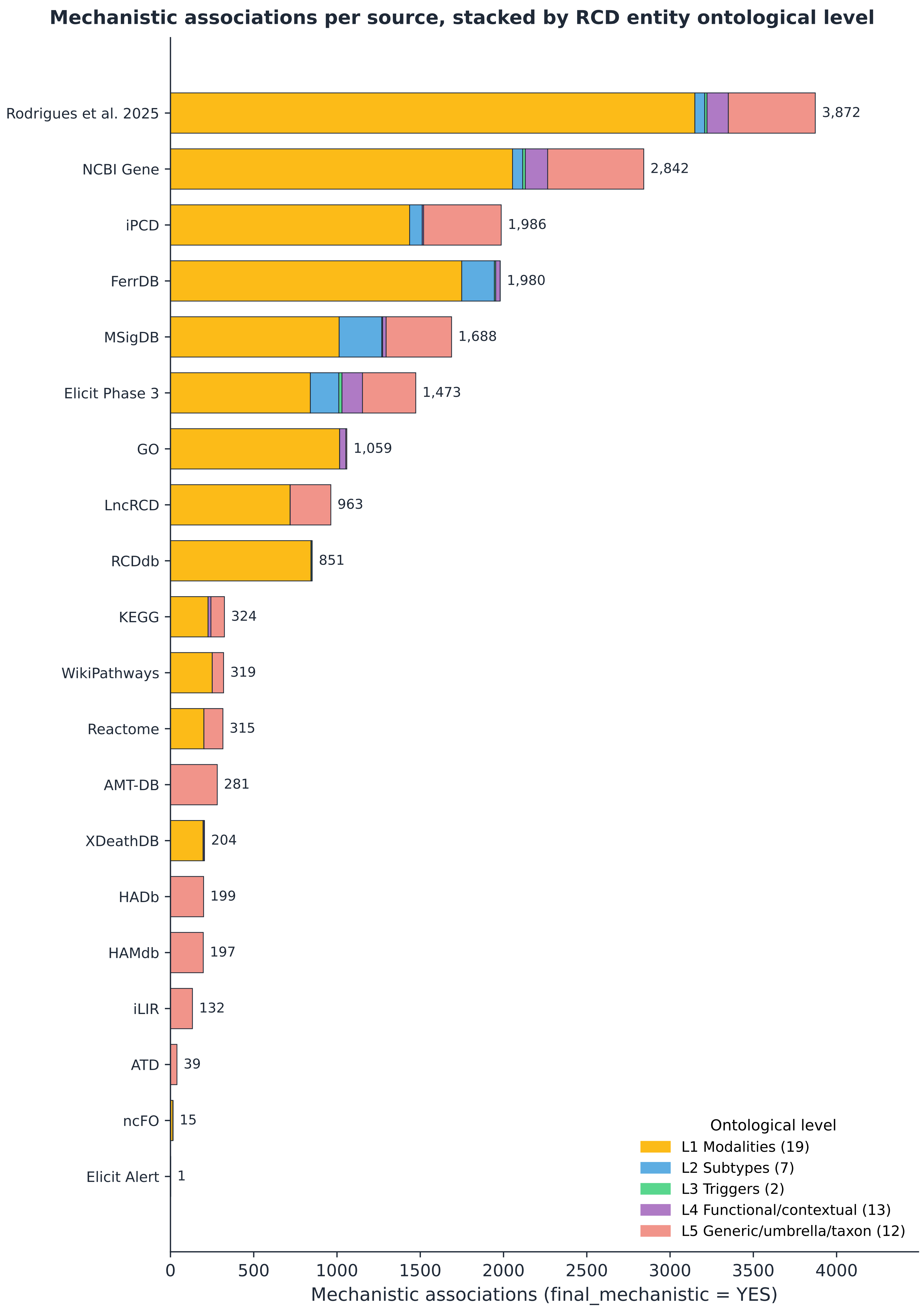

Mechanistic associations per source, stacked by RCD entity ontological level
Rodrigues et al. 2025
NCBI Gene
iPCD
FerrDB
MSigDB
Elicit Phase 3
GO
LncRCD
RCDdb
KEGG
WikiPathways
Reactome
AMT-DB
XDeathDB
HADb
HAMdb
iLIR
ATD
ncFO
Elicit Alert
3,872
2,842
1,986
1,980
1,688
1,473
1,059
963
851
324
319
315
281
204
199
197
132
39
15
1
0
500
1000
1500
2000
2500
3000
3500
4000
Mechanistic associations (final_mechanistic = YES)
Ontological level
L1 Modalities (19)
L2 Subtypes (7)
L3 Triggers (2)
L4 Functional/contextual (13)
L5 Generic/umbrella/taxon (12)
